# DiSCERN - Deep Single Cell Expression ReconstructioN for improved cell clustering and cell subtype and state detection

**DOI:** 10.1101/2022.03.09.483600

**Authors:** Fabian Hausmann, Can Ergen-Behr, Robin Khatri, Mohamed Marouf, Sonja Hänzelmann, Nicola Gagliani, Samuel Huber, Pierre Machart, Stefan Bonn

**Affiliations:** Institute of Medical Systems Biology, University Medical Center Hamburg-Eppendorf, Hamburg, Germany; Center for Biomedical AI, University Medical Center Hamburg-Eppendorf, Hamburg, Germany; Section of Molecular Immunology and Gastroenterology, I. Department of Medicine, University Medical Center Hamburg-Eppendorf, 20246 Hamburg, Germany; Hamburg Center for Translational Immunology (HCTI), University Medical Center Hamburg-Eppendorf, 20246 Hamburg, Germany; Department of General, Visceral and Thoracic Surgery, University Medical Center Hamburg-Eppendorf, 20246 Hamburg, Germany

**Keywords:** Single cell RNA-seq, RNA sequencing, imputation, cell clustering, cell type identification, expression reconstruction, Deep Learning, Machine Learning, auto encoder, batch effect correction, transfer learning, probabilistic modeling, reference atlas mapping, COVID-19, T helper cell, transcription factor analysis, single nuclear RNA-seq, 00-01, 99-00

## Abstract

Single cell sequencing provides detailed insights into biological processes including cell differentiation and identity. While providing deep cell-specific information, the method suffers from technical constraints, most notably a limited number of expressed genes per cell, which leads to suboptimal clustering and cell type identification. Here we present DISCERN, a novel deep generative network that reconstructs missing single cell gene expression using a reference dataset. DISCERN outperforms competing algorithms in expression inference resulting in greatly improved cell clustering, cell type and activity detection, and insights into the cellular regulation of disease. We used DISCERN to detect two unseen COVID-19-associated T cell types, cytotoxic CD4^+^ and CD8^+^ Tc2 T helper cells, with a potential role in adverse disease outcome. We utilized T cell fraction information of patient blood to classify mild or severe COVID-19 with an AUROC of 81% that can serve as a biomarker of disease stage. DISCERN can be easily integrated into existing single cell sequencing workflows and readily adapted to enhance various other biomedical data types.

## 1. Introduction

Single-cell RNA sequencing (scRNA-seq) technologies allow the dissection of gene expression at single-cell resolution, which improves the detection of known and novel cell types and the understanding of cell-specific molecular processes [1, 2]. The extension of the basic scRNA-seq technology with epitope sequencing of cell-surface protein levels (CITE-seq), allows for the simultaneous surveillance of the gene and protein surface expression of a cell [3]. Another recent technological innovation was TCR-seq, which enables the simultaneous sequencing of essential immune cell features and the variable segments of T cell antigen receptors (TCRs) that confer antigen specificity [4, 5].

While several commercial platforms have enabled researchers to use single cell sequencing methods with relative ease and at reasonable cost, the analysis of the high-dimensional scRNA-seq data still remains challenging [6, 7]. The main technical downside of single cell sequencing that impedes downstream analysis is the sparsity of gene expression information and high technical noise. Depending on the platform used, single cell sequencing detects around three thousand genes per cell, giving almost an order of magnitude less genes detected than bulk RNA-sequencing [8]. The term ‘dropout’ refers to genes that are expressed by a cell but cannot be observed in the corresponding scRNA-seq data, a technical artifact that afflicts predominantly lowly to medium expressed genes, as their transcript number is insufficient to reliably capture and amplify them. This missing expression information limits the resolution of downstream analyses, such as cell clustering, differential expression, marker gene and cell type identification [9].

To improve the lack and stochasticity of gene expression information in single cell experiments, several in silico gene imputation methods have been designed based on different principles. Gene imputation infers gene expression in a given cell type or state, based on the information from other biologically similar cells of the same dataset. Several methods utilizing this principle have been developed [10], amongst them DCA, MAGIC, scImpute, DeepImpute and CarDEC [11, 12, 13, 14, 15]. DCA is an autoencoder-based method for denoising and imputation of scRNA-seq data using a zero-inflated negative binomial model of the gene expression. MAGIC uses a nearest neighbor diffusion graph to impute gene expression and scImpute estimates gene expression and drop-out probabilities using linear regression. DeepImpute is an ensemble method, splitting the expression data into multiple pieces and trying to learn imputation of highly correlated genes using deep learning. CarDEC uses a two step procedure of imputation and batch correction using a neural network. All of these algorithms use information from similar cells with measured expression of the same dataset for imputation. Another class of imputation algorithms use bulk RNA-seq data to constrain scRNA-seq expression imputation. Bfimpute [16] uses Bayesian factorization, SCRABBLE [17] matrix regularization, and SIMPLEs [18] a prior distribution on the bulk data to impute scRNA-seq expression. Unfortunately, SCRABBLE and Bfimpute do not scale beyond small single cell datasets and few genes (3000 cells and genes in our hands), and SIMPLEs requires matching single cell and bulk RNA-seq samples, severely constraining their usability.

Similarly, methods (e.g. multigrate[19]) were developed, which use scRNA-seq in combination with complementary, matching data (e.g. CITE-seq, ATAC-seq) to improve imputation. While complementary CITE-seq information is available for many scRNA-seq datasets, other information such as ATAC-seq data of the same sample is usually missing.

While current imputation methods provide improved gene expression information, they still rely on the comparison of similar cells with largely absent gene expression information, for example by using clustering approaches. Genes that are not expressed in neighboring cells cannot be imputed, limiting the value of classical imputation. In an ideal case, it would be possible to obtain information of the expected true gene expression per cell, or at least expression information with less technical noise, to reconstruct the true expression at single cell level. Additionally, recent studies question the number of technical dropouts in UMI-based sequencing technologies [20, 21] and thus challenge classical imputation based methods. However, there are still batch specific changes, e.g. capture rate of specific genes and differences in sample processing, which affect the single cell data, beyond dropout. These changes can be wanted (enforced by the experimental setup) or unwanted (stochastic changes in the experimental setup, material).

Recent work has shown the effectiveness of deep generative models (e.g. Autoencoders and Generative Adversarial Networks) to infer realistic scRNA-seq data and augment scarce cell populations using Generative Adversarial Networks [22] or the prediction of perturbation response using Autoencoders [23]. We hypothesized that a deep generative model could allow for the reconstruction of missing single cell gene expression information (low quality - lq) by using related data with more genes expressed (high-quality - hq) as a reference, a reference-based approach to gene expression inference (Figure 1A). In other words, lq data with many missing gene expression values and bad clustering could be transformed into data with few missing genes and improved clustering if the “style” of a related hq dataset could be transferred to it. In the best case, it would be possible to infer gene expression information for single cell data (lq) by using purified bulk RNA-seq data (hq), obtaining over ten thousand genes expressed per cell. We envision that this approach, when properly calibrated, gains deep mechanistic insights into data beyond what is currently measurable. It is important to note that the concept of using hq data to reconstruct gene expression in lq data is different from classical imputation algorithms that infer gene expression based on nearby cells from the same dataset, as outlined above.

**Figure 1:**
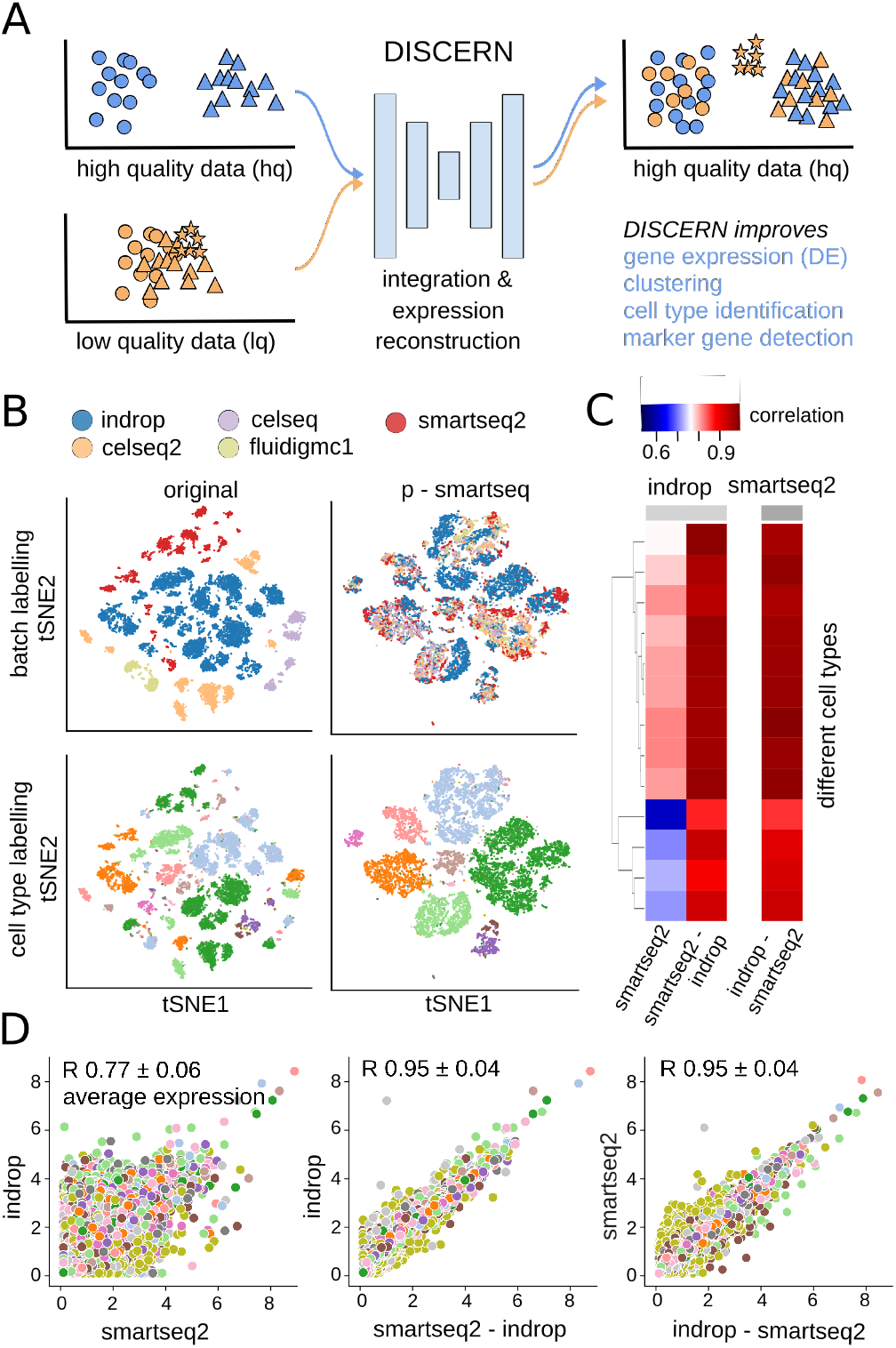
Integration and expression reconstruction of single cell sequencing data. **A**: DISCERN transfers the style of a high-quality (hq) dataset to a related low quality (lq) dataset, enabling gene expression reconstruction that results in improved clustering, cell type identification, marker gene detection, and mechanistic insights into cell function. The hq and lq datasets have to be related but not identical, containing for example several overlapping cell types but also exclusive cell types of cell activity states for one or the other dataset. **B**: t-SNE visualization of the pancreas dataset before reconstruction (original) and after transferring the style of the smartseq2 dataset using DISCERN (p-smartseq2). The upper row shows the dataset of origin before and after projection colored by batch and the lower row colored by cell type annotation (details of 13 cell types in supplements). **C** and **D**: Average gene expression (over all the cells of a given type) of the pancreas indrop and smartseq2 datasets before (first column and panel) and after smartseq2 to indrop (second column and panel), and after indrop to smartseq2 projection (third column and panel). **C**: Gene correlation by cell type shown in colored heatmap. **D**: Each colored point represents a single gene colored by the cell type. The mean Pearson correlation with one standard deviation over all cell types is shown in the figure title.

Based on the above considerations, we developed DISCERN, a novel deep generative neural network for directed single cell expression reconstruction. DISCERN allows for the realistic reconstruction of gene expression information by transferring the style of hq data onto lq data, in latent and gene space. Our experiments on real and simulated data show that DISCERN outperforms several existing algorithms in gene expression inference across a wide array of single cell datasets and technologies, improving cell clustering, cell type and activity detection, and pathway and gene regulation identification. To obtain deep insights into the cellular changes underlying COVID-19, we reconstructed single cell expression data of patient blood and lung immune data. While in our initial analysis [24] of blood data we detected few immune cell types, expression reconstruction with DISCERN resulted in the detection of 28 cell types and states in blood, including two unseen disease-associated T cell types, cytotoxic CD4^+^ and CD8^+^ Tc2 T helper cells. Reconstructing a second COVID-19 blood dataset with disease severity information, we were able to classify mild and severe COVID-19 with an AUROC of 81%, obtaining a potential biomarker of disease stage. DISCERN can be easily integrated into existing workflows, as an additional step after count mapping. Given that DISCERN is not limited by a predefined distribution of data, we believe that it can be readily adapted to enhance various other biomedical data types, especially other omics data such as proteomics and spatial transcriptomics.

## 2. Results

### 2.1. The DISCERN algorithm for directed expression reconstruction

We aim to realistically reconstruct gene expression in scRNA-seq data by using a related hq dataset. Ideally, this expression reconstruction algorithm should meet several requirements [7]. First, it needs to be **precise** and model gene expression values realistically. It shouldn’t remove information of cellular identity to form ‘average cells’ or collapse different cell types or states into one. Second, the network should be **robust** to the presence of different cell types in hq and lq data, or an imbalance in their relative ratios. It shouldn’t, for instance, ‘hallucinate’ hq-specific cells into the lq data. Lastly, the network should be directional, as the user should be able to choose the target (reference) dataset.

With these prerequisites in mind, we designed a deep neural network for directed single cell expression reconstruction (DISCERN) (Figure S1B) that is based on a modified Wasserstein Autoencoder [25]. A unique feature of DISCERN is that it transfers the “style” of hq onto lq data to reconstruct missing gene expression, which sets it apart from other batch correction methods such as [26], which operate in a lower dimensional representation of the data (e.g. PCA, CCA). To allow DISCERN to accurately reconstruct single cell RNA-seq expression based on reference data, the structure of the network had to be adapted in several ways. First, we implemented Conditional Layer Normalization (CLN) [27, 28, 22] to allow for directed expression reconstruction of lq data based on reference hq data (Figure S1B & S2). Second, we added two decoder heads to the network to enable it to model dataset-specific dropout rates and gene expression separately. Lastly, we extended DISCERN’s loss function with a binary cross-entropy term for learning the probability of dropouts to increase general inference fidelity. Further algorithmic details of DISCERN can be found in the methods and Figure S1.

We first demonstrate DISCERN’s capabilities to faithfully reconstruct gene expression using five pancreas single cell expression datasets from 5 different studies [29, 30, 31, 32, 33], with varying quality (Tables S1 and S2). The pancreas data is widely used for benchmarking and it is ideal to evaluate expression reconstruction for many cell types and sequencing technologies. We consider a dataset as hq when the average number of genes detected per cell (GDC) (e.g. smartseq2, GDC 6214) is much higher than in a comparable lq dataset (Table S2). Conversely, a dataset is lq when the average cell has lower counts and fewer genes expressed than a comparable hq dataset (e.g. indrop, GDC 1887). Throughout this text, we will name sequencing technologies with capital (e.g. Smart-Seq2, InDrop) and datasets with lower case first letters (smartseq2, in-drop). We trained DISCERN on these five pancreatic single cell datasets and assessed the integration of data in gene space and the expression reconstruction per cell type. While uncorrected data cluster by batch and not by cell type, DISCERN-integrated data show good batch mixing and clustering of cells by cell type across all five datasets (Figure 1B & Figure S2). To get a clearer picture of DISCERN’s expression reconstruction capabilities we next calculated correlation coefficients of measured expression between the lowest quality inDrop and highest quality Smart-Seq2 data, before and after expression reconstruction using DISCERN. The mean expression reconstruction of indrop-lq to smartseq2-hq and smartseq2-hq to indrop-lq data is very accurate, showing a Pearson correlation of *r* = 0.95, while mean expression correlation between uncorrected indrop-lq and smartseq2-hq data is only *r* = 0.77 due to strong batch effects (Figure 1C & D, Figures S3 and S4). The improved quality of indrop-lq data reconstructed to smartseq2-hq level is validated by the strong increase of genes expressed per cell, ranging from ≈2000 genes per cell in the uncorrected indrop-lq data to ≈6000 genes in the indrop-lq data after reconstruction (Figure S5).

We next investigated the effect of reconstruction of three cell type-specific genes, before and after correction across the five pancreas datasets (Figure S6). Insulin expression in the pancreas should be largely restricted to beta cells [34], which can be observed in the uncorrected smartseq2-hq and celseq2 datasets, while the indrop-lq batch shows a diffuse pattern of insulin expression across cell types (Figure S6A left panel). This diffuse insulin expression is corrected by reconstructing the smartseq2-hq expression pattern from the indrop-lq data (Figure S6A middle panel). In general, the expected specificity of insulin expression in beta cells can be recovered for all datasets when using DISCERN’s reconstruction using the smartseq2-hq reference. Conversely, the reconstruction from hq to the indrop-lq reference results in diffuse insulin expression across all reconstructed datasets (Figure S6A right panel). We obtained similar results for the pancreatic acinar cell-specific gene REG1A and the delta cell-specific gene SST, both of which show diffuse expression across cell types in the uncorrected inDrop data and cell-specific expression after reconstruction using smartseq2-hq reference (Figure S6B & C). Interestingly, DISCERN can not only recover biological expression information, but it is also able to apply sequencing method-specific effects after reconstruction. The smartseq2-hq dataset, for instance, displays nearly no ribosomal protein coding gene expression after sequencing as previously reported by [8], while data sequenced using InDrop, Cel-Seq, or Cel-Seq2 shows prominent ribosomal protein coding gene expression (Figure S6D, left panel). When reconstructing smartseq2-hq data to indrop-lq data, ribosomal protein coding gene expression is re-instantiated (Figure S6D, right panel).

We further corroborated DISCERN’s capability to integrate and reconstruct gene expression in the more complex difftec dataset (Tables S1 and S2), consisting of 14 single cell peripheral blood mononuclear cell (PBMC) datasets across a wide range of technologies. Similar to pancreas, the difftec dataset is widely used for benchmarking and it is ideal to evaluate expression reconstruction for even more cell types and sequencing technologies. The different single cell technologies show large variation in quality, with an GDC ranging from 422 in Seq-Well to 2795 in Smart-seq2. We trained DISCERN on these 14 PBMC single cell datasets and observed very good integration in gene space (Figure S7). We then reconstructed chromium-v2-lq (GDC 795) using a chromium-v3-hq reference (GDC 1514) and observed high mean gene expression correlation between the reconstructed and reference datasets (Figures S8 and S9). These results across 19 single cell datasets provide first evidence for the high-quality data integration and expression reconstruction that can be obtained with DISCERN.

### 2.2. Specific and robust gene expression inference

We next investigated the precision and robustness of DISCERN’s expression reconstruction in more detail and compared DISCERN’s performance to several state-of-the-art algorithms for expression imputation and data integration.

We explored the robustness of DISCERN to the choice of its hyperparameter by testing various non-default combinations of the four hyperparameters influencing the model training. In all combinations DISCERN was able to achieve a pearson correlation of > 0.94 and a correlation of 0.95 with the default parameter when reconstructing the indrop-lq batch to the smartseq-hq batch of the pancreas dataset (Figure S10). This provides strong evidence that DISCERN’s performance is robust to the choice of hyperparameters.

Since expression reconstruction can be seen as a generalization of expression imputation, we compared DISCERN to DCA, MAGIC, and scImpute, CarDEC, and DeepImpute, five state-of-the-art imputation algorithms [11, 12, 13, 14, 15]. Expression reconstruction can also be viewed as a batch correction task in gene space, which is why we additionally compared DISCERN to scGEN, Seurat, trVAE and scVI [23, 26, 35, 36]. It is important to note, however, that these batch correction methods were not designed for the expression reconstruction task and use a lower dimensional representation to align different batches. Seurat uses canonical correlation analysis and scGEN uses the bottleneck layer representation of an autoencoder to calculate and apply linear transformations. trVAE and scVI explicitly encode the conditional information in the autoencoder architecture.

We compared the ability of these models to adjust expression information on the pancreas dataset by reconstructing the indrop-lq expression based on the smartseq2-hq expression. Generally deep learning methods, which allow for projection (scGEN, scVI, trVAE, DISCERN), show the best performance, with DISCERN showing the lowest deviation between cell types (Figure S11). We also investigated the gene expression standard deviation on the same data, showing that DISCERN reconstructs the variation in the indrop-lq best, with scVI showing only slightly worse performance (Figure S12). A factor which has a high impact on the variation is the number of dropouts found in each gene. While most imputation methods try to remove them, we think they contain useful information as well [37]. DISCERN is able to capture the batch-specific dropout rate much better compared to other batch correction or imputation methods (Figure S13). Interestingly deep learning methods, scVI, scGEN, DeepImpute and DCA for example, achieve a similar correlation of the dropout rate than classical methods, for example Seurat and MAGIC, even if deep learning methods seem to be better in reconstruction of mean expression (Figure S11). It is important to highlight that the proper estimation of expression variation and the dropout rate is pivotal for the reliable computation of differentially expressed genes. Since DISCERN displays the best variance estimation, it also achieves the best median correlation of the differentially expressed genes (Figure S14).

To investigate the precision of gene expression reconstruction, we created an artificial dataset by dividing the smartseq2-hq pancreas data into two batches, smartseq-lq and smartseq2-hq. In the smartseq-lq batch, the top one KEGG pathways per cell type were removed by setting the expression of genes contained in these pathways to zero, while the smartseq2-hq remained unaltered. Therefore, a reconstruction of smartseq-lq data using smartseq2-hq reference (reconstructed-hq) should ideally recover the smartseq-lq expression to its original state, prior to the removal of the genes. DISCERN is able to reconstruct the mean expression for all cell types, achieving a correlation *r* = 0.99 (Figure 2A). DCA (*r* = 0.66), MAGIC (*r* = 0.34), scImpute (*r* = 0.80), Deep-Impute *r* = 0.89 and Seurat (*r* = 0.76) have significantly lower correlation between the smartseq2-hq and reconstructed-hq gene expression (Figure 2A). scGen (*r* = 0.98), scVI (*r* = 0.99) and trVAE (*r* = 0.99) show similar performance compared to DISCERN. Moreover, scGEN and trVAE however perform worse in reconstruction of highly expressed genes, while scVI slightly overestimates the expression in general (Figure 2A). We obtained similar results on the difftec dataset, with DISCERN (*r* = 0.98) outperforming DCA (*r* = 0.47), MAGIC (*r* = 0.21), scImpute (*r* = 0.04), Seurat (*r* = 0.92), scVI (*r* = 0.96), trVAE (*r* = 0.95), DeepImpute (*r* = 0.58), and scGEN (*r* = 0.94) (Figure S15). To further investigate gene expression reconstruction specificity, we compared the correlation of reconstructed-hq to smartseq2-hq data after performing differential gene expression (DEG) for each cell type against all other cell types (Figure 2B, upper panel). DISCERN is able to recover the correct DEG t-statistics with a median correlation of 0.92, improving over state-of-the-art tools by more than 6 percentage points. In the corresponding experiment using the difftec dataset, DISCERN achieves a median correlation of 0.86, which is a 21 percentage point improvement over competing methods (Figure S16).

**Figure 2:**
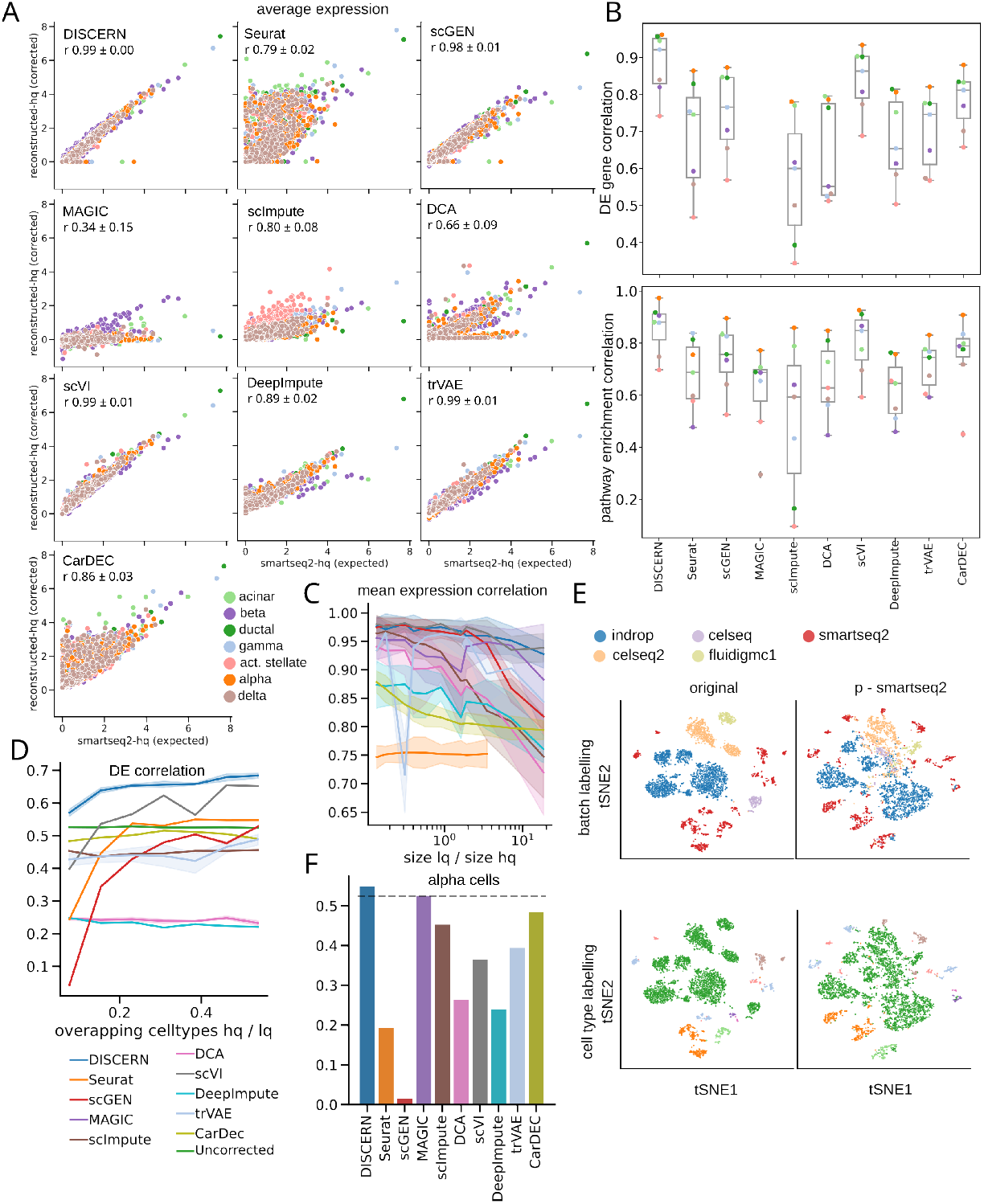
Expression reconstruction benchmark of DISCERN and five state-of-the-art batch correction and imputation algorithms. Comparison of the expression reconstruction performance of Seurat, scGEN, MAGIC, scImpute, DCA, scVI, trVAE, DeepImpute, and DISCERN using smartseq2 data. The smartseq2 data was split into a smartseq2-lq and a smartseq2-hq batch. The smartseq2-lq batch was modified such that the expression of all genes of a cell type determining pathway (top ranked by GSEA) was set to zero. The expression of the in silico altered pathway genes was then compared between reconstructed-hq data and the unaltered smartseq2-hq data. **B**: Differential gene expression and pathway enrichment correlation of the reconstructed-hq to the expected values before removal. The smartseq2-lq data was the same as in **A**. The DEG analysis was restricted to genes which were removed in the smartseq2-lq batch. Correlation of the DEG analysis was based on the t-statistic and for the pathway enrichment analysis on the normalized enrichment scores. **C**: Mean expression correlation of reconstructed-hq with the expected expression in smartseq-hq data for different ratios of lq to hq data. The standard deviation indicates the deviation in correlation of the cell types. The datasets were created as described in **A**. **D**: Alpha cells were removed from the smartseq-hq batch and left in the low quality batches. The number of overlapping cell types between the hq and lq data was then altered by removing cell types, which overlap between lq and hq data, from the lq data before preprocessing and expression reconstruction. The ratio of the intersection size to to the total number of cell types is shown on the x-axis. The y-axis shows the correlation of the t-statistics of alpha cells from lq-batches vs other cells from the smartseq2 batch with ground truth alpha cells from the smartseq2 batch vs other cells from the uncorrected smartseq2 batch. We used Spearman rank correlation for the comparison, since no gene subset was used. **E**: t-SNE visualization of the cell type removal experiment where alpha cells are removed from the smartseq2 batch and all non-alpha cells are removed from the lq-batches, such that there is no overlap between lq and hq. **F**: Spearman correlation of the t-statistics of alpha cells from lq-batches vs other cells from the smartseq2 batch with ground truth alpha cells from the smartseq2 batch vs other cells from the uncorrected smartseq2 batch. The dataset was the same as in **E** (no cell type overlap between hq and lq data). The dotted line indicates the correlation achieved without reconstruction.

Since the genes were initially selected using KEGG gene set enrichment analysis, the reconstruction of the corresponding pathways was investigated by performing KEGG gene set enrichment analysis on the DEG results. DISCERN is able to recover the pathway expression enrichment scores with a median correlation of 0.88, exceeding the performance of scVI by more than3 percentage points on median (Figure 2B, lower panel). In the corresponding experiment using the difftec dataset, DISCERN achieves a median correlation of 0.77, out-performing Seurat and scGen by more than 16 percentage points (Figure S17).

While DISCERN outperforms competing algorithms in expression and pathway reconstruction correlation, it achieves the fourth-best correlation for the DEG fold-change (FC) of reconstructed-hq to smartseq2-hq data for the pancreas (Figure S18) and reconstructed-hq to chromium-v3-hq difftec datasets (Figure S19). In both cases Seurat, scVI and CarDEC achieve better correlation, which is due to the fact that DISCERN slightly underestimates FC in favor of superior DEG variance estimation.

Next, we show DISCERN’s expression reconstruction robustness with respect to varying sizes of lq to hq data. It is conceivable to assume that a large amount of hq data would benefit the expression reconstruction of the lq data, which makes it important to understand at what ratio good results can be expected. Interestingly, DISCERN seems to be very robust across a wide range of smartseq2-lq to smartseq2-hq ratios, with correlations of 0.98 (ratio of lq/hq 0.14) to 0.93 (ratio of lq/hq 18.4), while the second-best performing algorithm scGen showed a 11 percentage point decrease in performance (0.82 for ratio of lq/hq 18.4) (Figure 2C, Figure S20). We observed similar results for the correlation of t-statistics, showing a slight dependence of DISCERN’s performance on the lq/hq ratio (Figure S21). In general, all methods show better performance with a small ratio of lq/hq data, while DISCERN and scVI shows least dependence and outperform other algorithms in the correlation of expression and t-statistics, especially in the case of high lq/hq ratio.

Another aspect of expression reconstruction robustness is the dependence of the algorithm on the cell type or cell state similarity of the lq and hq datasets. In the optimal case, DISCERN would not require that the lq and hq datasets have overlapping cell types to perform an accurate expression reconstruction, which is theoretically possible if the network learns the general gene-regulatory expression logic of the hq data (see discussion). To understand the dependence on dataset similarity, we removed a complete cell type, pancreas alpha cells, from the smartseq2-hq data and left the alpha cells in the smartseq2-lq data. We then additionally varied the number of common cells in the lq and hq data, starting with no overlapping cells (only alpha cells in the lq and all cells except alpha in the hq data) and ending with almost complete overlap (all cells overlap between the smartseq2-hq and -lq data, except for the alpha cells only present in lq data) (Figure 2D). When evaluating DEG correlation, DISCERN was the only method consistently achieving better performance than uncorrected data, outperforming scVI by 2 to 17 percentage points (Figure 2D). Similarly, DISCERN was consistently achieving better performance than uncorrected data in the FC correlation task (Figure S22).

We next took a closer look at the integration and expression reconstruction performance when no cell types overlap between the lq (alpha cells only) and hq (all other cells) data. Notably, Seurat seems to over-integrate cell types, mixing smartseq2-hq beta and gamma cells with reconstructed-hq alpha cells from other batches (Figure S23), while all other methods keep the smartseq2-hq and reconstructed-hq exclusive cell types separate (Figure 2E & Figure S23). This over-integration seems to be causal for Seurat’s poor DEG correlation performance (*r* = 0.19), while DISCERN (*r* = 0.55) is the only method achieving better performance than uncorrected cells (*r* = 0.52) (Figure 2F). Thus, DISCERN is able to keep existing expression correlations and improves the detection of cell type specific genes by reconstruction using an hq batch as reference. In conclusion, DISCERN is both a precise and robust method for expression reconstruction that outperforms existing methods by a significant margin.

### 2.3. Improving cell cluster, type, and trajectory identification

The comparison to competing methods provided evidence for DISCERN’s superior expression reconstruction. Now, we will delineate how DISCERN’s expression reconstruction improves downstream cell clustering, cell type and activity state identification, marker gene determination, and gene regulatory network and cell trajectory analysis.

Batch correction algorithms are usually evaluated by comparing their ability to integrate cells coming from the same cell type but different batches, using the silhouette score, the adjusted rand index (ARI), and adjusted mutual information (AMI). DISCERN often outperforms all competing methods across all metrics, achieving state-of-the-art performance in batch mixing and cell type clustering (Figures S24 to S26).

To understand if cell-determining gene expression and pathways could be recovered with expression reconstruction, we used a single nuclear sequencing (sn-lq) and scRNA-seq (sc-hq) data pair that was prepared from the same liver metastasis biopsy [38]. We reconstructed sn-lq data using the sc-hq reference, obtaining reconstructed-hq data. While single nuclear sequencing provides reduced expression information in the average counts per cell as compared to scRNA-seq (Table S2) [38], it is still the method of choice to obtain cell-specific expression information when intact single cells cannot be recovered from a tissue (e.g. after tissue fixation or freezing). It is important to note that nuclear transcripts reflect current gene activity, which in part might not correlate with transcripts that have lifetimes of up to days. Before integration, the sn-lq and sc-hq datasets cluster by batch and not by cell type, while after expression reconstruction with DISCERN cells cluster by type and not by batch (Figure S27). This is reflected in an expression correlation of 0.49 (sc-hq vs. sn-lq) before and 0.97 after reconstruction (sc-hq vs. reconstructed-hq) (Figure S28). DISCERN reconstruction resulted in the expression of T cell receptor signaling genes in reconstructed T cells (Figure S29) and antigen presentation genes in macrophages (Figure S30), providing evidence that DISCERN faithfully recreates cell-determining genes and pathways based on the hq data. Seurat, CarDEC and scImpute are not able to reconstruct the expression information and show a similar expression pattern as the uncorrected sn-lq dataset. In their reconstructions (seurat-hq, CarDEC–hq, scImpute-hq and sn-lq) the expression of important T cell marker genes such as *CD3E, CD3D* and *CD8A* is largely absent, while in sc-hq and DISCERN-hq the expression is easily detectable (Figure S29). DCA, scVI,scGEN, MAGIC and trVAE show a strongly disturbed expression pattern, where many genes show a much larger expression than in the sc-hq or the sn-lq datasets (Figures S29 and S30).

To further corroborate the advantage of single nuclear expression reconstruction, we next aimed to increase the T cell subtype resolution of human single nucleus acute kidney injury data (kidney-lq) by using matching single cell data (kidney-hq). Only 1% of kidney-lq nuclei show *CD3D, CD3E* or *CD3G* expression, compared to 7% of the cells in the kidney-hq dataset. Seurat and DISCERN were able to detect T cells in the reconstructed kidney-lq (reconstructed-hq) and the kidney-hq data with notable *CD3D* expression in this cluster (Figure S31). The reconstructed-hq and the kidney-hq T cells were further classified into T cell subtypes and activation states (Figure S31C). While a large proportion of T cells detected in Seurat reconstructed data could not be annotated due to missing *CD3D, CD4,* and *CD8A* expression, DISCERN reconstructed data does not present these limitations.

It is intriguing to observe that many marker genes are still hard to detect in kidney single cell RNA-seq data but also in the antigen presentation pathway in macrophages (Figure S30). This is most probably due to dropout. Thus, we rationalized that bulk RNA sequencing (RNA-seq) data of purified cell types (e.g. FACS sorted immune cells) is a suitable hq proxy for the expected gene expression per cell. RNA-seq data of purified cells is readily available from public repositories, making it possible to obtain thousands of purified immune cell RNA-seq samples (see methods). We therefore set out to increase cluster, cell type, gene regulatory network, and trajectory identification of scRNA-seq data by reconstructing gene expression using a related RNA-seq reference (Figure S32). For the scRNA-seq data we chose a cord blood mononuclear cite-seq dataset (cite-lq) that was labeled with 15 antibodies (Table S3) to allow for surface protein-based cell type discovery [39]. The CITE-seq information allowed us to confirm expression reconstruction by DISCERN in cases where gene expression is absent but protein expression and cell identity are validated via antibody labeling. For the RNA-seq data, we selected 9852 purified immune samples (bulk-hq) and proceeded to reconstruct cite-lq (GDC 798) using a bulk-hq (GDC 13 104) reference to obtain reconstructed-hq data with DISCERN. We first investigated the correspondence of gene expression prior (cite-lq) and post reconstruction (bulk-hq) with antibody-based surface protein labeling of *CD3D*, *CD4, CD8A, CD2, B3GAT1, FCGR3A, CD14, ITGAX* and *CD19* (Figure 3A, Figure S33). For several proteins (CD8A, B3GAT1, CD4), the corresponding cite-lq gene expression was absent and cell type-specifically re-instantiated in the reconstructed-hq expression data with DISCERN (Figure 3A, Figure S33).

**Figure 3:**
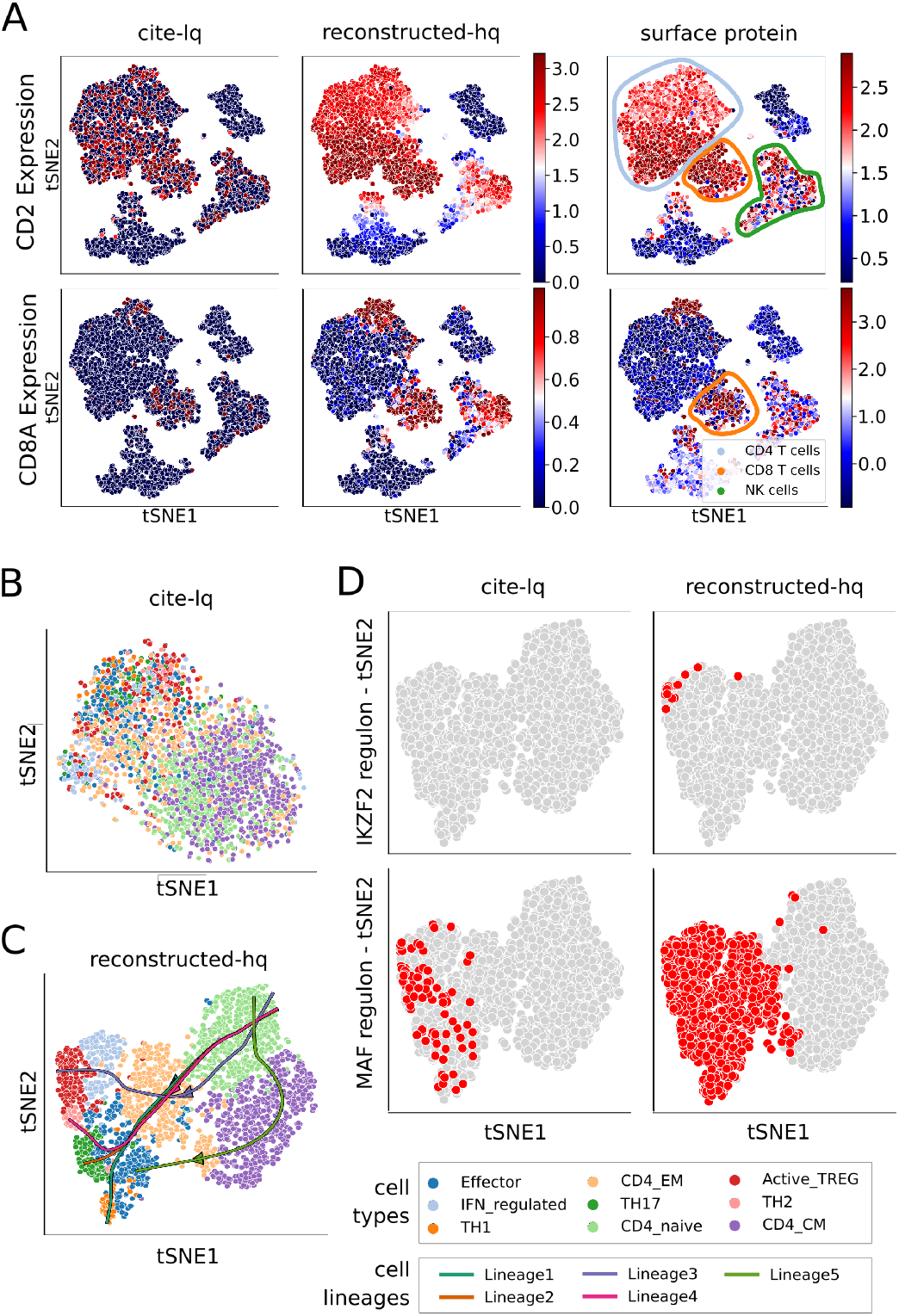
Expression reconstruction improves downstream analyses including cell identification, gene regulation, and trajectory inference. The cite-lq dataset was reconstructed using bulk-hq data and compared to ground truth CITE-seq (surface protein) information. The CITE-seq information was not used during training of DISCERN. **A**: t-SNE visualization of *CD2* (first row) and *CD8A* (second row) gene (first two columns) and protein (last column)) expression. The first column depicts gene expression for uncorrected cite-lq, the second for reconstructed-hq, and the third protein surface expression ground truth information. Cell types commonly known to express these genes are highlighted with colored circles in the last column. **B**: t-SNE visualization of CD4^+^ T cells in the cite-lq dataset. Cell types were assigned using louvain clustering on the reconstructed-hq data (see C) and show no clear clustering. **C**: t-SNE and trajectory information of CD4^+^ T cell subtypes found by Slingshot analysis on reconstructed-hq data. While uncorrected data shows no clear cell type clustering (see **B**), reconstructed data shows a clear grouping of cell types. Trajectories were calculated using CD4_naive as starting point and TH2, TH17, TH1, Active_TREG, CD4_CM as end-points. Lineage1 indicates TH1, Lineage2 TH17, Lineage3 Active_TREG, Lineage4 TH2, and Lineage5 Effector cell differentiation. **D**: Detection of regulons that are specific for CD4^+^ T cell subtypes using pySCENIC. The first column shows regulons found in the uncorrected cite-lq and the second column in reconstructed-hq data.

In cases where cell type-specific gene and protein expression matched cite-lq data (*CD3D, CD14*) the expression in reconstructed-hq data was left unaltered (Figure S33). In some instances, we observed low cell type-specific expression in the cite-lq data (*CD8A, CD2, FCGR3A, CD19*) that matched protein expression (Figure S33). In these cases, gene expression was increased in the correct cell types in the reconstructed-hq data. In general, we observed increased agreement between cell type-specific surface protein and gene expression after reconstruction, showing that DISCERN doesn’t invent or ‘hallucinate’ cell types but reconstructs the expected expression specific for each cell type. We further corroborated these results by selecting eight known cell type-specific cytosolic proteins and investigated their expression before and after expression reconstruction. *MS4A1* (B cells), *IL7R* (CD4^+^ T cells), *MS4A7* (Monocytes), *GNLY* and *NKG7* (NK cells) showed consistent expression before and after reconstruction (Figure S34). The chemokine receptors *CCR2* (Monocytes, activated T cells), *CXCR1* (NK cells), and *CXCR6* (CD8^+^ T cells) showed the correct cell type-specific expression only after expression reconstruction (Figure S34) [40]. It is notoriously hard to obtain cell subtype-specific information from blood mononuclear scRNA-seq data, especially for CD4^+^ T helper cells due to their limited activation status in healthy individuals. This doesn’t mean that polarized CD4^+^ T helper cells do not exist in healthy blood, as they are commonly detected after stimulation using FACS (Table S3) [41]. This lack of resolution in scRNA-seq impedes clustering, marker gene, and trajectory analyses, a drawback that could be overcome using DISCERN’s expression reconstruction. We therefore compared CD4^+^ T cell (gene expression of *CD4* > 1 and *CD3E* > 2.5) clustering and subtype identification using cite-lq and reconstructed-hq data. While clustering with the leiden algorithm [42] using highly variable genes of cite-lq data resulted in an unstructured distribution of CD4^+^ T cell subtypes (Figure 3B), clustering of reconstructed-hq data yields detailed insights into T helper cell subtypes of blood mononuclear data (Figure 3C). After reconstruction, we were able to characterize TH17, TH2, TH1, HLA-DR expressing TREG (Active_TREG), naive CD4^+^ T cells (CD4_naive), effector-memory CD4^+^ T cells (CD4_EM), central-memory CD4^+^ T cells (CD4_CM), and effector cells expressing IFN-regulated genes (IFN_regulated) (Figure 3C). We selected published cell-determining marker genes and observed that many of them were dropped out in the uncorrected data but were present after reconstruction (Figure S35). The absence of marker genes in uncorrected data results in poor clustering and cell type identification, while single positive cells are detectable in the respective neighborhood identified by reconstructed counts (Figure S35). Importantly, we observed that in all cases the DISCERN-estimated proportions of T helper subsets fall within the range of expected proportions as assessed by previous FACS studies (Table S3, Figure S36). These findings are important, as they prove once more that DISCERN discovers the correct cell subtypes and cell proportions, in this case substantially outperforming the available CITE-seq information in cell subtype resolution.

To further verify the cell type annotations, we extracted the top cluster-determining genes from the reconstructed-hq data. Members of the TNF-receptor superfamily are known to be expressed in T helper cell subtypes [43], which can be observed after reconstruction in TH17 cells and partially in TH1, TH2, Active_TREG and IFN_regulated cells (Figure S37). Similarly, reconstructed TH1 cells show the expected high expression of granzymes *GZMK* and *GZMA* [44], while *MIAT* and *HLA* expression are found in activated TREG cells after reconstruction (Active_TREG cluster, Figure S37) [45, 46]. *NOG* expression is detected in reconstructed CD4_naive cells, as previously described [47]. In addition, reconstructed CD4_naive, CD4_EM and CD4_CM show low expression of the genes important for the T helper subtypes TH1, TH2, TH17, Active_TREG and IFN_regulated. We further corroborated our cell type annotation of reconstructed-hq data by observing the expected expression of several established T cell subtype markers (Figure S38). We compared these newly found clusters to representations found with Seurat, multigrate, and in uncorrected cite-lq data. The uncorrected cite-lq data manifests cluster separation for some cell types, most notably IFN_regulated and Active_TREG cells (Figure S39A). Seurat reconstruction and multigrate imputation with CITE-seq information results in the mixing of cell types and clusters (Figure S39B & C). A further comparison to Bfimpute and SCRABBLE was impossible due to the dataset size, as outlined in the introduction.

Similar to improved clustering and cell subtype detection, DISCERN reconstructed-hq data resulted in improved gene regulatory network inference with SCENIC [48]. SCENIC infers transcription factor-regulated gene expression modules of single cell data. While cite-lq data resulted in a scattered distribution of transcription factor networks across several T helper cell subtypes, SCENIC with reconstructed-hq data showed transcription factor regulation in the correct subtypes (Figure 3D). After expression reconstruction the IKZF2 regulon is detected in activated TREG cells [49] and the MAF regulon is found in differentiated CD4^+^ T cells but not in naive CD4^+^ T cells [50]. A weak signal of the MAF regulon is already detectable in the cite-lq data, yet strongly increased in reconstructed-lq, while maintaining differentiated T helper cell specificity (Figure 3D). Furthermore, after reconstruction with DISCERN we could identify the TH17 associated master transcriptional regulators RORC(+) and RORA(+) [51], which were scattered over all TH17 cells before reconstruction (Figure S40). Seurat is able to partially reconstruct the expression of the RORC(+) regulon but fails to detect the more specific RORA(+) expression (Figure S40).

Finally, we wanted to investigate if DISCERN could also enhance cell trajectory analyses with Slingshot of the citeseq data [52]. We focused on the differentiation of effector and other T helper cell subtypes and found five lineages that either pass through or terminate in the effector cell cluster in reconstructed-hq data (Figure 3C). Two trajectories were of special interest to us: Lineage1 from CD4_naive to TH1 cells (Figure S41) and Lineage2 from CD4_naive to TH17 cells (Figure S42). While the expression change along the trajectory in uncorrected data (Figure S41A, Figure S42A) is hardly visible, cell type-specific clusters can be easily observed after DISCERN reconstruction (for lineage details see Figure S41B, Figure S42B). The detailed insights into cell differentiation that we obtained with reconstructed data are in stark contrast to the Slingshot results obtained with cite-lq data. While terminal effector molecules can be detected with cite-lq data and seurat-hq data, intermediate stages remain hidden, which prohibits the detection of trajectories and results in a shuffling of marker gene expression (Figures S41 and S42). Taken together these results highlight how expression reconstruction using DISCERN improves downstream analyses and yields deeper biological insights into cell type and state identification, gene regulation, and developmental trajectories of cells.

### 2.4. Discovering COVID-19 disease-relevant cells in lung and blood

The previous sections have demonstrated DISCERN’s utility to reconstruct single cell expression data based on an hq reference, vastly improving the detection of cell (sub-) types and their signaling. Given these advantages, we wondered if DISCERN’s expression reconstruction could deepen our understanding of cell type-composition and signaling changes of immune cells in COVID-19 disease (Figure S43), using two published datasets [53, 24]. To obtain best reconstruction results, we again resorted to using bulk-hq immune reference data (Table S1) [54], as outlined in the previous section.

First, we used a COVID-19 blood dataset (covid-blood-lq) with limited cell type resolution, which was originally analyzed by our group using Seurat (Table S1) [24]. While CD4^+^, CD8^+^, and NK cells formed separate clusters we were unable to visibly distinguish subpopulations of these cells in covid-blood-lq data [24]. Reconstruction of gene expression using bulk-hq data led to the identification of 24 subtypes of CD4^+^ and CD8^+^ T cells in covid-blood-hq data (Figure S44). Several cell clusters identified in covid-blood-hq data showed the correct cell type-specific marker gene expression in covid-blood-lq data, albeit in fewer cells, reduced in magnitude, and in some cases less specific (Figures S45 and S46). Reconstruction also led to the identification of CD4^+^ TH17 helper cells that express *RORC* Figure 4A & B, Figure S47). Based on the molecular footprint of these TH17 cells they were further subdivided into TH17_cluster1 that exhibits a memory T cell phenotype with elevated *IL7R* expression and TH17_cluster2 that exhibits an activated T cell phenotype with elevated *MHC-II, CCR4* and *RBPJ* expression (Figure 4B, Figure S47). The expression of *RBPJ* is of particular interest, as it is linked to TH17 cell pathogenicity, suggesting a role of pathogenic TH17 cells in COVID-19 [55]. It is common practice to stimulate memory T cells in vitro to trigger IL-17A production and a shift towards a TH17 phenotype was previously described in COVID-19 [56]. With DISCERN we are able to distinguish these cells in COVID-19 patient blood without stimulation, identifying cytokine producing memory cells with a TH17-like phenotype (Figure S47).

**Figure 4:**
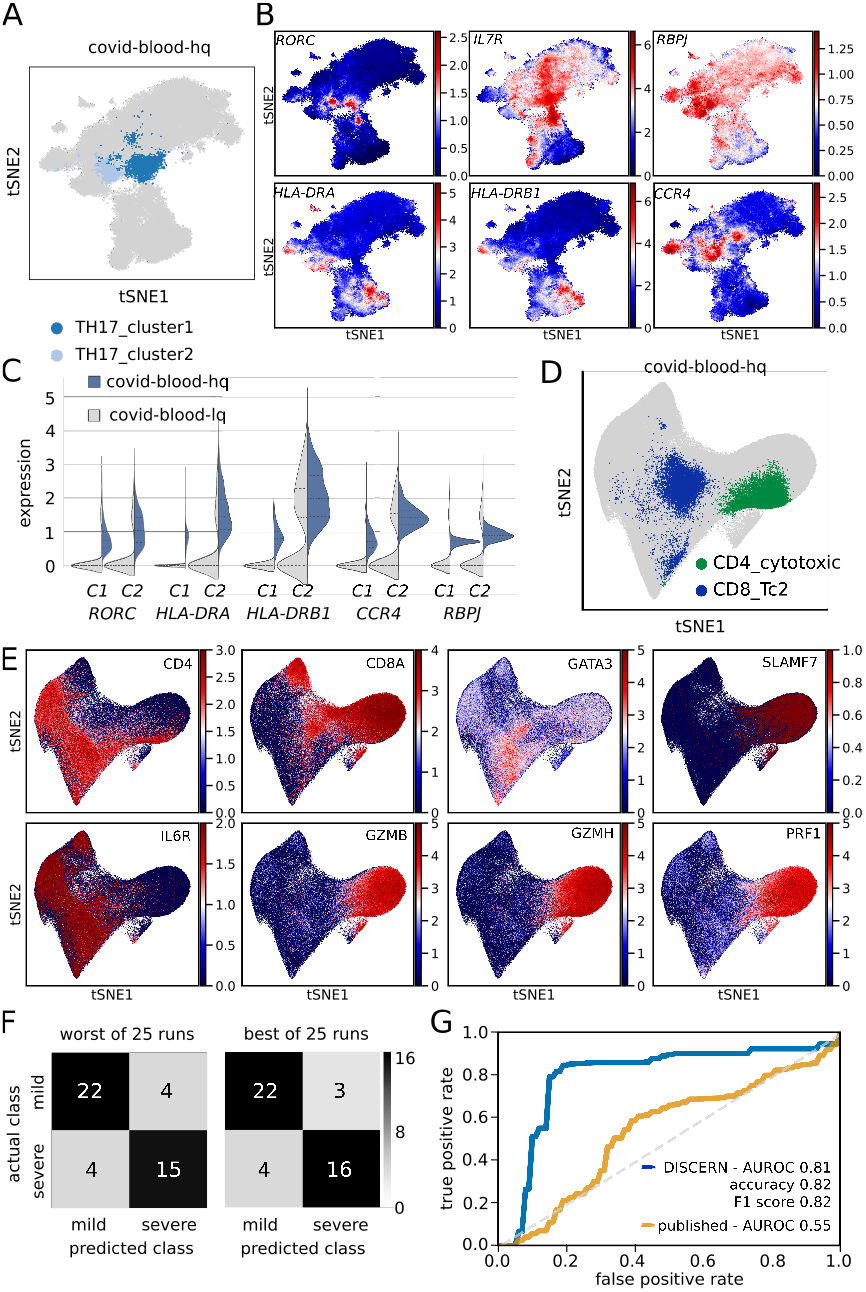
Expression reconstruction improves COVID-19 cell type identification and allows for efficient disease severity prediction. Two COVID-19 blood datasets were reconstructed and analyzed. Hamburg covid-blood-lq and covid-lung-lq data was reconstructed using bulk-hq data, resulting in the respective-hq datasets. Similarly, Cambridge covid-blood-severity-lq data, which contains disease severity information, was reconstructed using bulk-hq data. **A**: t-SNE representation of TH17 subclusters using reconstructed covid-blood-hq data. Clusters were defined using the leiden clustering algorithm on CD4^+^ T cells. **B**: t-SNE representation colored by expression of reconstructed genes distinguishing TH17_cluster1 and TH17_cluster2 cells. TI I17_cluster1 displays a central memory and TH17_cluster2 a more activated phenotype. **C**: Violin plots of expression levels for genes distinguishing TH17_cluster1 (C1) and TH17_cluster2 (C2) cells before (covid-blood-lq) and after (covid-blood-hq) reconstruction with DISCERN. **D**: Rare and unexpected cell types found in the reconstructed covid-blood-hq data with covid-blood-severity and bulk data. Cytotoxic CD4^+^ T cells (CD4_cytotoxic) are displayed in green, CD8^+^ Tc2 helper cells (CD8_Tc2) in blue, and all other cells in gray color. **E**: t-SNE representation of key marker genes in covid-blood-hq data for CD4_cytotoxic and CD8_Tc2 cells displayed in **D**. **F**: Best and worst confusion matrix for disease severity prediction using GBM classifiers trained on fractions of five T cell types (CD4_EM, CD4_cytotoxic, CD4_naive, CD8_EM, CD8_effector) using reconstructed covid-blood-severity-hq data. Category “critical” was combined with “severe” and “asymptomatic” with “mild”. **G**: ROC curve of the GBM predictions outlined in **F** using reconstructed (blue color) covid-blood-severity-hq (CD4_CM, CD4_cytotoxic, CD4_naive, CD8_EM, CD8_effector) and published T cell information from uncorrected (yellow color) data (CD4.CM, CD4.Tfh, CD8.EM, NKT, Treg). Confidence intervals (color shades) indicate one standard deviation.

To further validate the existence of activated TH17 cells in COVID-19 patient blood, we next analyzed the corresponding lung data (covid-lung) of the patients for shared T cell receptor clones (Figure S48). The underlying assumption is that cells with the same T cell receptor in lung and blood originate from the same progenitor and therefore have a high probability of belonging to the same cell type. For this comparison we used the cell type annotation and representation of our original analysis of the covid-lung data, in which memory T and TH17 cells were readily observed without reconstruction [24]. TH17_cluster1 cells showed strong clonal overlap with covid-lung CD4^+^ memory T cells (Figure S48) and expressed comparable levels of *RORC* to covid-lung effector memory TH17 cells (Figure S49), indicating that these CD4^+^ central memory T cells could be TH17 (-like) cells. TH17_cluster2 in blood exhibited strong clonal overlap with effector memory and resident memory TH17 cells in covid-lung data (Figure S48) that express *RORC* and *IL-17A* (Figure S49). Using the clonotype information of resident memory cells producing *IL-17A* in inflamed lung (TRM17), we further corroborated the existence of the newly identified population of IL-17A-producing TH17 cells in reconstructed COVID-19 blood data (Figure S48). In general, the T cell receptor clonal information in blood and lung therefore corroborated our cell type annotation in covid-blood-hq data.

To understand the role of T cell subtypes in COVID-19 disease progression we analyzed a second blood single cell dataset (covid-blood-severity-lq) containing disease-severity information for 130 COVID-19 patients [53]. To obtain optimal cell type resolution, we combined the covid-blood-severity-lq T cell data [53] with CD3^+^ covid-blood-lq cells [24] and reconstructed gene expression for the combined dataset using bulk T cell sequencing reference data[54], resulting in covid-blood-severity-hq data. Many of the 15 CD4^+^ T cell clusters identified in covid-blood-severity-hq data (Figure S50) were also present in the covid-blood-hq data, further validating the consistency of our cell type identification. This is also corroborated by the available surface protein data for covid-blood-severity data, substantiating that naive cells are CD45RA, memory cells are CD45RO, and effector cell types are CD45RO positive (further details in Figure S51). We compared the clusters that we identified in the covid-blood-hq with clusters identified in the covid-blood-severity-hq data and found confined and overlapping regions of TFH, TH17_cluster1, and TH17_cluster2 cells (Figure S52). We also compared the identified clusters to clusters defined in the original publication (Figure S53). Cells identified as TFH in the original publication show significant overlap with naive CD4^+^ T cells (defined on transcriptome and protein level) and CD4^+^ IL22^+^ cells (CD4.IL22) show marked overlap with TREG cells. These results confirm once more the precise and robust cell type identification that can be achieved with DISCERN.

Interestingly, we also identified two rather unexpected cell types after reconstruction. One cluster is positive for *CD4* and negative for *CD8A* while otherwise expressing a signature of CD8^+^ effector memory cells with high expression of *GZMB, GZMH* and *PRF1* (Figure 4D & 4E). This signature points to a CD4^+^ cytotoxic phenotype and indeed virus-reactive CD4^+^ cytotoxic cells were described to be increased in blood during COVID-19 [57]. The other cell type expresses *CD8, IL6R*, and *GATA3,* while being negative for *SLAMF7* (Figure 4D & 4E). These cells were described in the literature to be CD8^+^ T helper cells [58], exert T helper function, and have been shown to lack cytotoxicity. They lack expression of a significant number of cytokines and key transcription factors pointing to a TH17 or TH22 phenotype. On a protein level these cells express *CCR4,* while being negative for CCR6, making them cytolytic CD8^+^ T helper type 2 cells (Tc2) cells. Part of this cluster overlaps with CD4 single-positive cells and might explain why T helper type 2 cells are missing in the CD4 cell clustering.

Overall, the highly specific and sensitive cell type identification in covid-blood-severity-hq data enabled us to correlate the five COVID-19 disease severity categories to shifts in cell type and activity information. We first validated the decrease in TFH cells with increasing disease severity, as described in the original work (Figure S54) [53]. TH17 cells have been extensively studied using flow cytometry and in accordance with our results MHC-II positive as well as *CCR4* positive cells were described in COVID-19 patients (Figure 4B) [56]. We observed a strong decrease in naive T helper cells in severe disease, most pronounced for naive TREGs, while the fraction of TH17 cells showed little correlation with disease severity (Figure S54). Of the two mixed cell types we detected in COVID-19 data, cytotoxic CD4^+^ cells were increased in moderate and severe disease (Figure S55). A similar increase is visible in patients with severe respiratory disease without COVID-19 (Figure S56) and these cells might therefore be a general marker of severe respiratory illness. Cytolytic CD8^+^ Tc2 cells are increased in patients with severe symptoms and in those who died from COVID-19 (Figure S55) and are described to be reduced after recovery from COVID-19 [59]. This positive correlation and the known role of Tc2 cells in fibroblast proliferation induction and tissue remodeling could pinpoint a mechanistic role of these cells in lung fibrosis as witnessed in severe COVID-19 patients. The possibility to observe these cells in reconstructed single cell data may pave the way to study the functional role of these cells in adverse COVID-19 outcome.

The relatively strong correlation of some cell types with COVID-19 outcome suggests that blood cell fraction information might be used for patient severity prediction. We trained a Gradient Boosting Machine (GBM) using leave-one-out-cross-validation (LOOCV) on the fractions of all T cell types and performed a forward feature elimination, to obtain a sparse, optimal model for patient blood-based severity prediction. We first classified patients into three groups, mild (union of asymptomatic and mild, *n* = 26), moderate (*n* = 26), and severe (union of severe and critical, n = 19), reaching an AUROC of 0.63 (Table S4). We noticed that the mild and moderate groups were indistinguishable for the classifier (Figure S57). Training a GBM classifier on mild and severe cases substantially increased classification performance, reaching an AUROC of 0.81 and accuracy, and F1 score of 0.82 (Table S4, Figure 4F & G). Compared to the original T cell types and fractions reported (accuracy 0.61) [53], DISCERN reconstructed T cell fractions are 33% more accurate in the prediction of COVID-19 disease severity (Figure 4G, Table S4). This classification improvement is remarkable, given that DISCERN has no notion of disease severity when it reconstructs gene expression. These results further demonstrate DISCERN’s precise and robust expression reconstruction that enabled the discovery of a potential new blood-based biomarker for COVID-19 severity prediction.

## 3. Discussion

The sparsity of gene expression information and high technical noise in single cell sequencing technologies limits the resolution of cell clustering, cell type identification, and many other analyses. Several algorithms such as scImpute, MAGIC, DeepImpute, and DCA have addressed this problem by imputing missing gene expression in single cell data by borrowing expression information from similar cells within the same dataset. While gene imputation clearly improves gene expression by inferring values for dropped out genes, it comes with several shortcomings. Andrews and Hemberg (2018) showed that several state-of-the-art imputation tools increase the number of false positives [60] by imputing biological absent genes. Additionally the data generated by imputation methods often violate the statistical assumptions made by downstream algorithms, e.g. negative binomial distribution. Furthermore, imputation relies on the comparison of similar cells with largely absent gene expression information in the same dataset. With DISCERN we approach to gene expression inference of single cell data, by realistic reconstruction of missing gene expression in scRNA-seq data using a related dataset (single cell or bulk RNAseq) with more complete gene expression information. We thus propose to call this procedure ‘expression reconstruction’ to highlight the fundamental difference to classical imputation and refer to the dataset with missing gene expression information as low quality (lq) and the reference dataset as high-quality (hq). We considered a dataset high quality, if it showed a good tradeoff between the mean number of expressed genes and the cell number. For example in the pancreas dataset the smartseq2 (6214.0 genes) and the fluidigmc1 (8127.4 genes) show a the highest number of expressed genes, but the fluidigmc1 batch only consists of 638 cells compared to the smartseq2 batch with 2394 cells, thus we selected the smartseq2 batch as the high-quality batch. However, DISCERN does not require the definition of a high quality batch a priori and it can depend on the scientific question, e.g. a specific batch shows enriched expression of specific genes. In this case the evaluation of multiple reconstructions with different “high quality” batches can be useful. Furthermore, the use of the dropout estimation procedure in the decoder allows to achieve a single-cell data-like distribution of the reconstructed data and thus is not as strongly violating statistical assumption of downstream analysis. Thus, we consider DISCERN as an approach for expression reconstruction including batch correction, where the reference does not need to be defined *a priori* and can come from single cell as well as bulk RNAseq experiments, which enables DISCERN to improve over current state-of-the-art batch correction and imputation methods.

We provide compelling evidence that our reference-based reconstruction out-performs contemporary expression imputation algorithms as well as batch correction algorithms such as Seurat, scGen, scVI, and CarDEC when they are repurposed for expression reconstruction. To obtain an objective and thorough performance evaluation for expression inference, we used seven performance metrics on 19 datasets, including 12 single cell sequencing technologies. These datasets cover a range of differences, both technical and biological. While we do not distinguish them in this work, DISCERN could be conditioned on technical as well as biological differences to, for instance, generate ‘diseased’ expression programs from healthy data. We focused our performance evaluation on three scenarios with available ground-truth information, i) the in silico creation of defined gene and pathway drop out events in scRNA-seq data, ii) published hq and lq data pairs from the same tissue (pancreas, difftec, sn/scRNA-seq datasets), and iii) CITE-seq protein expression as ground-truth for cell types (citeseq dataset). In total, DISCERN achieved best performance in 21 out of 27 experiments. While DISCERN yields first place to other methods in FC expression correlation comparisons, it always obtains best results across all datasets in gene expression, gene regulatory network analysis, pathway reconstruction, and cell type and activity identification and is the most stable algorithm for different lq to hq size ratios and cell type overlaps. Furthermore it reaches best performance in several batch correction evaluation metrics.

It is important to note that DISCERN is a **precise** network that models gene expression values realistically while retaining prior and vital biological information of the lq dataset after reconstruction. The network is also **robust** to the presence of different cell types in hq and lq data, or an imbalance in their relative ratios, and is robust to ‘hallucinating’ hq-specific cells into the lq data. Thus, DISCERN evidently shows less increase in the number of false positives compared to other data smoothing and imputation algorithms. Several algorithmic choices are the foundation of DISCERN’s precision and robustness. The network was designed to model the sequencing-technology-specific and the underlying biological signals in separate components of its architecture. Disentanglement of those two components is necessary to accurately reconstruct expression information in the case where lq and hq datasets have different content, i.e. cell type compositions. If the component designed to model the effect of sequencing technology also captures the difference in the biological signal, the reconstruction will lead to a lack of integration across the two datasets where some cell types are still clustered by dataset (similar to scGen in Figure S27). On the contrary, if the component modeling the biological signal captures sequencing-technology-specific features, the reconstruction will lead to an over-integration of the datasets where cells of different types are mixed together (similar to Seurat in Figure S23). The demonstrated ability of DISCERN to avoid those shortcomings, even in scenarios where there is very little to no overlap between cell types across datasets, lies in the carefully crafted balance between the expressivity of its components. The representational capabilities of DISCERN, achieved via batch normalization, five loss terms, and a dual head decoder, would reduce DISCERN’s usability, if they would require frequent dataset-specific tuning. The stability and usability was therefore a central concern in the design and evaluation phase of DISCERN, which resulted in an algorithm that gave very good results with a single set of default (hyper-) parameters. All comparisons to other algorithms, for instance, were performed with default settings. Only the expression reconstruction of the exceptionally large COVID-19 datasets required the fine-tuning of the learning rate, cross entropy term, sigma, and the MMD penalty term. Another important technical feature of DISCERN is that it can easily be integrated into existing workflows. It takes a normalized count matrix, as created by nearly all existing single cell analysis workflows, as input and produces a reconstructed expression matrix. This can be used for most downstream applications (i.e. cell clustering, cell type identification, cell trajectory analysis, and differential gene expression). DISCERN can be trained on standard processors (CPU) for small and medium-sized datasets and requires graphical processing units (GPU) for the expression reconstruction of large datasets. Altogether, the usability and robustness of DISCERN should enable even non-expert users to perform gene expression reconstruction.

A unique feature of DISCERN is the use of an hq reference to infer biologically meaningful gene expression. While we consider this a main strength of DISCERN, the dependence on a suitable reference dataset might also limit its application. We took great care in this manuscript to mitigate this concern by showing how DISCERN is able to reconstruct gene expression for many different types of lq and hq pairs, ranging from indrop - smartseq2 to single nucleus - single cell data pairs. Remarkable in this context is DISCERN’s robustness to differences between the cell type compositions of lq and hq data pairs, with DISCERN being the only algorithm obtaining robust expression reconstruction when few or no cell types overlap. We have also shown that purified bulk RNA-seq samples can be used as hq reference, as successfully applied to PBMC and COVID-19 datasets in this study. We used 9852 FACS purified immune cell bulk sequencing samples [54], comprising 27 cell types, to successfully reconstruct single cell expression data. This implies that most single cell studies involving immune cells (with or without other cell types present) can be reconstructed with DISCERN using a single published bulk RNA-seq dataset. Furthermore, public RNA-seq repositories such as NCBI GEO contain tens of thousands of samples of immune and non-immune cells that could serve as reference for most expression reconstruction experiments. Conversely, pure cell type or subtype bulk RNA-seq data could be hard to obtain as the sorting of cells might have limited resolution or might be partially impure. In consequence, the usage of bulk RNA-seq data as reference for expression reconstruction could lead to a grouping or averaging of cell subtypes. While these potential caveats might adversely affect expression reconstruction, we have not observed merging or averaging effects of single cell subtypes when corresponding bulk RNA-seq cell type information was not present or present at different proportions (Figure 3B & 3C, Figure S36). Importantly, cells do not necessarily cluster into distinct classes but can build cell continua, as shown in the trajectory analysis in Figure 3B & 3C, where T cells seem to differentiate into each other and do not form clearly separable clusters. In general, handling continua of cell types is challenging for imputation and batch correction algorithms, as many of them, including for instance scGEN, Bfimpute, SIMPLEs, and cscGAN, require or recommend cluster or cell type annotation. This might lead to under- or over-integration of cell continua. DISCERN does not rely on cluster (or cell type) information and seamlessly integrates and reconstructs cell clusters and continua (Figure 3C, Figure S44). In conclusion, we provide strong evidence that DISCERN is widely and easily applicable to many single cell experiments.

While DISCERN gave good reconstruction results using default parameters for most datasets we analyzed, we would like to highlight that the immense representational power of generative neural networks can remove or hallucinate biological information if not properly handled [6]. This is true for data integration [61, 62] as well as for expression reconstruction algorithms and we would highlight two guiding principles for optimal results. For non-expert users, we would recommend the use of default settings and a careful selection of a related hq dataset. When datasets are large and complex, with many cell types in the lq and several non-overlapping cell types in the hq data, one should always ensure that training does not merge or mix non-overlapping cell types with other cells, by investigating that these cells keep their cell type-specific marker gene expression. Keeping these ‘checks and balances’ will usually result in good reconstruction results even for complex datasets such as covid-blood-severity.

To obtain novel insights into COVD-19 disease mechanisms and a new blood-based biomarker for disease severity we reconstructed two published datasets with DISCERN, Hamburg COVID-19 patients (covid-lung, -blood) and the COVID-19 cell atlas (covid-blood-severity). The application of DISCERN to the covid-blood dataset (COVID-19 patient blood) enabled us to detect 24 different immune cell types and activity states, which is quite remarkable given that we find these cells in blood. Two TH17 subtypes caught our attention, as they share the TCR clonality with the lung data from the same patients (covid-lung), suggesting bloodstream re-entry of lung TH17 cells. We linked these two subclusters to their functional role by separating them into a memory-like and activated-like phenotype. The clonal overlap of activated TH17 cells in blood with previously discovered lung-resident cells suggests that activated TH17 cells in blood are resident T cells from the lung reentering circulation. These cells might in part explain the multi-organ pathology observed in COVID-19, as activated T cells might travel via the blood to secondary organs and cause inflammation and tissue damage. Future work might demonstrate the effect of these activated T cells on tissue inflammation.

Given the detailed cell type and activity information we reached with gene expression reconstruction, we wondered if changes in blood immune cell populations might be useful as a biomarker for disease severity prediction. We used DISCERN to reconstruct the covid-blood and the covid-blood-severity datasets and again identified a plethora of different T cell subtypes in the blood of patients with COVID-19. Using these cell proportions, we were able to classify mild and severe disease using a GBM machine learning algorithm with 82% accuracy, outperforming classification with the originally published T cell types by 21 percent points. This improvement is absolutely striking, as DISCERN has no notion of the classification groups. It simply reconstructs gene expression and thereby improves cell type detection. These results are a convincing implicit proof not only of the usefulness of DISCERN but more importantly of its precision and robustness. While the use of this scRNA-seq-based biomarker would be too expensive and time-consuming for clinical care, it strongly suggests that FACS-based T cell fraction or count information from blood could be used to trace and predict the severity state and potentially the disease trajectory of COVID-19 patients.

Interestingly, we also discovered two atypical T cell types in reconstructed COVID-19 patient blood single cell data. While cytotoxic CD4^+^ T cells have been observed in COVID-19, we can show that this increase is not COVID-19 specific and is also observed in other types of pneumonia. Interestingly, we also detected cytolytic CD8^+^ Tc2 cells that express *CD8A, GATA3, IL6R* and are negative for *SLAMF6*. This cell type is linked to tissue fibrosis and steroid refractory disease in asthma [63]. The increase in CD8^+^ Tc2 cells that we observe specifically in COVID-related death could be associated with COVID-19 patients that do not respond to steroids. Demonstration of increase of this cell type in patients dying of COVID-19 points to a potential therapeutic intervention with the drug Fevipiprant, which blocks CD8^+^ Tc2 cell activation and its pro-fibrotic effects by inhibiting prostaglandin D2 signaling [64]. Functional analysis of these cells has to demonstrate whether these cells are an early marker of later death or whether it is a marker of already escalated treatment.

The basic concept of utilizing a high-quality reference to improve lower quality data might be applied to many other research areas where technological limitations restrict biological insights. The usage of deep generative networks and other artificial intelligence methodology to infer information beyond what is technically measurable could be transformative in future biomedical research.

## Acknowledgements

We thank Immo Prinz, Manuel Friese, Johannes Soeding, Robert Zinzen, Yu Zhao and Stefan Kurtz for their helpful comments and suggestions. We thank Rajasree Menon and Matthias Kretzler for providing kidney single cell and single nuclear RNA-seq data and their support for corresponding analysis. FH, RK, MM, PM, & SB were supported by the LFF-FV 78, EU ERare-3 Maxo-mod, SFB 1286 Z2, FOR 296, and FOR 5068 research grants. Further support was obtained from the UKE R3 reduction of animal testing grant. CE was supported by DFG ER 981/1-1 and the clinician scientist programme of university Hamburg, NG was supported by ERC StG-715271, and SHu was supported by ERC CoG-865466 and has an endowed Heisenberg-Professorship awarded by the Deutsche Forschungsgemeinschaft. SHa was funded by the BMBF STOP-FSGS-01GM1518C and SFB 1192 B08 research grants.

## Author contributions

SB initiated and SB, PM, FH, and CE conceptualized the study with help from MM. FH and CE implemented DISCERN, MM refactored the code, and PM reviewed the DISCERN implementation. FH, CE, and RK performed the analyses. SB, PM, NG, and SHu supervised the study. SB, FH, and CE wrote the manuscript. SHu, PM, NG, RK and SHa provided ideas, contributed to the manuscript text and critically reviewed the manuscript. All authors read and approved the final manuscript.

## 4. Methods

### 4.1. Data availability

In this manuscript many different scRNA-seq and RNA-seq datasets were used. A comprehensive overview of dataset, method, cell type, origin, size, and naming convention can be found in Tables S1 to S3. All datasets are publicly available as listed in Table S1.

### 4.2. Dataset description

#### Pancreas

The pancreas dataset is a collection of different scRNA-seq datasets, profiling pancreas cells in the context of diabetes [65]. The pancreas dataset is a widely used dataset for batch correction benchmark experiments and due to its high number of cell types and sequencing technologies it allows to evaluate differences between cells and sequencing technologies at the same time. The expression table, including the annotation, is available from SeuratData (https://github.com/satijalab/seurat-data) as panc8.SeuratData (v3.0.2) [65]. The dataset was sequenced using five sequencing technologies (Smart-Seq2, Fluidigm C1, CelSeq, CEL-Seq2, inDrop) and consists of 13 cell types (alpha, beta, ductal, acinar, delta, gamma, activated_stellate, endothelial, quiescent-stellate, macrophage, mast, epsilon, schwann). In total, before preprocessing, the dataset contains 14 890 cells.

#### difftec

The difftec dataset was created for a systematic comparative analysis of scRNA-seq methods [66]. Similar to pancreas, the difftec dataset is ideal for the evaluation of expression reconstruction across many cell types and sequencing technologies. Seven sequencing technologies (10x Chromium v2, 10x Chromium v3, Smart-Seq2, Seq-Well, inDrop, Drop-seq, CEL-Seq2) were used with at least two replicates each. In this dataset 10 different cell types (Cytotoxic T cell, CD4^+^ T cell, CD14^+^ monocyte, B cell, Natural killer cell, Megakaryocyte, CD16^+^ monocyte, Dendritic cell, Plasmacytoid dendritic cell, Unassigned) were annotated, and make up for 31 021 cells in total before filtering. The expression table including the annotation is available from SeuratData as pbmcsca.SeuratData (v3.0.0).

#### snRNA & scRNA

The dataset was created for the validation of a single cell and single nuclei analysis toolbox [38]. Since snRNA-seq and scRNA-seq data varies in the amount of counts per cell and the genes detected, we tested if DISCERN could reconstruct snRNA-seq expression so that it would closely resemble scRNA-seq expression, providing a biological ground-truth. While we label snRNA-seq data as lq and scRNA-seq as hq, this distinction is incorrect from a biological perspective, as gene expression should be in part different between the nucleus and the cytosol. The dataset consists of a liver biopsy sample (HTAPP-963) of metastatic breast cancer with single cell sequencing and single nuclei sequencing. Eight cell types (Epithelial cells, Macrophages, Hepatocytes, T cells, Endothelial cells, Fibroblasts, B cells, NK cells) were found in the original publication in a total of 12 423 cells. The data was sequenced using the Chromium V3 technology on a Illumina HiSeq X sequencer.

#### covid-lung & covid-blood

The COVID-19 dataset we have previously published consists of blood and bronchoalveolar lavage (BAL) samples from four patients with bacterial pneumonia and eight patients with SARS-CoV-2 infection[24]. In total 155 706 cells were sequenced using TCR-seq technology, which allows for the comparison of clonal expansion in both tissues. While we investigated the lung data in detail in the original publication, the analysis of the blood was largely limited to cell type identification. Using DISCERN, we use the blood data to find previously unobserved cell types, link them to cell clones found in the lung, and derive a biomarker based on cell fractions (see also covid-blood-severity data). Cell type annotations for the BAL samples were used as in the original publication.

#### citeseq

This dataset contains CITE-seq information of healthy human PBMCs for 6 cell types (B cells, CD4 T cells, NK cells, CD14^+^ Monocytes, FCGR3A+ Monocytes, CD8 T cells) [39]. In our analyses we used the cell type information provided in the original publication [67]. The CITE-seq data is ideal to benchmark DISCERN, as the information of 13 surface proteins offers ground-truth information on the cell types and a good proxy for the expression of the 13 corresponding genes.

#### bulk

We used this large dataset of 28 FACS sorted and bulk sequenced immune cell types as ‘ultimate’ hq reference data for lq immune single cell sequencing data. Each of the 9852 samples provides an average expression information for 13 104 genes for a specific immune cell type, providing a hq reference for e.g. lq single cell PBMC CITE-seq data with only 798 expressed genes per cell. We further assume that this dataset is large enough to provide enough per cell type variability for our deep neural network to faithfully learn and represent its gene expression. In more detail, the dataset consists of 28 sorted immune cell types (Naive CD4, Memory CD4, TH1, TH2, TH17, Tfh, Fr. I nTreg, Fr. II eTreg, Fr. III T, Naive CD8, Memory CD8, CM CD8, EM CD8, TEMRA CD8, NK, Naive B, USM B, SM B, Plasmablast, DN B, CL Monocytes, Int Monocytes, NC Monocytes, mDC, pDC, Neutrophils, LDG) with ¿ 99% purity [54]. Total RNA was extracted using RNeasy Micro Kits (QIAGEN). Libraries for RNA-seq were prepared using SMART-seq v4 Ultra Low Input RNA Kit (Takara Bio). In total, the dataset contains 9852 samples collected in two phases from 416 donors, out of which 79 are healthy. For training DISCERN, bulk TPM counts and all cell types were used if not stated otherwise.

#### covid-blood-severity

This dataset is an aggregation of three COVID-19 sequencing studies using the 10X Genomics Chromium Single Cell 5’ v1.1 technology. It contains a large number of cell types with fine-grained cell type annotations that are complemented with information on COVID-19 disease severity for each patient sequenced. We used this dataset to obtain a blood-based biomarker of COVID-19 disease severity, based on T cell fractions observed with DISCERN. The data consists of PBMCs from 29 healthy, 89 COVID-19 and 12 LPS-treated patients. The authors detected 51 cell types in their original work (see Table S1) [53] and COVID-19 patients were classified by their disease severity (worst clinical outcome) into ‘asymptomatic’, ‘mild’, ‘moderate’, ‘severe’, ‘crit-ical’, and ‘death’. Count data together with CITE-seq information was used as provided in the original publication (https://covid19.cog.sanger.ac.uk/submissions/release1/haniffa21.processed.h5ad).

#### kidney-lq (snRNA-seq) & kidney-hq (scRNA-seq)

The kidney dataset consists of single cell RNA-seq and single nuclei RNA-seq data of 9 patients with acute kidney injury sequenced using 10X Genomics Chromium technology. It contains in total 82 701 cells with 52 934 cells sequenced using snRNA-seq and 29 767 cells sequenced using scRNA-seq. The dataset does not contain cell type annotation, but in initial analysis using a different subset [68] suggested that identification of T cells in the snRNA-seq data is challenging. For this reason, the analysis was focused on the detection of T cells and their subtypes.

### 4.3. Code availability

All original code has been deposited at github.com (https://github.com/imsb-uke/discern) and is publicly available as of the date of publication. Any additional information required to reanalyze the data reported in this paper is available from the lead contact upon request.

### 4.4. Preprocessing

Raw expression data (Counts) preprocessing was performed as previously described [69] using the scanpy (v1.6.1, [70]) implementation. In particular, the intersection of genes between batches was used. The cells were filtered to a minimum of 10 genes per cell and a minimum of 3 cells per gene. Library size normalization was performed to a value of 20 000 with subsequent log-transformation. As model input for DISCERN the genes were scaled to zero mean and unit variance. However, for all further evaluation the genes were scaled to their uncorrected mean and variance not considering the batch information.

### 4.5. Description of DISCERN

DISCERN is based on a Wasserstein Autoencoder with several added and modified features. We will describe the details of DISCERN’s architecture in the next paragraphs and a compact representation can be found in Figure S1B.

#### Wasserstein Autoencoder

While neural network-based autoencoders have been widely used for decades for dimensionality reduction [71, 72], recent advances have also allowed their use to build a generative model of the distribution of the data at hand[73]. More recently, leveraging results from optimal transport [74], Wasserstein Generative Adversarial Networks (WGAN) [75] and Wasser-stein Autoencoders (WAE) [25] have been designed to explicitly minimize the (Wasserstein, or earth-mover) distance between the distribution of the input data and their reconstruction. WGANs only implicitly encode their input into a latent representation (called latent code), while WAE has the useful property of using an explicit encoder, which makes it possible for the model to directly manipulate the different representations of single-cell data. Finally, the WAE framework, established in [25], allows the use of a wide range of architecture and losses, which we are going to detail now. First of all, in order to effectively use a number of latent dimensions that adaptively matches the intrinsic dimension of the scRNA-seq data at hand, DISCERN uses a random encoder as prescribed in [76].

#### Architecture

Autoencoders widely used for transcriptomics applications are shown to perform well on several tasks, like drug perturbation prediction [23] or dropout imputation [12]. Since the ordering of the genes in scRNA-seq count matrices is mostly arbitrary, fully-connected layers are usually used in this task. In our case, DISCERN consists of three fully connected layers in the encoder and the decoder. The bottleneck of the autoencoder (or latent space) contains 48 neurons, which is sufficient to accurately model all the datasets we used in our experiments. Additionally, we exploit a finding from [76] to let the network learn the appropriate amount of latent dimensions. While the encoder will be tasked to transform the distribution of the input data into a fixed, low-dimensional prior distribution (i.e. a standard Gaussian), the decoder will perform the opposite, i.e. transforming the fixed, low-dimensional prior distribution into gene space. scRNA-seq data is known to display a high level of zero measurements, called dropout, which is essential to accurately model the count distribution. To describe scRNA-seq data in a parametric way, it is common to model the expression level of a gene with zero-inflated negative binomial distribution [77]. Despite the several non-linearities in the decoder architecture, it is, however, difficult to learn an encoding function that maps a simple prior to the distribution leading to low quality modeling of low expressed genes. To address this issue, we scale the gene expression and attach a second head to the decoder (i.e. a second decoder sharing all weights with the first, except for the last layer). The task of the second decoder head is to predict, for each gene of a cell, the probability of its expression to be dropped out, giving rise to a random decoder. Thus, this second decoder head predicts dropout probabilities and models the dropout probabilities for different batches. This additional head allows modeling the dropout and the expression independently, to capture the specific distribution of single cell data without the need for further explicit assumption about the distribution. During inference the predicted expressions are randomly set to zero based on these predicted dropout probabilities. This sampling procedure does not have any trainable parameter, is therefore not part of the model training and only performed during inference.

#### Loss function

The loss optimized during the training of DISCERN is composed of four terms: a data-fitting (or reconstruction) loss, a dropout fitting (cross entropy) loss, a prior-fitting term (ensuring that DISCERN approximately minimizes the Wasserstein distance) and a variance penalty term (that controls the randomness of the encoder). Thus, DISCERN can be considered as a Wasserstein Autoencoder as introduced in [25]. For the reconstruction term, the framework introduced in [25] allows the use of any positive cost function. We elected to use the Huber loss [78] as it is well suited for modeling scaled scRNA-seq expression data, because it allows to select a threshold value to give lower weight to high differences in highly expressed genes and thus allows the model to learn a more robust expression estimate without focusing too much on outlier values. This reconstruction term is defined

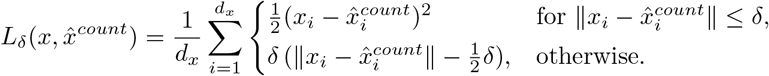

as where *x* is the input expression matrix, 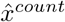 the predicted expression matrix from one decoder head, *d_x_* the number of genes, and *δ* a threshold deciding between the two conditions of the Huber loss.

For the prior-fitting term, following [25], DISCERN uses the Maximum Mean Discrepancy (MMD) [79] between the aggregate posterior (i.e. the distribution of the input single-cells after encoding) and a standard Gaussian. We use the sum over an inverse multiquadratic kernel with different sizes for this task.

Similar to [79], we define the MMD

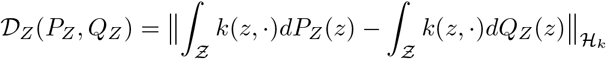

as where *P_Z_* is the gaussian prior distribution and *Q_Z_* the aggregated posterior in the latent space for a positive-definite reproducing kernel 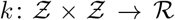 and a corresponding real valued reproducing kernel hilbert space 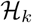. For the implementation details please refer to [25] or the provided implementation.

Then, to prevent the random encoder (with diagonal covariance) from collapsing to a deterministic one, a penalty term that enforces that some components of the variance are close to 1. Intuitively, that means that the superfluous latent dimensions will only contain random noise (see [76] for more details). We define this penalty term

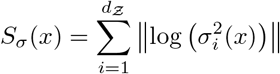

as where *d_z_* is the number of latent dimensions and *σ* the function generating the components of the variance in the latent space, in our case, the encoder network.

Another loss term, namely the binary cross-entropy loss, on the second decoder head is used to enable the model to learn a dropout probability for each gene and sample. The loss on the dropout layer enables the model to capture the bimodal distribution of single cell data. We define the binary cross-entropy loss as

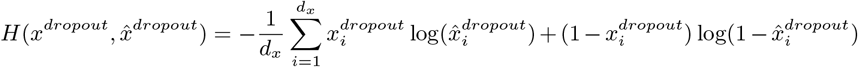

where *x^dropout^* is the binarized expression information, 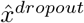 is the predicted binarized expression (probability of dropout) and *d_x_* the number of genes. Additionally, activity regularization is applied on the Conditional Layer Normalization (CLN), such that the weights of the conditional layers are only regularized in a batch-specific manner and the regularization is not applied for batches, which are not present in the current mini-batch. This has the advantage that the batch dependent weights are not influenced too much by different batch sizes. The four loss terms are added (and weighed using λs) together to form the loss that DISCERN minimizes during training:

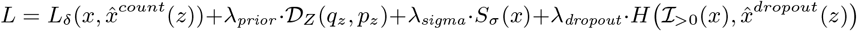

See also Figure S1B for a graphical depiction of the loss terms.

#### Conditional Layer Normalization

The weights of those fully-connected layers are shared for all the batches that DISCERN is trained on. However, to model the batch-specific differences, we use a Conditional Layer Normalization (CLN) that applies the idea proposed in [27] to Layer Normalization [28] after each fully connected layer (see Figure S1B). In essence, for each batch, different sets of shifting factors are learned. Note that in DISCERN, no scaling factors are used to limit the expressivity of the conditioning and therefore reduce the chance of over integration. This allows not only to accurately model the batch-specific differences between batches, but also to transfer the batch effect from one dataset onto another, in the spirit of the style-transfer approach developed in [27]. To make things clear, DISCERN does not explicitly train to integrate datasets. Instead, it trains to accurately model the input data, capturing the batch-specific differences with the weights of the CLN layers (i.e. conditioning), and the biological signal (which is mostly shared across the batches to integrate) with the weights of the fully-connected layers. After training, we encode all the cells we want to reconstruct, conditioning the process on their batch of origin. Then, we take the batch chosen by the user and proceed to decode all the cells conditioning on that specific batch, effectively transferring the batch effect of one specific batch onto all of the batches we want to integrate and reconstruct. The training loss is computed over the complete minibatch, thus it is not different per batch (dataset). The weights of the conditional layer normalization are learned together with the weights of the feed-forward network using the same loss function.

#### Activations & dropout

With the exception of the output layer, every other fully-connected layer of the encoder and the decoder was followed by a CLN, a Mish ([80] activation function, and dropout during model training to reduce overfitting.

#### Optimization

To optimize the weights of our model, DISCERN uses Rectified Adam ([81], which addresses some of the shortcomings of the widely used Adam [82] and generally yields more stable training. To prevent overfitting, the optimization is stopped early. It is implemented as a modification of the Keras EarlyStopping (with parameter minDelta set to 0.01 and the patience to 30) where the callback is delayed by a fixed number of 5 epochs. The delay was implemented to prevent too early stopping due to the optimization procedure.

#### Reconstruction

The reconstruction (or projection) to a reference batch is not performed during training and thus the network is not optimized to it. However, during inference, the reconstruction can be performed by providing the correct batch label in the encoder part of the network, while only providing the reference batch label for the decoder part. Therefore, The network will encode the dataset to a batch-independent latent representation and decode it using only the reference label and therefore project the complete dataset to the reference batch. This can be done for any number of batches without re-training of DISCERN.

#### Running time and memory usage

DISCERNs running time for training is linear in the number of cells and the number of training epochs. However, the use of the early stopping mechanism greatly reduces the running time and improves model performance. Additionally the running time, for training and inference, is dependent on the size of the mini-batches. The memory requirements are also linear in the number of cells and genes for training and inference. Since DISCERN is trained on mini-batches the memory requirements can also be slightly adjusted by changing the mini-batch size during training or inference.

### 4.6. Hyperparameters

As outlined in the architecture section of the methods and depicted in Figure S1, DISCERN features several learnable hyperparameters. The complexity of the hyper-parameter search space is a potential downside of DISCERN, if these hyperparameters would be unstable across different datasets or in other words, would require constant tuning. Fortunately, DISCERN’s hyperparameters are very stable across the multitude of datasets tested in this manuscript, which we will outline in this paragraph. Naturally, there is no rule without an exception, which in this manuscript are the COVID-19 datasets that required optimization for several hyperparameters.

#### Constant hyperparameters

DISCERN features a number of hyper-parameters that can be tuned through hyperparameter optimization (see below for details). Most of them have default values that yield reasonable performance across the different datasets we used and are being kept constant across experiments, including the COVID-19 dataset. Those constant hyperparameters are: the choice of the reconstruction loss (Huber loss), activation functions (Mish), CLN for the conditioning, number of fully-connected layers (3) and their size (1024, 512, 256 and 256, 512, 1024 neurons for the encoder and the decoder respectively), number of latent dimensions (48), learning rate (1 × 10^−3^), decay rates *β*_1_ and *β*_2_ of Rectified Adam (0.85 and 0.95 respectively), batch size (192), label smoothing for our custom cross entropy loss (0.1), dropout rates (0.4 in the encoder and 0 in the decoder), delta parameter of the Huber loss (9.0), weight on the penalty on the randomness of the encoder λ_*sigma*_ (1 × 10^−8^), weight on the cross entropy loss term λ_*dropout*_ (1 × 10^5^), weight on the MMD penalty term λ_*prior*_ (1500).

#### Dataset-specific hyperparameters

The optimal value of the L2 regularization applied on the weights of our custom CLN highly depends on the dataset at hand and thus requires dataset-specific tuning. For datasets with a very small variance in cell compositions the L2 CLN regularization can be turned off (weight set to 0). When datasets have different compositions the L2 CLN regularization requires higher values (typically between 1 × 10^−3^ and 0.2).

#### COVID-19 hyperparameters

For the experiments with COVID-19 datasets slightly adjusted hyperparameters were used: learning rate of 6e-3, label smoothing for our custom crossentropy loss of 0.05, weight on the penalty on the randomness of the encoder λ_*sigma*_ (1e-4), weight on the cross entropy loss term λ_*dropout*_ (2e3), weight on the MMD penalty term λ_*prior*_ (2000).

#### Hyperparameter optimization

DISCERN implements different techniques for hyper-parameter optimization by using the ray[tune] library [83]. For most use cases the model does not require hyperparameter tuning and the default parameter should be sufficient. However, DISCERN has a generic interface and supports nearly all techniques implemented in ray[tune]. The initial hyperparameters were found using grid search. The loss used for the hyperparameter selection is the classification performance of a Random Forest classifier trying to classify real vs. auto-encoded cells. Classification performance was measured using the area under the receiver operating characteristic curve (AUC / AUROC).

### 4.7. Competing algorithms and methods

We briefly discuss competing methods and have compared their performance to DISCERN in the results section. These algorithms can be grouped into two categories, i) imputation algorithms that were developed to estimate drop-out gene expression based on dataset inherent information (MAGIC, DCA, scImpute) and ii) algorithms designed for batch correction that we have modified or extended to reconstruct gene expression, although this is not their intended use (Seurat, scGen). Given the latter, it is clear that DISCERN could be used purely for batch correction in latent space, a subject beyond the scope of this manuscript.

#### MAGIC

[13] Markov affinity-based graph imputation of cells (MAGIC) denoises and imputes the single-cell count matrix using data diffusion-based information sharing. The construction of a good similarity metric is challenging for finding biologically similar cells due to high sparsity. MAGIC finds a good similarity metric using a sophisticated graph-based approach that builds less-noisy cell-cell affinities and information sharing across cells. A particular focus of MAGIC was to understand gene-gene relationships and to characterize other dynamics in biological systems. MAGIC is provided as a Python package.

#### DCA

[11] is a deep learning-based method for denoising single-cell count matrices. DCA is implemented in Python and uses an autoencoder with a Zero-Inflated Negative Binomial (ZINB) loss function. For each gene, DCA computes gene-specific parameters of ZINB distribution, namely dropout, dispersion and mean. By modeling gene distributions as a noise model and also computing dropout probabilities of each gene, DCA is able to denoise and impute the missing counts by identifying and correcting dropout events.

#### scImpute

[12] Similarly to MAGIC, scImpute focuses on identifying cells that are similar, which is challenging due to the high sparsity of single-cell count matrices. scImpute is a statistical model using a three step process to impute scRNA-seq data. In the first step spectral clustering is applied on principal components to find neighbors, which later can be used to detect and impute dropout values. In the second step scImpute fits a mixture model of a Gamma distribution and a Normal distribution to distinguish technical and biological dropouts. In the last step, the model uses a regression model for each cell to impute the expression of genes with high probability of dropout. With this approach, scImpute avoids hallucinations and keeps the gene expression distribution. scImpute is provided as an R package.

#### Seurat

[26] is an open-source toolkit for the analysis of single cell RNA-sequencing data. In addition to general analysis functions, Seurat offers batch-correction functionality. Seurat uses canonical correlation analysis to construct this lower dimensional representation and tries to find neighbors between batches in this shared space. These anchors are filtered considering the local neighborhood of the cell pairs and remaining anchors are finally used to construct correction vectors for all cells in this low dimensional representation. While Seurats is intended to work in a lower dimensional representation, it can also be used to reconstruct the expression information from this lower dimensional representation. Seurat is provided as an R package.

#### scGen

[23] is a variational autoencoder based deep learning method with a focus on learning features that help distinguish responding and non-responding genes and cells. scGen constructs a latent space in which it estimates perturbation vectors associated with a change between different conditions. Since scGen models the perturbation and infection responses in single cells, it is focused on in-silico screening with the use of cells coming from healthy samples.It can also be used for batch correction. For batch correction, and unlike DISCERN or Seurat, scGen uses both batch and cell type labels. scGen is built using the scvi-tools toolbox and implemented in python and pytorch.

#### Multigrate

[19] multigrate is an autoencoder based deep learning method developed for the integration of different modalities to improve single cell RNA-seq downstream analysis, mainly clustering. The main focus is the integration of CITE-seq protein abundance since it is often available together with scRNA-seq. They use individual encoders for each modality and build a shared latent representation by partially sharing the decoder. Multigrate is built using the scvi-tools toolbox and implemented in python and pytorch.

#### scVI

[36] is a variational autoencoder-based deep learning method developed for several single cell analysis approaches like batch correction, clustering, and differential expression analysis. It models expression data using a zero-inflated negative binomial loss during the training. For comparison of scVI to other models, only the batch correction functionality was used. For the differential expression analysis we used the same workflow as for the other methods to allow for a fair comparison. scVI is implemented in python and pytorch.

#### CarDEC

[14] is an autoencoder-based learning method developed for batch effect correction, denoising of expression data and cell clustering. The CarDEC pipeline computes highly variable genes across all batches and pre-trains an autoencoder to reconstruct the expression of these genes. In a second step, the weights are transferred to a bigger network, which is trained jointly on the highly variable and lowly variable genes using two reconstruction losses. Additionally, they include a self-supervised clustering loss in the latent space to improve batch mixing. CarDEC is implemented in python and Tensorflow.

#### DeepImpute

[15] is an ensemble method consisting of multiple autoencoder-like deep neural networks, where each network is trained to learn the relationship between a set of input genes and a set of target genes. Input and target gene sets are selected based on correlation of gene expression values. The estimated expression values from each of the networks is combined to yield the final imputed dataset. DeepImpute is implemented in python and Tensorflow.

#### trVAE

[35] is a variational autoencoder based deep learning method developed for the generation of unseen samples or conditions of single cell RNA-seq data. It uses an encoder with additional inputs for encoding the condition and a decoder which gets, together with the latent code, the target condition as input. To achieve a condition independence the first layer is regularized using maximum mean discrepancy. trVAE is implemented in python and Tensorflow.

### 4.8. Evaluation metrics

#### t-SNE & UMAP

For visualization of the datasets and to qualitatively assess the integration performance tSNE and UMAP were used. Both methods are based on PCA representation and use non-linear representations to create a 2D representation of the data. We used the scanpy [70] implementation. Default settings were used in nearly all cases except: In the combined COVID-19 dataset analogue to Kobak *et al.* [84] the dataset was subset to 25 000 cells and tSNE was computed using a perplexity of 250, and a learning rate of 25 000/12. These positions were taken and used as input to tSNE of all cells using a perplexity of 30 a learning rate of (number of observations)/12 and a late exaggeration of 4.0 using FIt-SNE [85]. Clustering was performed using PARC [86] with default parameters except dist_std_local=1.5 and small_pop=300. Methods were changed here due to computation time issues for 350 000 cells. covid-blood data was analyzed using a learning rate of (number of observations)/6 a perplexity of (number of observations)/120 and early_exaggeration=4. Clustering was performed using default parameters except knn=100 and small_pop=100 to reduce the number of clusters with limited cell number. Clustering of the T helper cells in healthy blood was performed using coarse clustering with 30 nearest neighbors and leiden clustering (https://github.com/vtraag/leidenalg) with a resolution of 0.6. Afterwards a combined cluster of IFN-regulated and TREG was reclustered using a resolution of 0.4 and effector T cells were reclustered using a resolution of 0.8. Resolution was chosen to dissect the raw gene expression changes of known cell types.

#### Mean gene expression

Mean gene expression was calculated as average over log-normalized expression over all cells, usually stratified by celltype. This evaluation of expression data consists of many data points where several have values close to zero, but could have a high weight on rank-based correlation methods. Thus Pearson correlation was used to evaluate the performance.

#### Differential gene expression

Differential gene expression was performed using the scanpy [70] rank_gene_groups function using the t-test method for calculating statistical significance on log-normalized expression data. Differential gene expression analysis was always performed under consideration of the cell type information. For comparison of differential gene expression analysis between conditions, the Pearson correlation was used. It is calculated either on the log2 fold-change or in most cases on the t-statistics, computed during significance estimation. The data was compared using the t-statistics, because it aggregates information on both the variance and the change in mean expression. Thus it allows, roughly speaking, for simultaneously evaluating the significance and the log2 fold change. Usually all available genes were used for correlation, except in the in-silico gene removal experiment, where only the removed genes were considered. We used spearman rank correlation when all genes were available and pearson correlation otherwise.

#### Pathway analysis

Pathway analysis or gene set enrichment analysis was done using the prerank function from gseapy [87] on the t-statistics, computed as described in the ‘Differential gene expression’ section of the methods. To this end, the gene set library “KEGG_2019_Human” provided by enrichr [88] was used. Top pathways were selected using the normalized enrichment score as previously described [87].

#### Gene regulation

[48] The python implementation of the SCENIC (pySENIC) was used to infer regulons specific for CD4^+^ T helper cells. SCENIC infers a gene regulatory network using GRNBoost2 and creates co-expression modules. The co-expression modules get associated with transcription factors using the transcription factor motif discovery tool RcisTarget. A pair of transcription factor and associated gene set is called a regulon. For each cell, the regulons get scored using the AUCell algorithm to examine if a cell is affected by the regulon. We used default parameters of the pySENIC implementation.

#### Silhouette Score

[89] - is a measure to evaluate clustering performance by comparing the mean intra-cluster distance to the mean nearest-cluster distance. The Silhouette score is computed for batch and cell type labels on the scaled and PCA-transformed data using a varying number of principal components (interval [10, 50]). The score is defined in the intervall [−1,1], where a positive value indicates separated clusters, a value of zero signifies cluster overlap, and a negative value when the closest cluster is not the wrong cluster. For accessing batch mixing a low, close to zero, value is best, while for cell type clusters a value close to 1 is best. The scikit-learn implementation was used.

#### Adjusted Rand Index

[90] - The Rand index estimates the similarity between two clusterings by comparing all possible pairings of samples. The Adjusted Rand Index is adjusted for chance, such that a random labeling would result in a value close to 0, while a perfect clustering yields a score of 1. The Adjusted Rand Index is computed on the result of the leiden clustering algorithm using 20 different resolution parameters in the interval of [0.1, 30]. The best value (lowest for batch mixing, highest for cell type clustering) was used as the final score. The neighborhood graph for the leiden clustering algorithm is computed on scaled and PCA-transformed values, similar to the silhouette score, for a varying number of principal components (interval [10, 5]). The scikit-learn implementation was used.

#### Adjusted Mutual Information

[91] - Mutual Information measures the similarity between two clusterings by computing the sizes of the intersection of all possible cluster label pairs. The Adjusted Mutual Information is adjusted for chance, such that a random labeling would result in a value close to 0, while a perfect clustering yields a score of 1. Additionally, this accounts for the fact that Mutual Information is generally higher for clusterings with larger numbers of clusters. The AMI was computed on clustering results as described for the Adjusted Rand Index. The scikit-learn implementation was used.

#### COVID-19 classification

To evaluate the importance of the cell types found in the covid-blood-severity-hq dataset after reconstruction with DISCERN, the fraction for all T cell subtypes was used to predict the disease severity, as provided in [53]. The data was classified using a Gradient boosting classifier ([92], implemented in scikit-learn v1.0.2, default settings) using 25 rounds of leave-one-out cross-validation (LOOCV). Each round consists of n training-prediction iterations with *n* – 1 samples for training and 1 sample for testing, such that after one round prediction results for all n samples could be evaluated. We chose LOOCV over k-fold cross-validation and testing due to the limited size of the dataset, consisting of only 71 patients. We used pycm ([93], v3.3) for the performance evaluation. The final evaluation was done using the accuracy and F1 score as provided by pycm. The area under the receiver operating characteristic (AUROC) curve is computed with scikit-learn. Before training the classifiers a forward feature selection was performed using the SequentialFeatureSelector implemented in scikit-learn with default parameters. In total four experiments were performed. In the first experiment, classification with three disease categories (mild, moderate, severe) was used. Patients who died were excluded. For the other two experiments only patients with asymptomatic, mild, severe and critical symptoms were included. In all experiments the asymptomatic and mild category was merged to mild and severe and critical to severe.

