## Supplementary Material for "DiSCERN - Deep Single Cell Expression ReconstructioN for improved cell clustering and cell subtype and state detection"

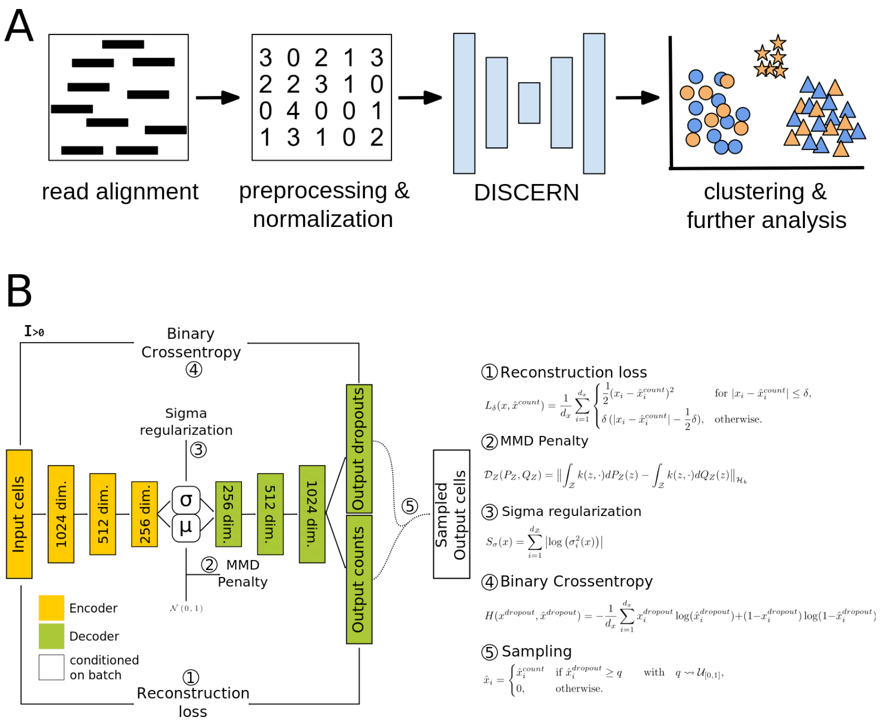


Figure S1: *DISCERN workflow and method details.* **A**: A standard scRNA-seq workflow starts by aligning the sequencing reads to a reference transcriptome to obtain a cell-by-gene count table, which is subsequently preprocessed, filtered and normalized. The normalized count matrices are then used for downstream analyses, such as clustering, differential expression between clusters, and marker gene identification. When combining multiple datasets an alignment or batch correction step is commonly performed to reduce differences between the datasets. DISCERN is used after the preprocessing and normalization steps to integrate the high quality and low quality datasets, reconstructing the gene counts of the low quality to that of the high quality dataset (or vice versa). DISCERN is able to correct for batch effects, provides a lower dimensional representation, and a corrected expression matrix. This corrected expression matrix can directly be used for downstream analysis or used with clustering algorithms. **B**: Overview of the DISCERN neural network architecture consisting of a random encoder (yellow) and a deterministic decoder (green) which can be conditioned on the batch information. DISCERN's loss function contains a (1) count fitting reconstruction loss, (2) a prior fitting MMD-penalty, (3) a sigma regulation term as to prevent the random encoder to collapse to a deterministic one, and (4) a binary cross-entropy term for learning the probability of a dropout event. The final output is generated by sampling from the estimated counts with the estimated dropout probabilities using formula (5).

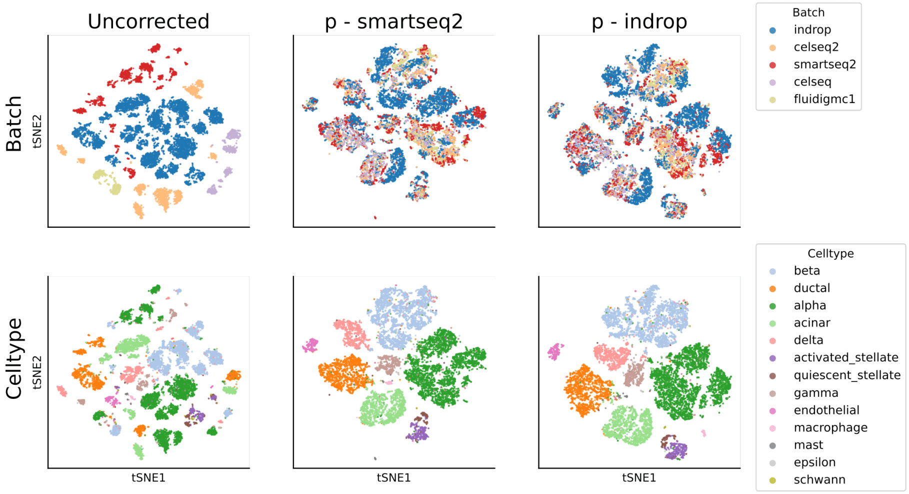


Figure S2: *t-SNE visualization of the pancreas dataset before reconstruction (Uncorrected) and after reconstruction with DISCERN.* The first row is colored by the origin of the dataset (batch) and the second is colored by the cell type annotations. Both batch and cell type annotations were taken from the published dataset. For DISCERN, two projections to the hq smartseq2 batch (second column) and to the lq indrop batch (third column) are shown.

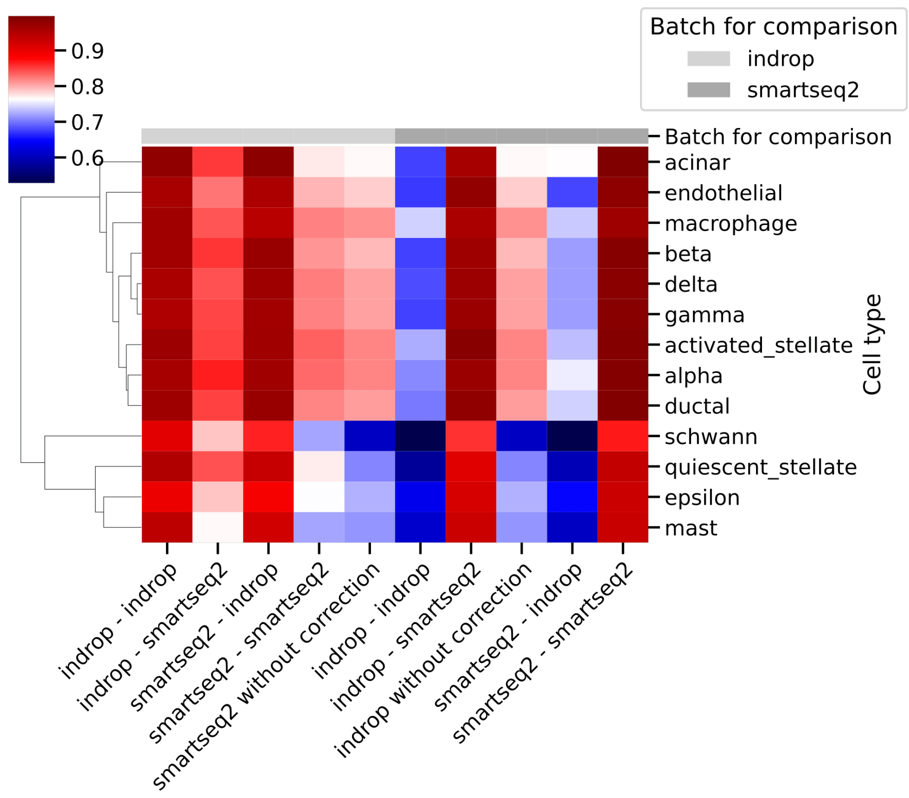


Figure S3: *Heatmap showing the Pearson correlation of the average gene expression per celltype (rows) for the pancreas dataset.* The starting datasets and target dataset for correction are listed on the x-axis. The second entry, for instance, signifies that an indrop dataset was projected to a smartseq2 dataset using DISCERN's expression reconstruction. The correlation is computed between the batch shown in the top row (light gray = indrop, dark gray = smartseq2) and the expression-reconstructed data as listed on the x-axis.

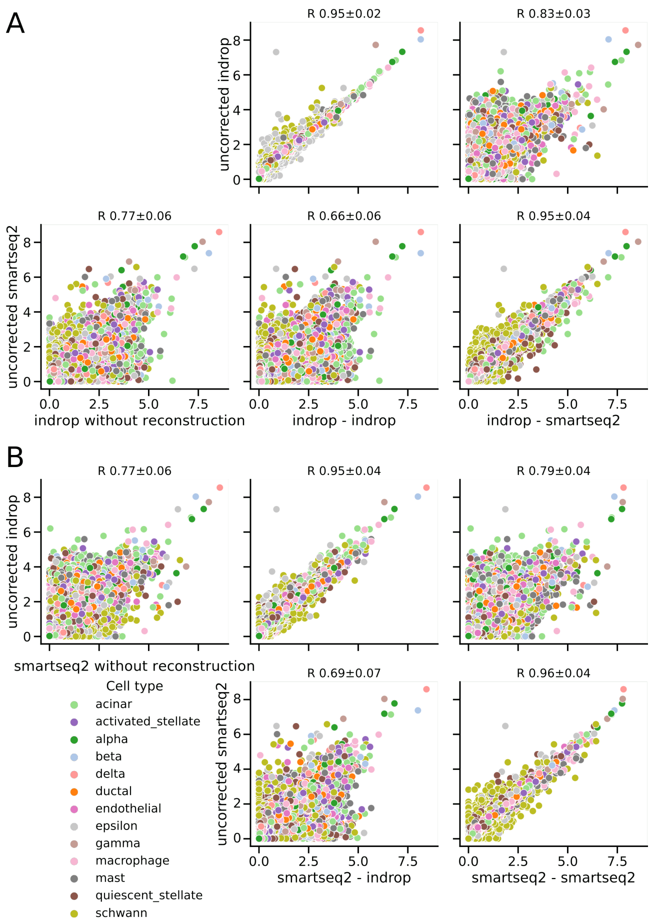


Figure S4: *Average gene expression of the pancreas dataset.* The figures are organized in three columns extended over A and B indicating before DISCERN reconstruction (first column) and after reconstruction using DISCERN (second and third column) stratified by cell type (color). The average expression is compared to the average gene expression of only the indrop (upper row) or the smartseq2 data (lower row). The dataset that is used for projection with DISCERN is shown at the x-axis of each plot after "-", e. g. "smartseq2 - indrop" means smartseq2 projected to indrop. Each colored dot represents one gene. The mean Pearson correlation with one standard deviation over all cell types is displayed in the figure title. **A:** Reconstruction of the indrop batch. **B:** Reconstruction of the smartseq2 batch.

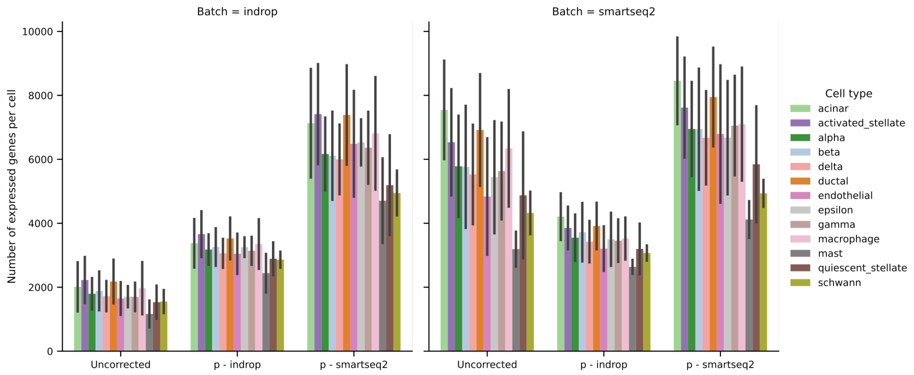


Figure S5: *Number of expressed genes in the indrop (left panel) and the smartseq2 data (right panel) of the pancreas dataset before (Uncorrected) and after projection using DISCERN stratified by cell type (color).* In the right panel, p-indrop displays the gene expression per cell after smartseq2 data was projected to indrop data using DISCERN. Bar heights indicate the average number of expressed genes per cell type and batch. Error bars indicate one standard deviation of the mean over cells in the corresponding batch and cell type.

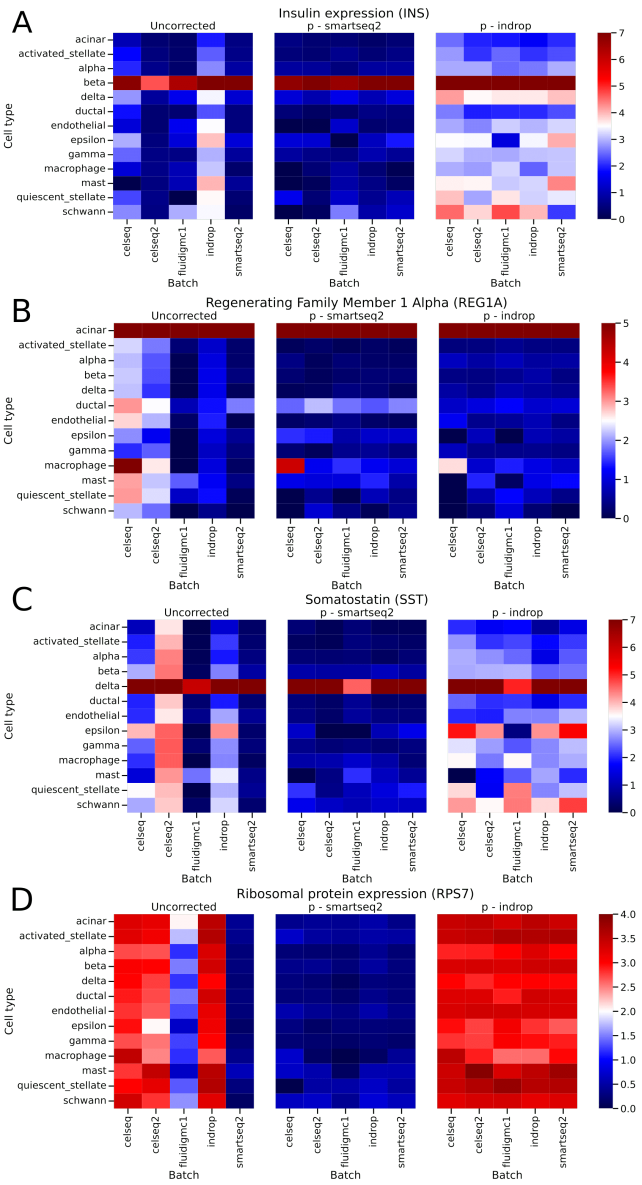
A
Insulin expression (INS)
Uncorrected
p - smartseq2
p - indrop
acinar
activated_stellate
alpha
beta
delta
ductal
endothelial
epsilon
gamma
macrophage
mast
quiescent_stellate
schwann
Cell type
celseq
celseq2
fluidigmc1
indrop
smartseq2
Batch
0
1
2
3
4
5
6
7
B
Regenerating Family Member 1 Alpha (REG1A)
Uncorrected
p - smartseq2
p - indrop
Cell type
Batch
0
1
2
3
4
5
C
Somatostatin (SST)
Uncorrected
p - smartseq2
p - indrop
Cell type
Batch
0
1
2
3
4
5
6
7
D
Ribosomal protein expression (RPS7)
Uncorrected
p - smartseq2
p - indrop
Cell type
Batch
0.0
0.5
1.0
1.5
2.0
2.5
3.0
3.5
4.0

Figure S6: *Average gene expression of Insulin (INS), a ribosomal gene (RPS7), REG1A, and Somatostatin (SST) by cell type (rows) and by batch (columns) in the pancreas dataset.* The first column shows the uncorrected datasets, while the second and third column show projections using DISCERN to the smartseq2 and the indrop dataset, respectively. **A:** *INS* was selected because it is a cell type-determining gene for beta cells and *RPS7* is known to be expressed in nearly all cells. While nearly all batches display exclusive *INS* expression in beta cells in uncorrected data, the indrop data shows a more dispersed expression of *INS* in several cell types. Projection to the smartseq2 batch results in a beta cell-specific expression in the corrected indrop data (second column). Projection to the indrop batch results in dispersed INS expression for all batches (third column). **B:** *REG1A* is a acinar cell specific gene, shown to be involved in acinar cell carcinoma [1]. For most pancreatic datasets, it is exclusively expressed in acinar cells in the uncorrected data. Only celseq shows a more dispersed expression across several cell types. After reconstruction to indtop or smartseq2 data the expression of *REG1A* is restricted to acinar cells and macrophages in the celseq batch. **C:** *SST* is known to be produced by delta cells in the pancreas [2], which can be observed for instance in the smartseq2 batch. After reconstruction to the smartseq2 batch delta cell-specific expression of *SST* is observed for all datasets. **D:** *RPS7* shows high expression in the indrop, celseq and the celseq2 batch, whereas smartseq2 and fluidigmc1 show low to no expression, as described previously [3]. This expression of *RPS7* can be removed by projecting to smartseq2 or reconstructed by projection to indrop data.

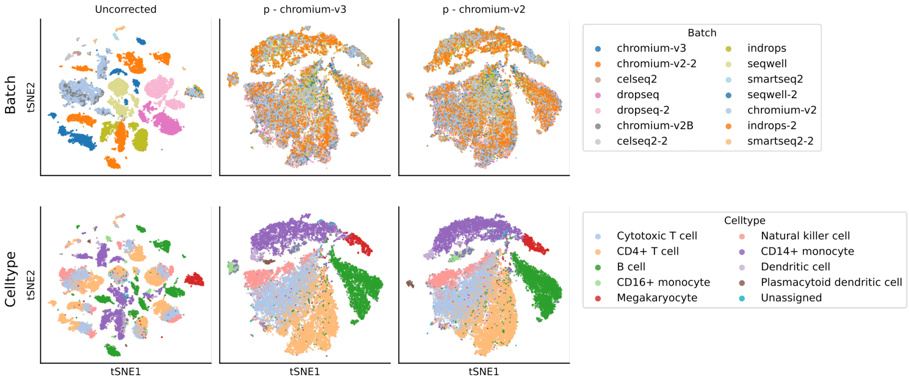


Figure S7: *t-SNE visualization of the difftec dataset before reconstruction (Uncorrected) and after reconstruction with DISCERN.* The first row shows the dataset of origin (batch) and the second row shows the cell type annotations which are available together with the dataset. For DISCERN two projections, one to the hq chromium-v3 batch and one to the lq chromium-v2 batch is shown.

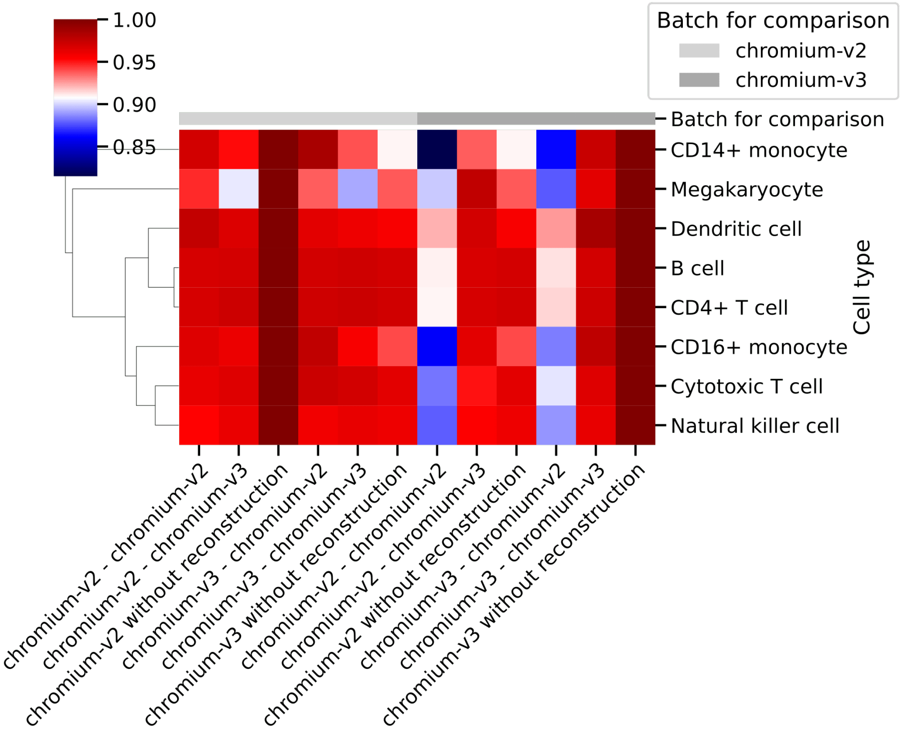


Figure S8: *Heatmap showing the Pearson correlation of the average gene expression per celltype (rows) for the difftec dataset.* The starting datasets and target dataset for correction are listed on the x-axis. The second entry, for instance, signifies that a chromium-v2 dataset was projected to a chromium-v3 batch using DISCERN's expression reconstruction. The correlation is computed between the batch shown in the top row (light gray = chromium-v2, dark gray = chromium-v3) and the expression-reconstructed data as listed on the x-axis.

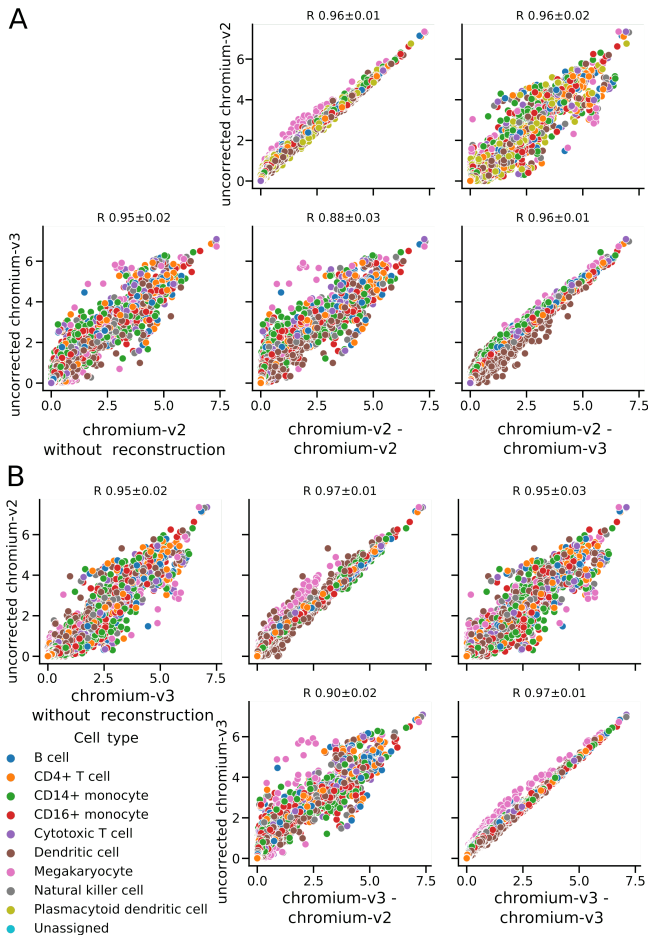


Figure S9: *Average gene expression of the difftec dataset.* The figures are organized in three columns extended over A and B indicating before DISCERN reconstruction (first column) and after reconstruction using DISCERN (second and third column) stratified by cell type (color). The average expression is compared to the average gene expression of only the chromium-v2 (upper row) or the chromium-v3 data (lower row). The dataset that is used for projection with DISCERN is shown at the x-axis of each plot after "-", e. g. "chromium-v2 - chromium-v3" signifies chromium-v2 data was projected to the chromium-v3 batch. Each colored dot represents one gene. Colors indicate the cell type identity. The mean Pearson correlation with one standard deviation over all cell types is displayed in the figure title. **A:** Reconstruction of the chromium-v2 batch. **B:** Reconstruction of the chromium-v3 batch.

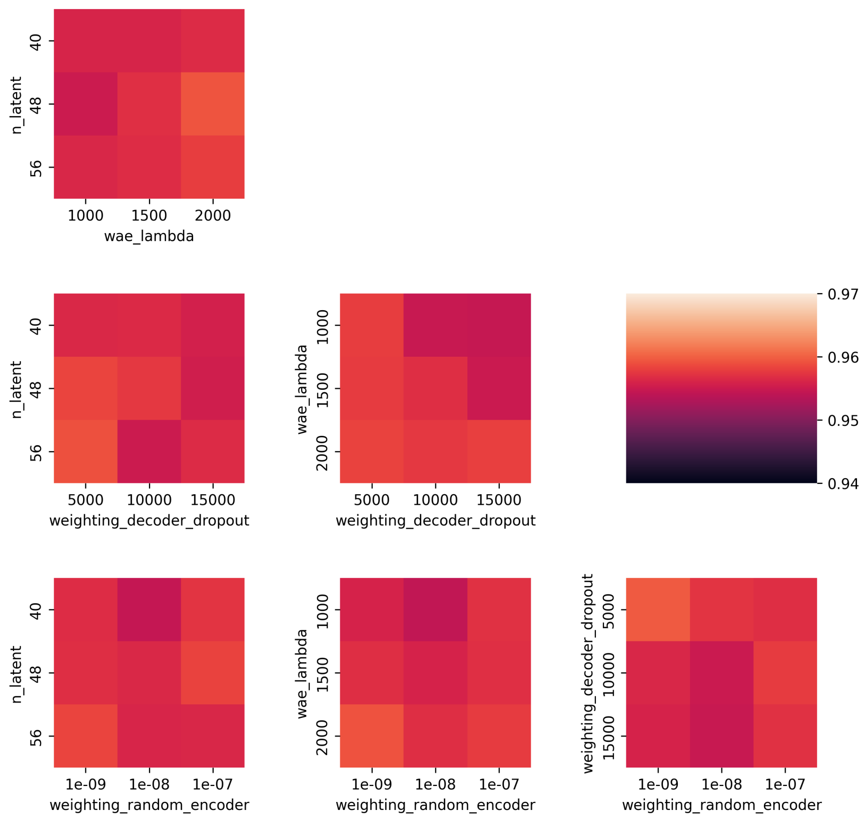


Figure S10: *Average gene expression correlation in the pancreas dataset for multiple combinations of the four main hyperparameters of DISCERN.* Pearson correlation was computed on the average gene expression per cell type between the uncorrected smartseq2-hq batch and all other batches projected to the smartseq2-hq batch. All possible combinations between the four hyperparameters were trained. Each box in the heatmaps represent the average of all models sharing this combination of hyperparameter. n_latent specifies the number of latent dimensions (default 48), wae_lambda is a scaling factor for the MMD penalty on the latent dimensions and the prior (default 1500), weighting_random_encoder is a scaling factor of the loss on the estimated variance in the latent dimensions (default $1 \times 10^{-8}$) as described in [4] and the weighting_decoder_dropout is the scaling factor of the cross entropy loss of the dropout estimation (default 10 000).

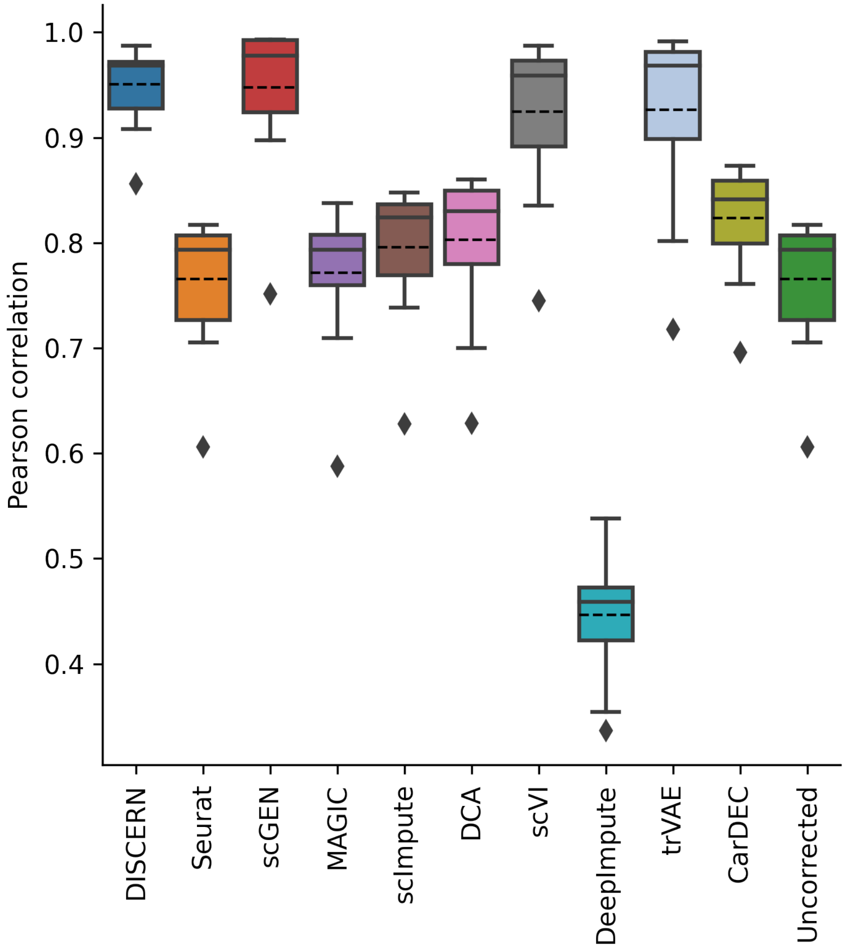


Figure S11: *Pearson correlation of the average gene expression in the pancreas dataset comparing the smartseq-hq and reconstructed indrop-lq batch.* The average gene expression was calculated per cell type and gene and correlated per cell type. For DISCERN, scGEN, scVI and trVAE the projection to smartseq-hq is shown. Boxplots represent median, quantiles, minimum, maximum, and potential outliers. The dotted line shows the arithmetic mean.

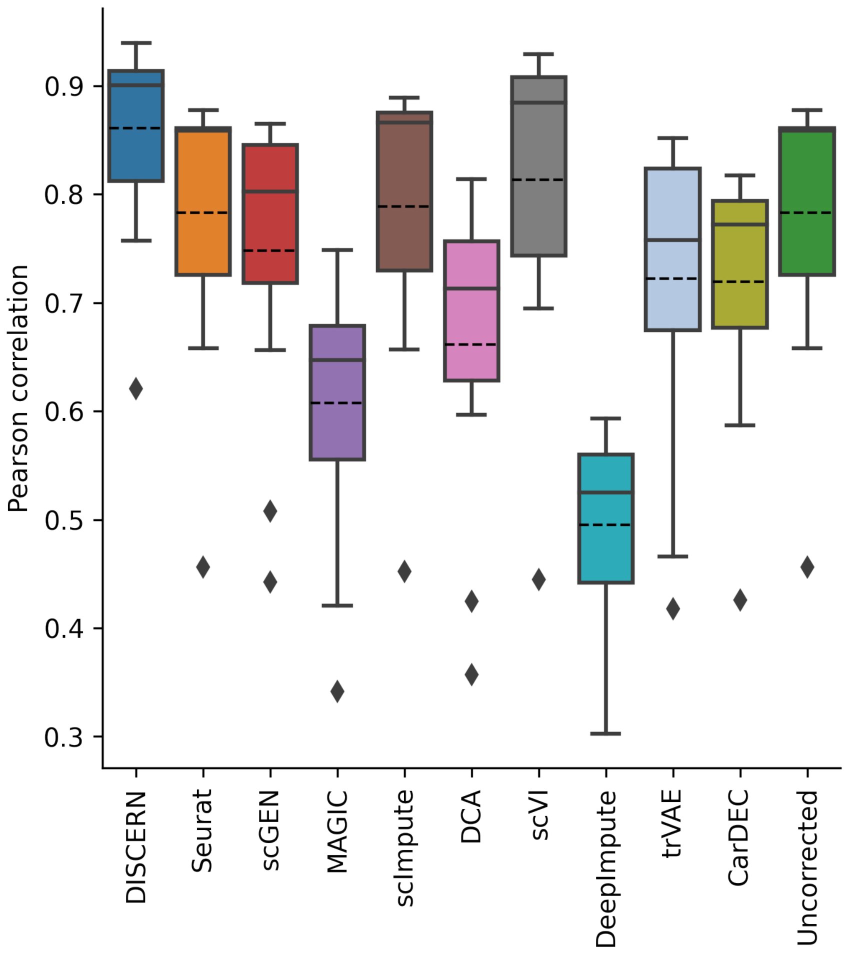


Figure S12: *Pearson correlation of the standard deviation of gene expression in the pancreas dataset comparing the smartseq-hq and reconstructed indrop-lq batch.* The standard deviation was calculated per cell type and gene and correlated per cell type. For DISCERN, scGEN, scVI and trVAE the projection to smartseq-hq is shown. Boxplots represent median, quantiles, minimum, maximum, and potential outliers. The dotted line shows the arithmetic mean.

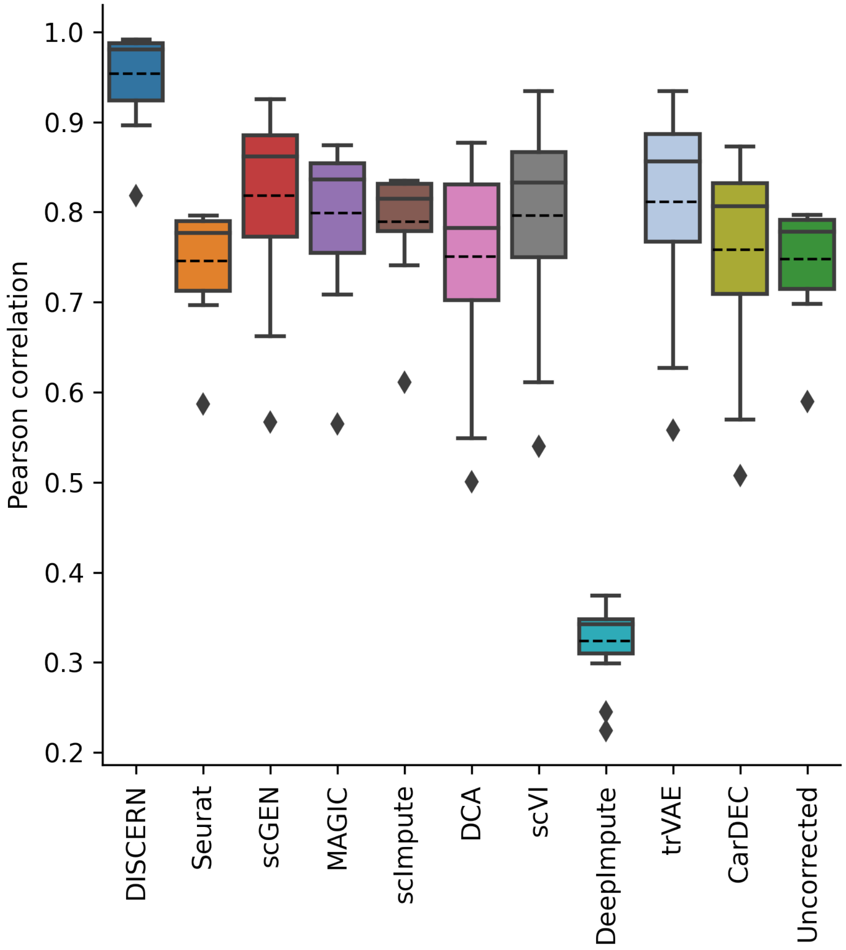


Figure S13: *Pearson correlation of the dropout rate of gene expression in the pancreas dataset comparing the smartseq-hq and reconstructed indrop-lq batch.* The dropout rate was calculated per cell type and gene and correlated per cell type. A gene with a log-normalized expression below 0.1 was treated as dropout. For DISCERN, scGEN, scVI and trVAE the projection to smartseq-hq is shown. Boxplots represent median, quantiles, minimum, maximum, and potential outliers. The dotted line shows the arithmetic mean.

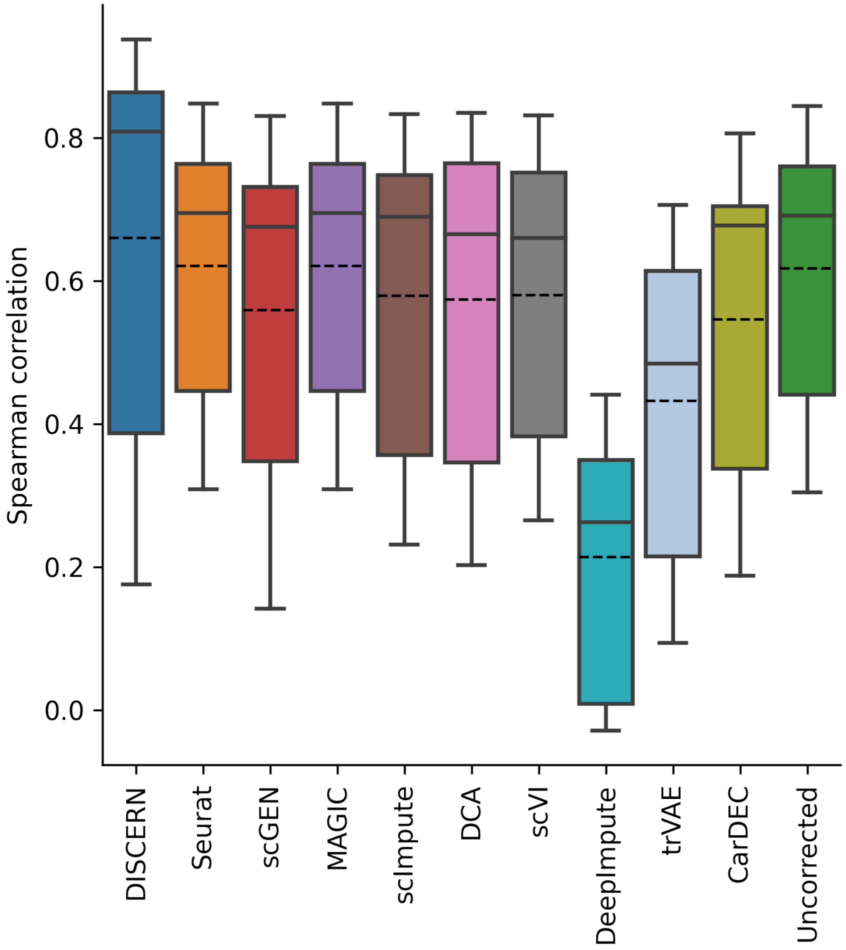


Figure S14: *Spearman correlation of the t-statistics in one vs rest differential expression analysis in the pancreas dataset comparing the smartseq-hq and reconstructed indrop-lq batch.* The t-statistics was calculated per cell type and gene and correlated per cell type. For DISCERN, scGEN, scVI and trVAE the projection to smartseq-hq is shown. All available genes were used for calculating the correlation. Boxplots represent median, quantiles, minimum, maximum, and potential outliers.

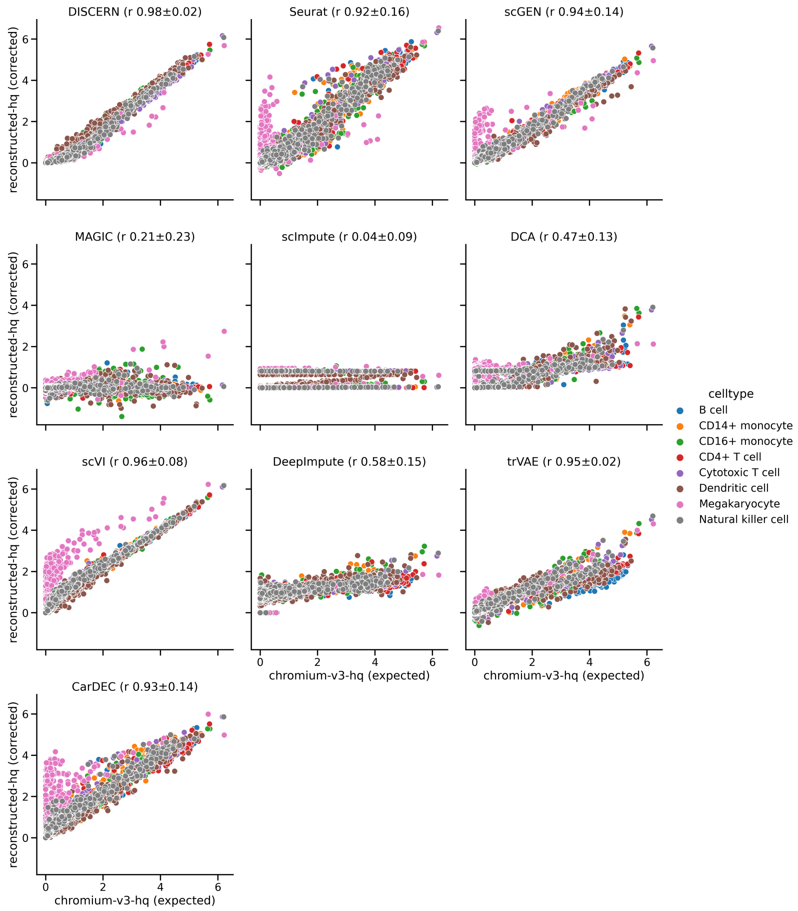


Figure S15: *Comparison of the average gene expression reconstruction performance for several methods for the difftec dataset.* Four imputation (DCA, MAGIC, scImpute, DeepImpute), four batch correction methods (Seurat, scGEN, scVI, trVAE), and DISCERN are compared. The dataset is based on the difftec dataset where the chromium-v3 batch was split into chromium-v3-lq and chromium-v3-hq and selected genes were removed (in silico gene drop out) from chromium-v3-lq. The corrected average gene expression (y-axis) is based on the reconstructed or imputed chromium-v3-lq data. For DISCERN, scGEN, scVI, and trVAE the projection onto the chromium-v3-hq reference is depicted. The expected average gene expression (x-axis) is based on the unmodified chromium-v3-lq batch. Mean Pearson correlation with one standard deviation over all cell types is displayed in parentheses of the figure title. Colors indicate the cell type identity.

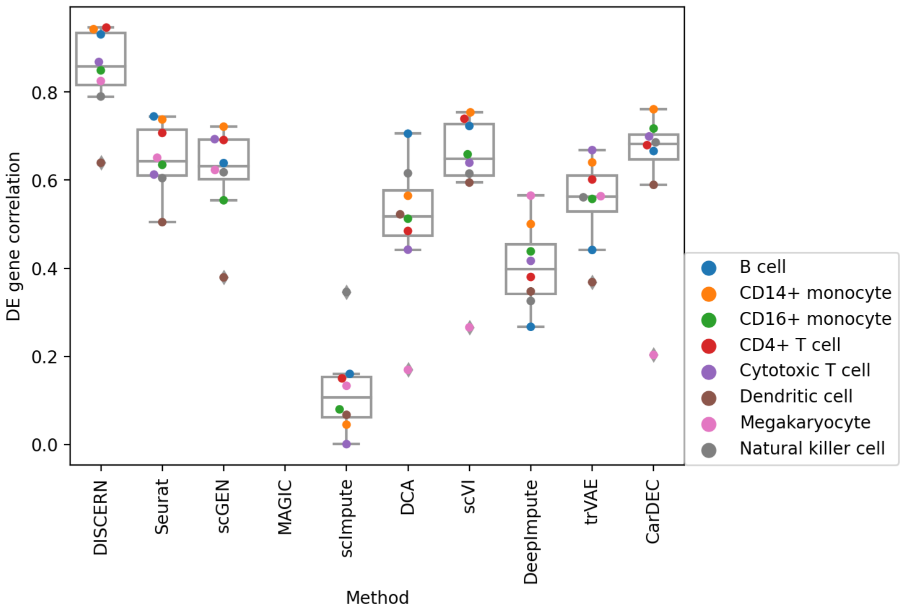


Figure S16: *Pearson correlation of DEG t-statistics for a one-vs-rest cell type comparison and in silico gene removal.* The dataset is based on the difftec dataset where the chromium-v3 batch was split into chromium-v3-lq and chromium-v3-hq and selected genes were removed from chromium-v3-lq data. The corrected average gene expression is based on reconstructed or imputed chromium-v3-lq only, while the expected average gene expression is based on the unmodified chromium-v3-lq batch. For DISCERN, scGEN, scVI, and trVAE the projection to chromium-v3-hq is shown. Only genes, which were removed in chromium-v3-lq were used for calculating the correlation. Boxplots represent median, quantiles, minimum, maximum, and potential outliers. Colors indicate the cell type identity.

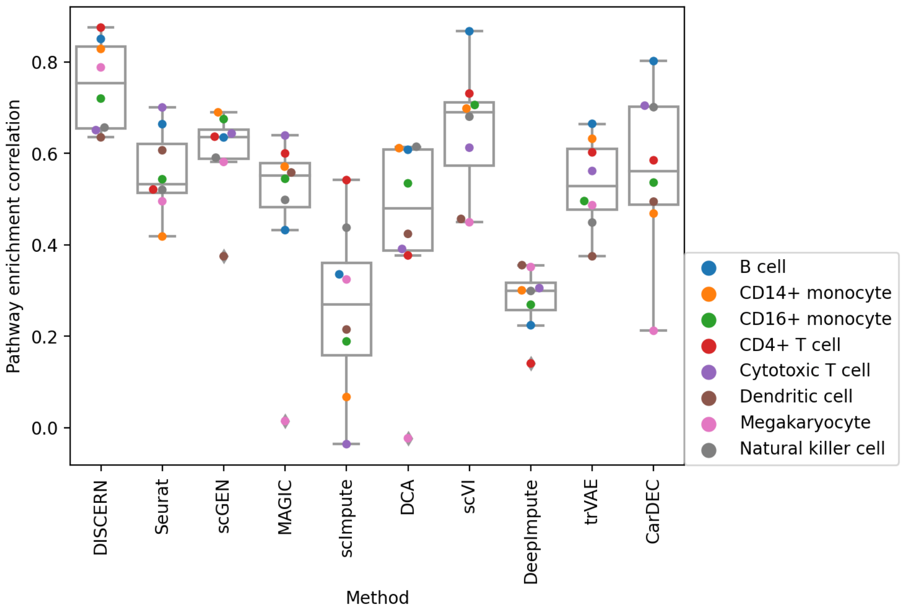


Figure S17: *Pearson correlation of KEGG gene set enrichment scores for a one-vs-rest cell type comparison and in silico gene removal.* The dataset is based on the difftec dataset where the chromium-v3 batch was split into chromium-v3-lq and chromium-v3-hq and selected genes were removed from chromium-v3-lq data. Instead of directly measuring DEG correlation as in Figure S16 a gene set enrichment analysis was performed for DEGs and correlated to the ground-truth 'expected' information. The corrected average gene expression is based on reconstructed or imputed chromium-v3-lq only, while the expected average gene expression is based on the unmodified chromium-v3-lq batch. For DISCERN and scGEN the projection to chromium-v3-hq is shown. Boxplots represent median, quantiles, minimum, maximum, and potential outliers. Colors indicate the cell type identity.

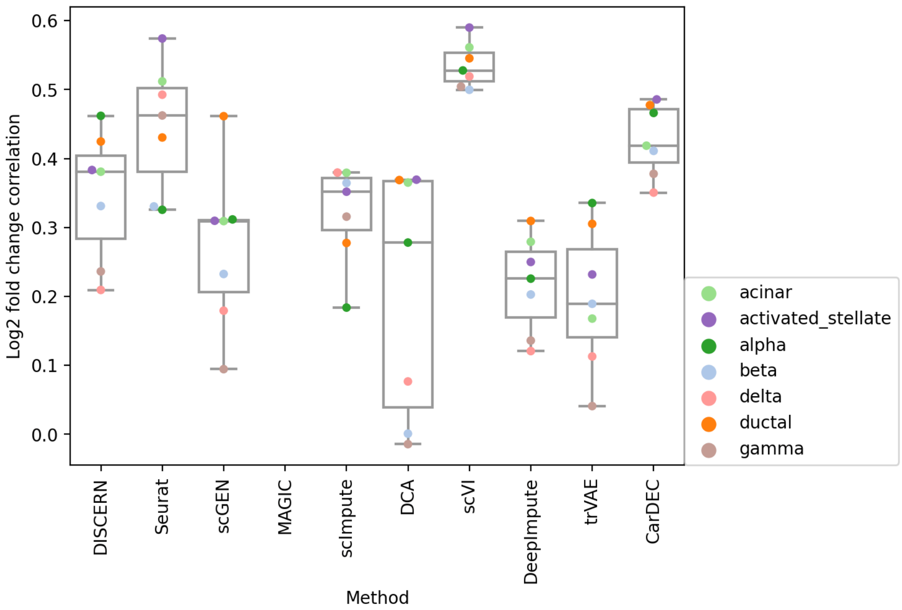


Figure S18: *Pearson correlation of the log2 fold-change (FC) for a one-vs-rest cell type comparison and in silico gene removal. The dataset is based on the smartseq2 dataset where the smartseq2 batch was split into smartseq2-lq and smartseq2-hq and selected genes were removed from smartseq2-lq data. For each cell type, DEG and FC were calculated against all other cell types. For DISCERN, scGEN, scVI, and trVAE* the projection of indrop-lq to smartseq2-hq data is shown, resulting in reconstructed-hq data. Only genes, which were removed in smartseq2-lq, were used for calculating the correlation. Boxplots represent median, quantiles, minimum, maximum, and potential outliers. Colors indicate the cell type identity.

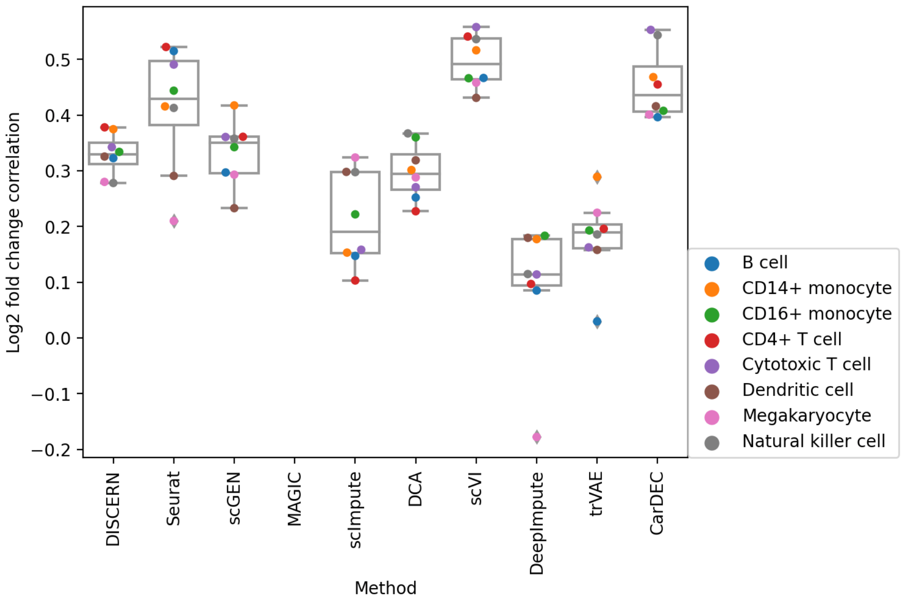


Figure S19: *Pearson correlation of the log2 fold-change (FC) for a one-vs-rest cell type comparison and in silico gene removal.* The dataset is based on the difftec dataset where the chromium-v3 batch was split into chromium-v3-lq and chromium-v3-hq and selected genes were removed from chromium-v3-lq data. For each cell type, DEG and fold-change were calculated against all other cell types. For DISCERN, scGEN, scVI, and trVAE the projection of chromium-v2-lq to chromium-v3-hq data is shown, resulting in reconstructed-hq data. Only genes, which were removed in chromium-v3-lq, were used for calculating the correlation. Boxplots represent median, quantiles, minimum, maximum, and potential outliers. Colors indicate the cell type identity.

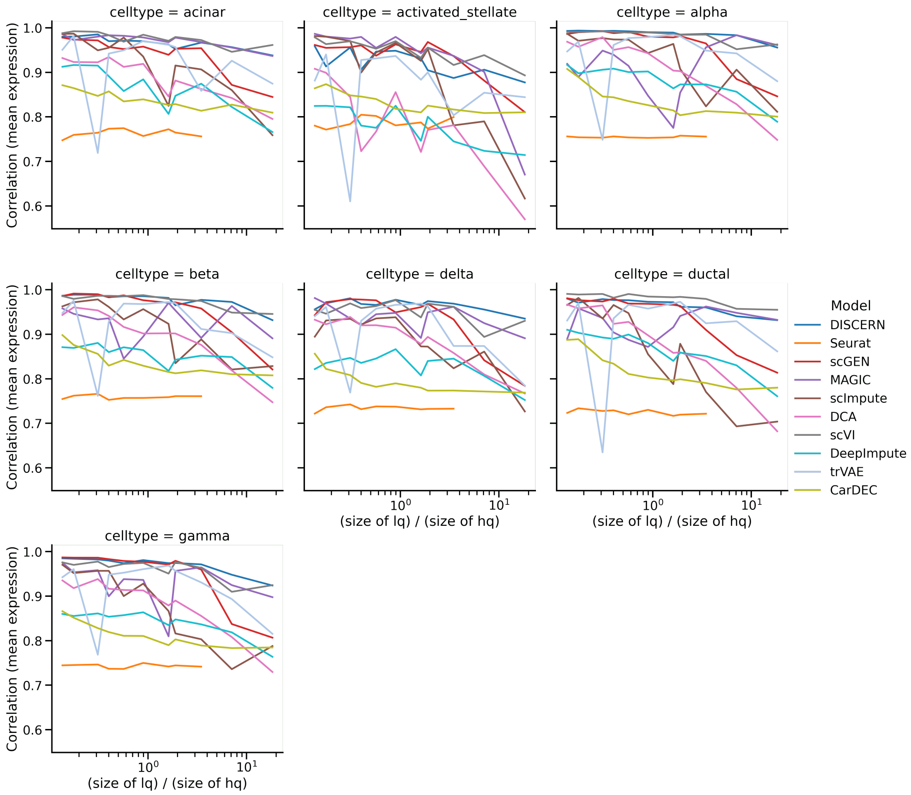


Figure S20: *Pearson correlation of the mean gene expression for the pancreas reconstructed-hq and smartseq2-hq data for different ratios of lq to hq training data.* The plot shows the dependency of the mean gene expression reconstruction on the ratio of lq to hq training data, showing increased performance for lower ratios and a marked decrease in performance for higher ratios, especially for scGen, while DISCERN remains relatively stable for all ratios tested. For DISCERN, scGEN, scVI, and trVAE the projection to smartseq2-hq is shown. Colors indicate different methods.

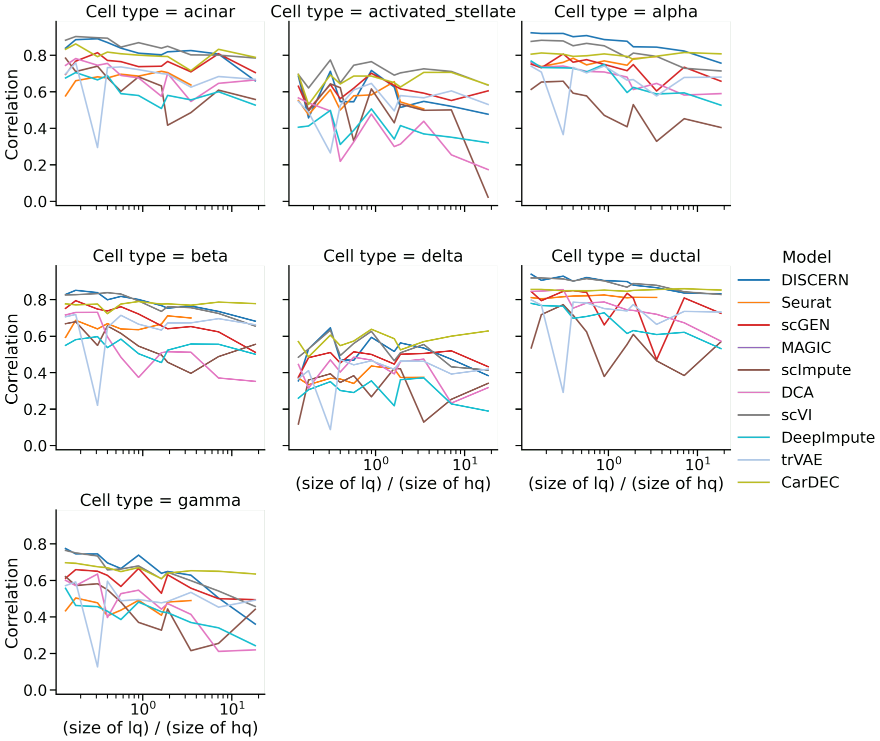


Figure S21: *Pearson correlation of DEG t-statistics for a one-vs-rest cell type comparison and in silico gene removal.* The dataset is based on the pancreas dataset where the smartseq2 batch was split into smartseq2-lq and smartseq2-hq and selected genes were removed from smartseq2-lq. The t-statistic is computed on removed genes after reconstruction of the smartseq2-lq batch to reconstructed-hq data and compared to the t-statistic of the unmodified smartseq2-lq batch. For DISCERN, scGEN, scVI, and trVAE the projection to smartseq2-hq is shown. Uncorrected, MAGIC, and CarDEC corrected data have close to zero gene expression in the smartseq2-lq for the selected genes and thus cannot be shown. Colors indicate different methods. Only genes, which were removed in smartseq2-lq, were used for calculating the correlation.

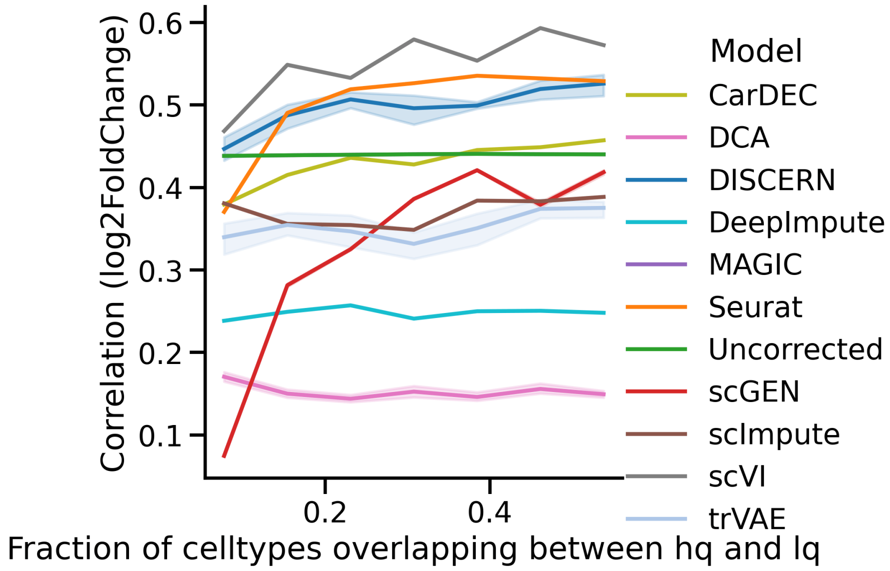


Figure S22: *Spearman correlation of the log2 fold-change (FC) of alpha cells that were reconstructed and ground-truth alpha cells that were excluded from training using pancreas data.* Different fractions of cell type overlap in the indrop-lq and smartseq2-hq training data were used to estimate the reconstruction performance when datasets become dissimilar. Alpha cells were only present in the indrop-lq data and smartseq2-hq alpha cells were extracted as ground truth information. X-axis shows fractions of cell types, which are non-alpha cells and overlap between lq and hq batches. Confidence intervals indicate the standard deviations from five independently trained models. The violet line for MAGIC and the light green line for CarDEC are not visible as the correlation is the same as achieved for uncorrected data. Best performance is observed for DISCERN and Seurat. All available genes were used for calculating the correlation.

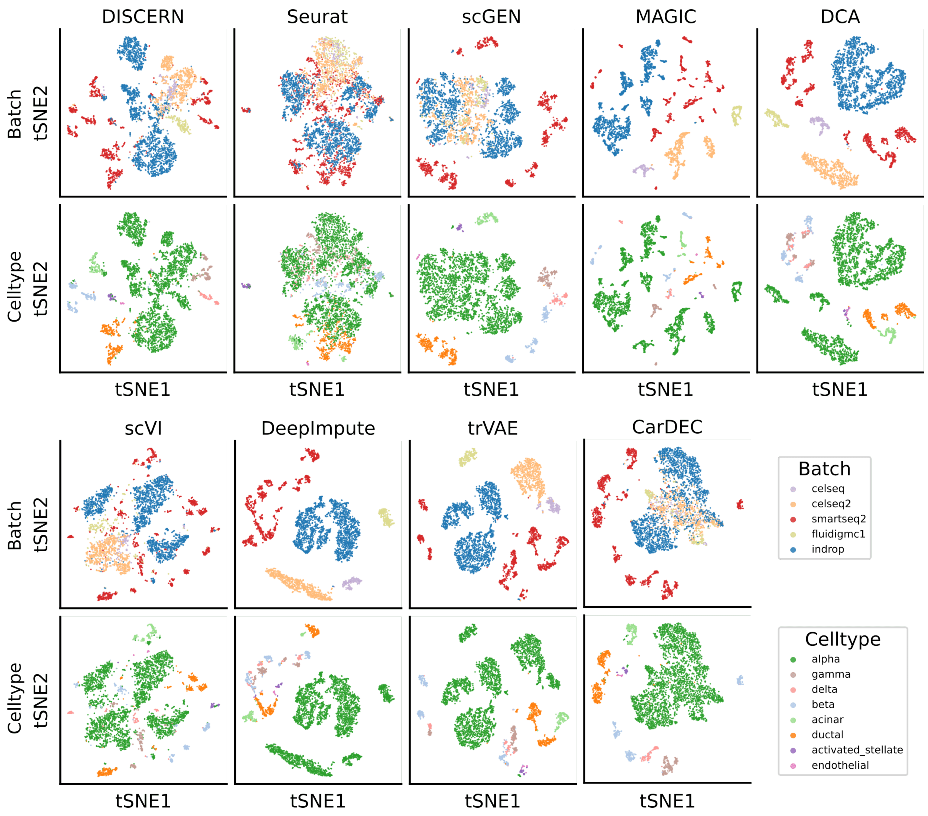


Figure S23: *t-SNE visualization of the pancreas dataset where alpha cells are removed from the smartseq2-hq batch and all cell types except alpha cells are removed from all other batches.* This means that there is no cell type overlap between the smartseq2 and the other batches. The columns show the data before reconstruction (Uncorrected) and the corrected dataset using DISCERN, Seurat, scGEN, MAGIC, DCA, scVI, DeepImpute, trVAE, and CarDEC. The cells are colored by batch (upper row) or by cell type (lower row). For DISCERN, scGEN, scVI, and trVAE the dataset was projected to the smartseq2 batch.

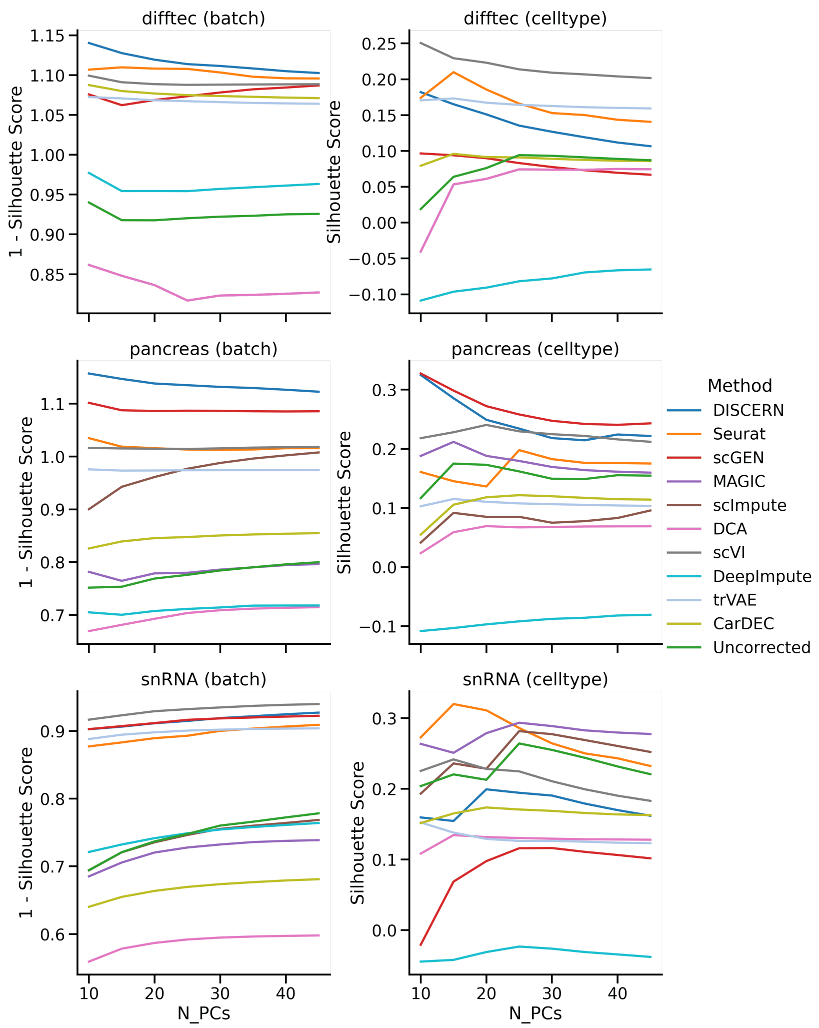


Figure S24: *Silhouette scores for batch and cell type clusters on the difftec, the pancreas, and the snRNA-seq datasets for varying numbers of principal components.* The first column measures batch clustering (1 - Silhouette score, higher is better) and the second column cell type clustering (higher is better). Varying numbers of principal components were selected because they influence downstream applications, for example clustering, and the models show their best metrics at different numbers of components.

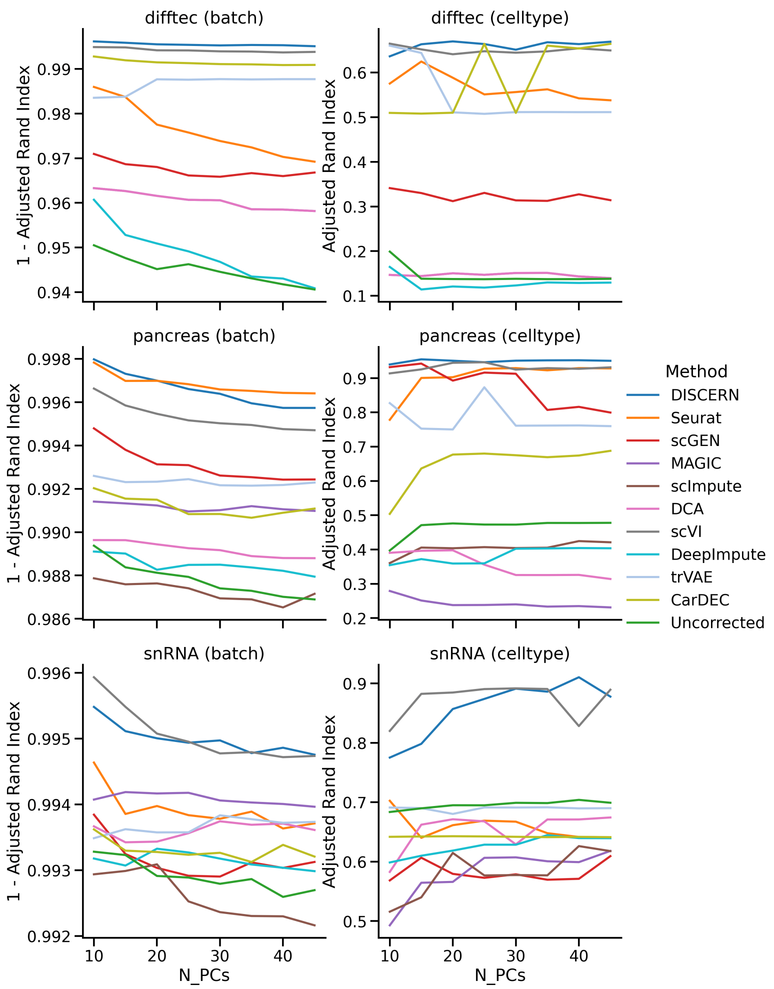


Figure S25: *Adjusted Rand Index (ARI) for batch and cell type clusters on the difftec, the pancreas, and the snRNA-seq dataset for varying numbers of principal components.* The first column measures batch clustering (1 - ARI, higher is better) and the second column cell type clustering (higher is better). Varying numbers of principal components were selected because they influence downstream applications, for example clustering, and the models show their best metrics at different numbers of components. Clustering was performed using the Leiden algorithm on 20 different resolutions per number of components. The best ARI (lowest for the batch, highest for the cell type metric) for each number of components is displayed.

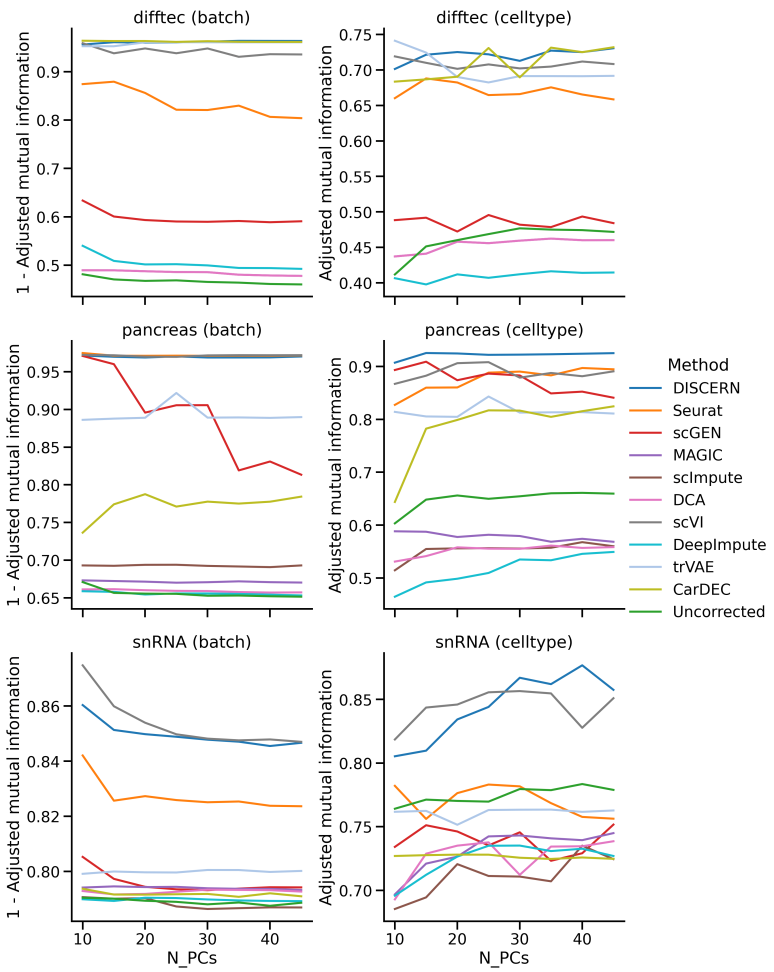


Figure S26: *Adjusted Mutual Information (AMI) for batch and cell type clusters on the difftec, the pancreas and the snRNA-seq dataset for varying numbers of principal components.* The first column measures batch clustering (1 - AMI, higher is better) and the second column cell type clustering (higher is better). Varying numbers of principal components were selected because they influence downstream applications, for example clustering, and the models show their best metrics at different numbers of components. Clustering was performed using the Leiden algorithm on 20 different resolutions per number of components. The best AMI (lowest for the batch, highest for the cell type metric) for each number of components is displayed.

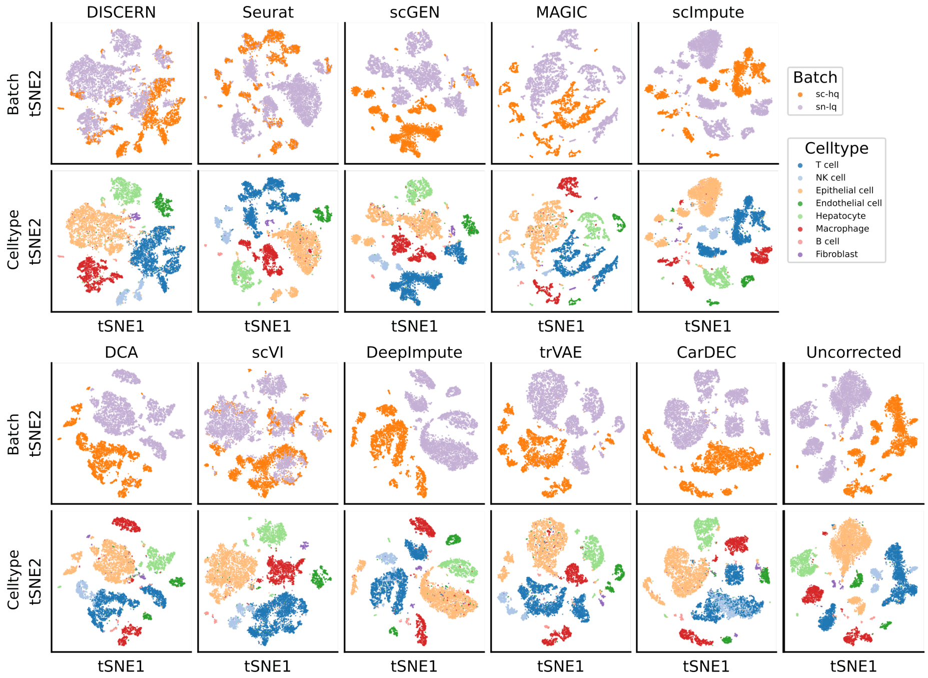


Figure S27: *t-SNE visualization of scRNA-seq and snRNA-seq data before (Uncorrected) and after reconstruction with DISCERN, Seurat, scGEN, scVI, DeepImpute, trVAE, and CarDEC.* In this dataset the same sample from a metastatic liver biopsy was sequenced using scRNA-seq and snRNA-seq technology, yielding the sc-hq and sn-lq datasets. The sn-lq data was reconstructed using the sc-hq reference. The first row shows the color annotation by batch and the second row is colored by the different cell types found in the dataset. Annotation of the cell types was provided with the original data.

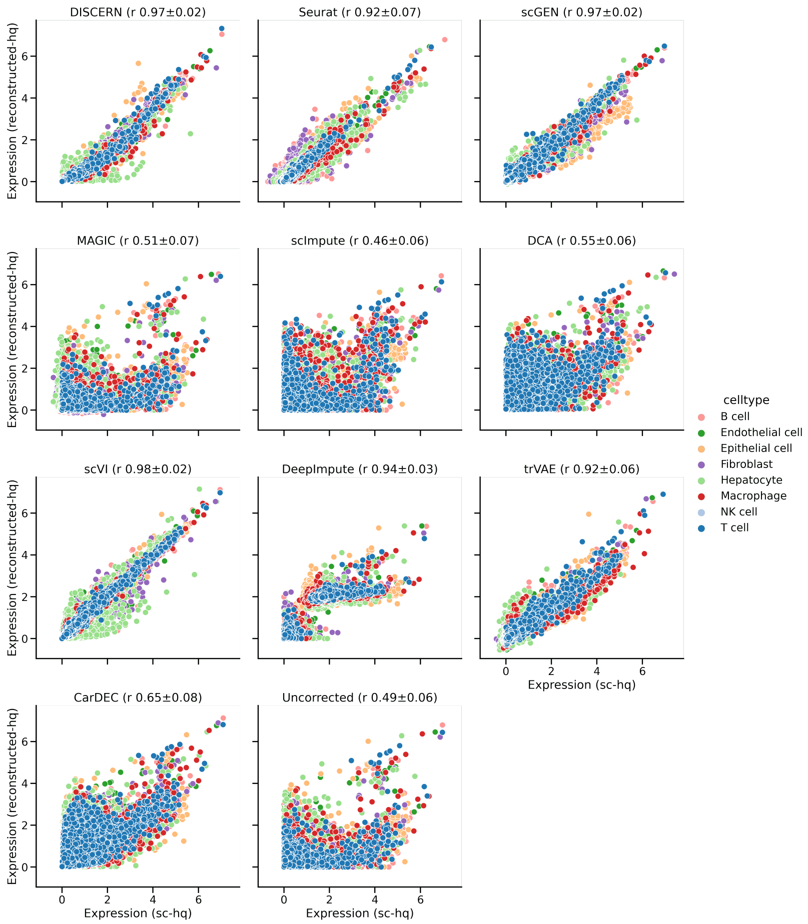


Figure S28: *Average gene expression of scRNA-seq and snRNA-seq data before (Uncorrected) and after reconstruction with DISCERN, Seurat, scGEN,MAGIC, scImpute, DCA, scVI, DeepImpute, trVAE, and CarDEC.* In this dataset the same sample from a metastatic liver biopsy was sequenced using scRNA-seq and snRNA-seq technology. The sn-lq data was reconstructed using the sc-hq reference to yield reconstructed-hq data. Each colored dot represents one gene. Colors indicate the cell type identity. The mean Pearson correlation with one standard deviation over all cell types is displayed in the figure title.

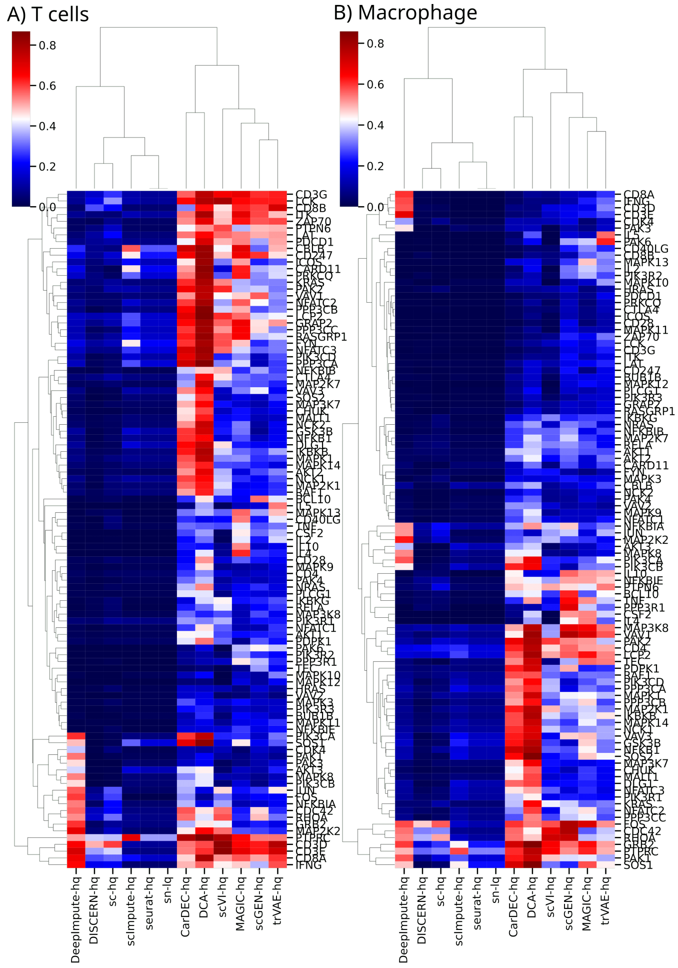


Figure S29: *Average gene expression of T cell receptor signaling genes in T cells (**A**) and Macrophages (**B**).* The columns show the data in the snRNA-seq (before reconstruction, sn-lq) and snRNA-seq dataset after reconstruction with DISCERN, Seurat, DeepImpute, scImpute, CarDEC, scGEN, scVI, DCA, MAGIC, and trVAE and in the scRNA-seq data (sc-hq). The average expression was min-max scaled with adding a pseudocount of $1 \times 10^{-3}$. The reconstructed-hq shows high similarity with the expression in the sc-hq dataset. Only genes with a maximum expression greater than 0.2 are shown.

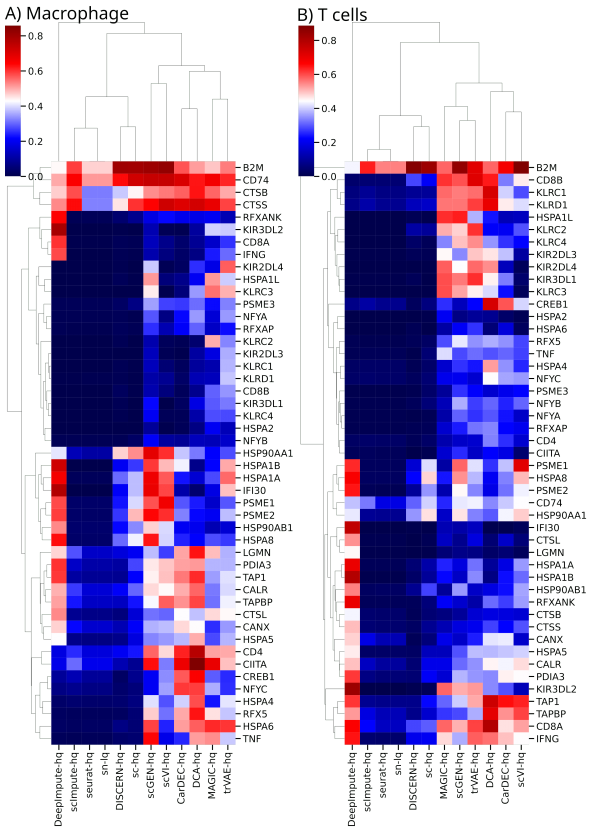


Figure S30: *Average gene expression of antigen presentation and processing genes in Macrophages (**A**) and T cells (**B**).* The columns show the data in the snRNA-seq (before reconstruction, sn-lq) and snRNA-seq dataset after reconstruction with DISCERN, Seurat, DeepImpute, scImpute, CarDEC, scGEN, scVI, DCA, MAGIC, and trVAE and in the scRNA-seq data (sc-hq). The average expression was min-max scaled with adding a pseudocount of $1 \times 10^{-3}$. reconstructed-hq shows high similarity with the expression in the sc-hq dataset. Only genes with a maximum expression greater than 0.2 are shown. *CD4* and *CD8A* genes are part of the antigen presentation and processing pathway (`https://www.genome.jp/pathway/hsa04612`) but are naturally not expressed in Macrophages, thus no expression of these genes is expected in **A**.

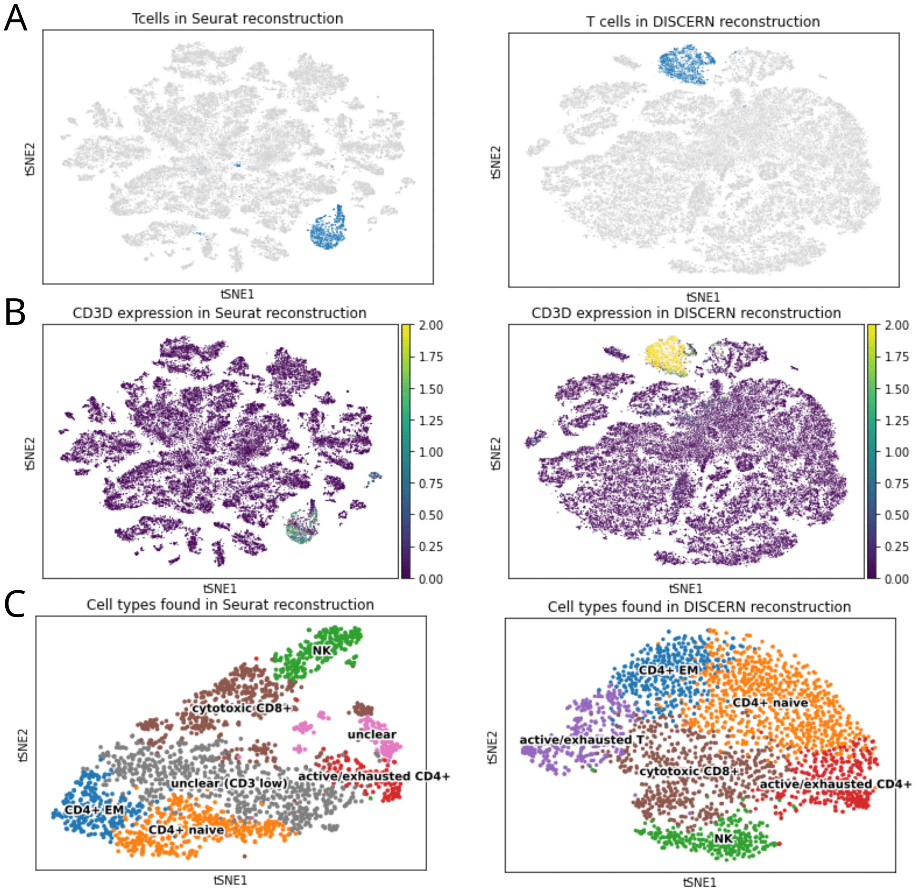


Figure S31: *T cell detection and sub-clustering in Kidney snRNA-seq (kidney-lq) and scRNA-seq (kidney-hq) data of patients with acute kidney injury.* **A**: tSNE representation of T cells found in Seurat (left) and DISCERN (right) reconstructed snRNA-seq and scRNA-seq data. **B**: tSNE representation of T cells found in Seurat (left) and DISCERN (right) reconstructed snRNA-seq and scRNA-seq data colored by *CD3D* expression as marker for T cells. **C**: tSNE representation of T cell subtypes found in Seurat (left) and DISCERN (right) reconstructed kidney-lq and kidney-hq data. A high number of cells in Seurat reconstruction could not be further classified due to absent or low expression of marker genes.

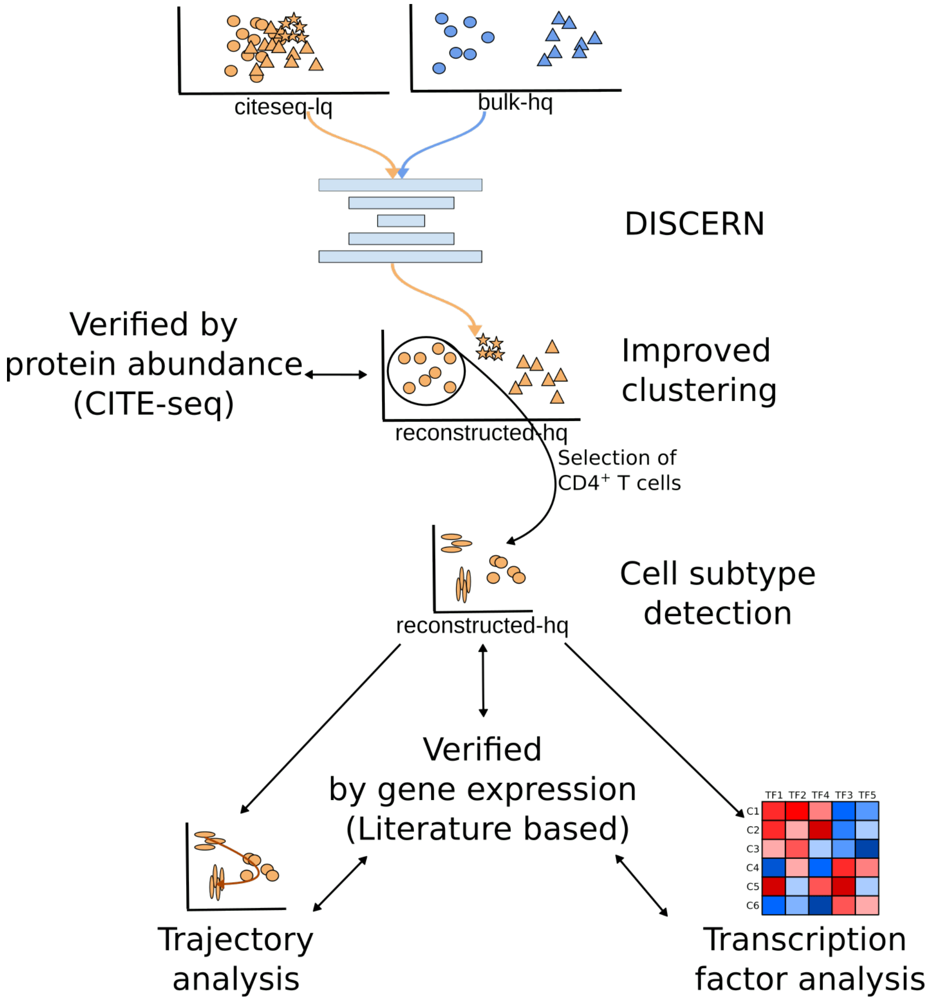


Figure S32: *Schematic representation of the experiments conducted with blood-based citeseq dataset.* To improve cite-lq data we reconstructed it with bulk-hq data to obtain reconstructed-hq data, which enabled improved clustering and $CD4^+$ T cell subtype detection. Additionally trajectory analysis and transcription factor analysis was performed on the $CD4^+$ T cell subset. Results were verified using protein abundance (CITE-seq) and literature information.

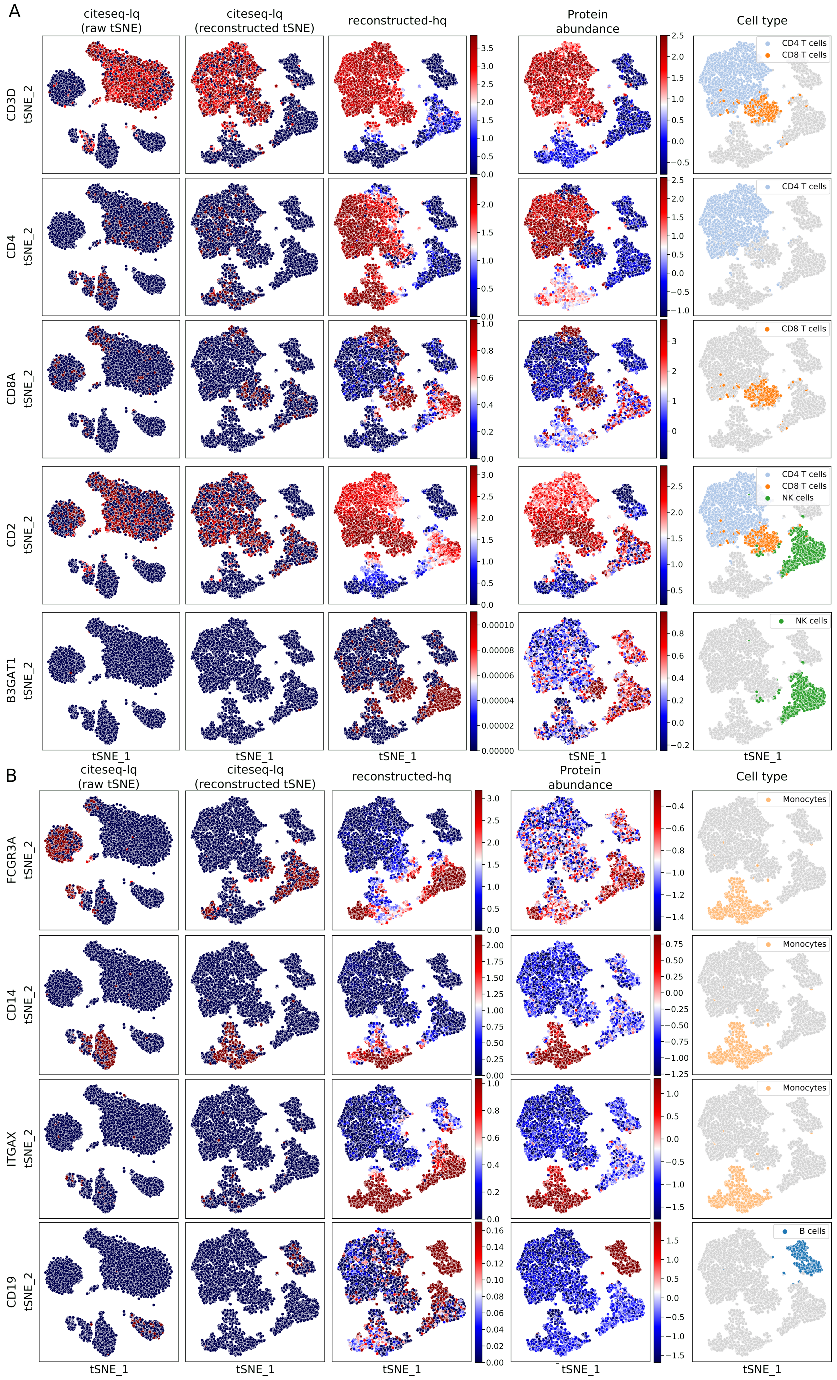
A
citeseq-lq (raw tSNE)
citeseq-lq (reconstructed tSNE)
reconstructed-hq
Protein abundance
Cell type
CD3D
CD4
CD8A
CD2
B3GAT1
tSNE_2
tSNE_1
CD4 T cells
CD8 T cells
NK cells
B
FCGR3A
CD14
ITGAX
CD19
Monocytes
B cells

Figure S33: *t-SNE representation of the gene expression and corresponding protein expression for the citeseq dataset before and after reconstruction.* **A** & **B**: Gene expression levels before reconstruction of the cite-lq data (first and second column) and after reconstruction to reconstructed-hq data using a bulk-hq reference (third column). Protein abundance measured using CITE-seq information is shown in the fourth column and the corresponding cell type in the fifth column. The first column shows tSNE representation computed on the uncorrected cite-lq data, while the others are computed on the reconstructed-hq data. Gene and protein expression levels are displayed in blue for low to red for high expression.

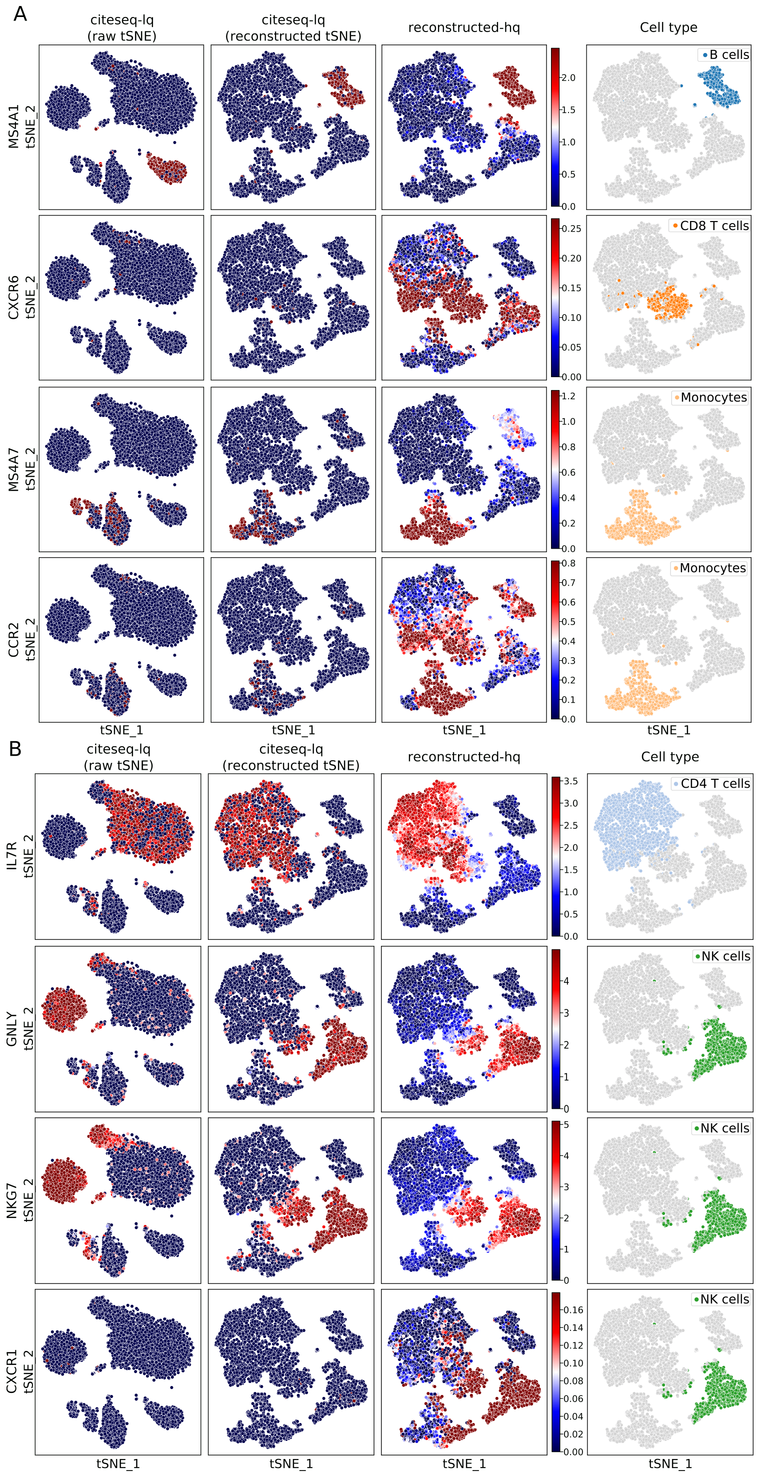
A
citeseq-lq (raw tSNE)
citeseq-lq (reconstructed tSNE)
reconstructed-hq
Cell type
MS4A1
CXCR6
MS4A7
CCR2
tSNE_2
tSNE_1
B cells
CD8 T cells
Monocytes
B
IL7R
GNLY
NKG7
CXCR1
CD4 T cells
NK cells

Figure S34: *t-SNE representation of the gene expression of genes without CITE-seq information for the citeseq dataset before and after reconstruction.* **A** & **B**: Gene expression levels before reconstruction of the cite-lq data (first and second column) and after reconstruction to reconstructed-hq data using a bulk-hq reference (third column). The corresponding cell type is displayed in the fourth column. The first column shows tSNE representation computed on the uncorrected cite-lq data, while the others are computed on the reconstructed-hq data. Gene expression levels are displayed in blue for low to red for high expression.

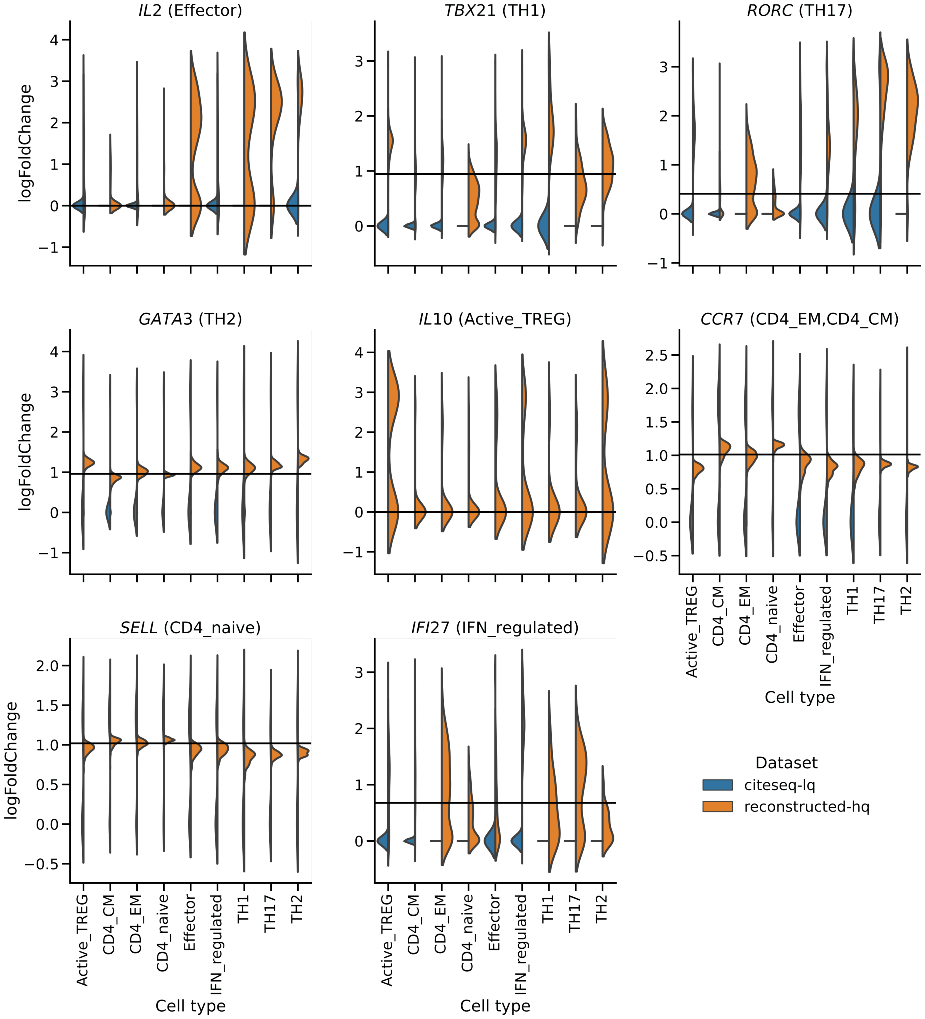


Figure S35: *Violin plots of cell type determining genes of CD4$^+$ T helper cell subtypes in the citeseq dataset. The expression is normalized by the mean over all cell types and log2-scaled.* Colors indicate whether they are shown for the uncorrected cite-lq data (blue) and the reconstructed-hq data (orange). The horizontal bars indicate median expression in the total citeseq dataset. The reconstructed gene expression is in many cases consistent with literature information. TH17 cells, for instance, are characterized by a high expression of *RORC* [5], TH2 cells express the transcriptional regulator *GATA3* [6], and TH1 cells the transcriptional regulator *TBX21* (*T-bet*) (fig. S36) [7]. *IL10* is produced by Foxp3 positive Treg cells (Active_TREG) [8]. *SELL* and *CCR7* are expressed in the CD4$^+$ T cell subtypes CD4_naive, CD4_EM, CD4_CM, but with a significantly lower expression of *CCR7* in CD4_EM cells [9], while CD4_naive cells show the highest expression of *SELL/CD62L* (fig. S37) [10].

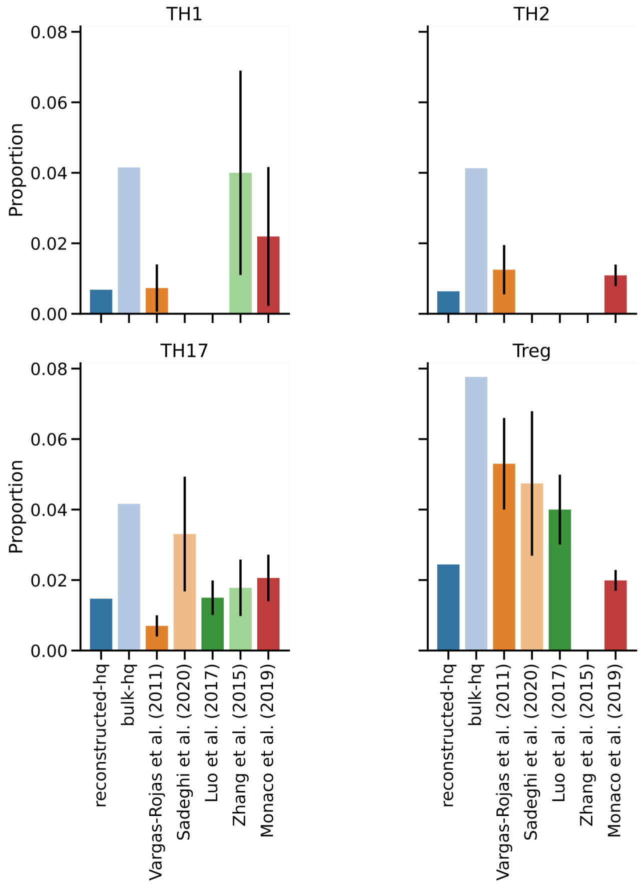


Figure S36: *Bar plot showing the proportions of CD4$^+$ T helper cell subtypes (TH1, TH2, TH17, and Treg) identified in the reconstructed-hq data, bulk-hq training data, and published ground-truth cell fractions in the citeseq data.* The proportions are calculated with respect to the total number of PBMCs. To compare the proportions in the reconstructed data with existing literature, five studies were considered (see also table S3). These studies estimate one or more of these subtypes using FACS and subsequent cell activation. For these references, bars represent means while error bars represent standard deviation. Missing bars indicate that the corresponding cell-type is not quantified in the referenced study.

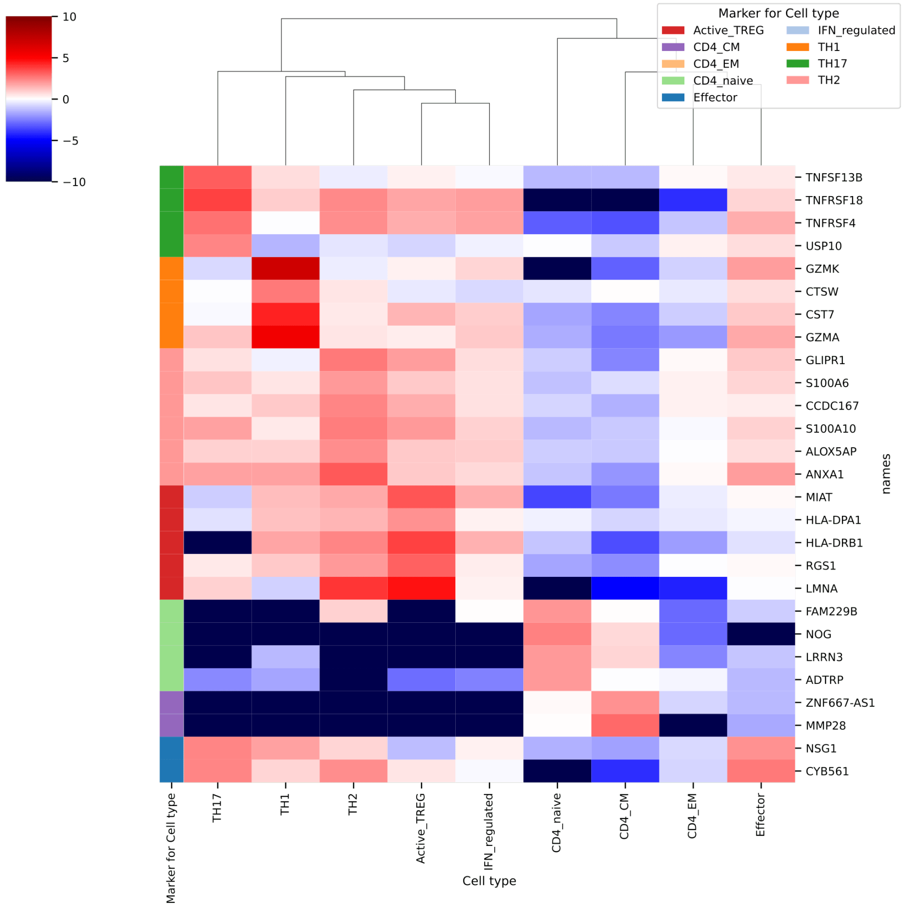


Figure S37: *Heatmap of log2 fold-change (FC) for the top CD4$^+$ T helper cell subtype marker genes in the citeseq dataset after reconstruction with DISCERN.* The cite-lq data was reconstructed with bulk-hq data to obtain reconstructed-hq data, which is displayed in the figure. Genes were filtered for an adjusted p-value $\leq 0.05$ and a log2 FC $\geq 2$. The top five genes with the lowest adjusted p-value were selected from the genes passing the threshold and duplicate gene entries were removed. For display reasons, the negative log2 FC was clipped at $-10$. The first column indicates cell type-specific expression according to the DEG analysis. FC magnitude is depicted with blue - low to red - high changes.

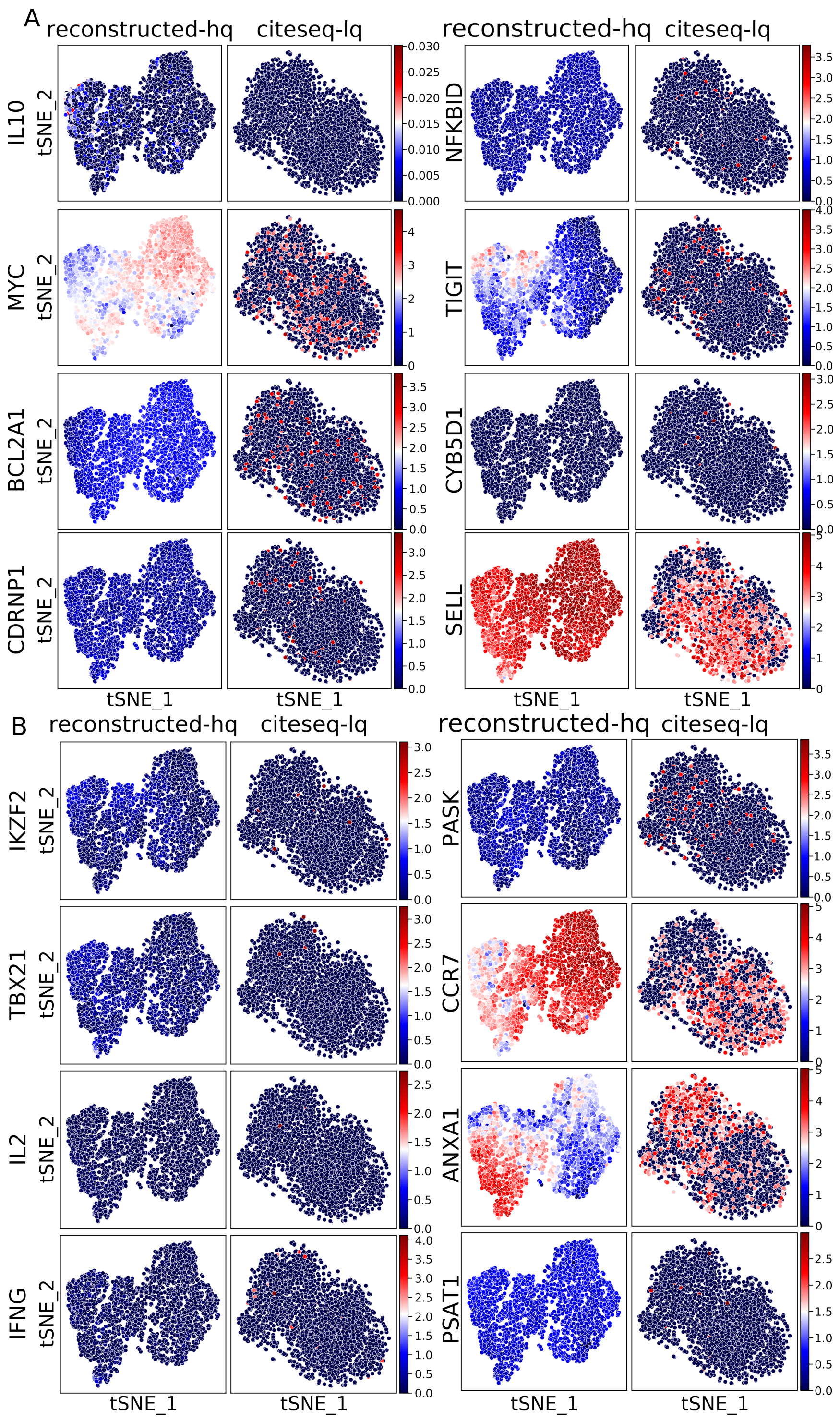

A
reconstructed-hq
citeseq-lq
IL10
MYC
BCL2A1
CDRNP1
NFKBID
TIGIT
CYB5D1
SELL
tSNE_2
tSNE_1
B
IKZF2
TBX21
IL2
IFNG
PASK
CCR7
ANXA1
PSAT1

Figure S38: *t-SNE representation of the gene expression of several established T cell marker genes in citeseq $CD4^+$ T cells before and after correction with DISCERN.* **A** & **B**: Gene expression levels before reconstruction of the cite-lq data (third and fourth column, t-SNE calculated on cite-lq data) and after reconstruction to reconstructed-hq data using a bulk-hq reference (first and third column, t-SNE calculated on reconstructed-lq data). Gene expression levels are displayed in blue for low to red for high expression. *MYC*, *NFKBID*, *BCL2A1*, *CYB5D1*, *CSRNP1*, *IL2*, and *PSAT1* are used as activation markers, *IL10*, *TBX21*, *ANXA1*, *IFNG* characterize TH1 cells, *TIGIT* and *PASK* TFH cells, *IKZF2* TREG, *CCR7* central memory T cells and *SELL* is a marker for naive T cells. In general, the cell type-specific expression of published marker genes in the reconstructed-hq data show good correspondence with the identified cell types.

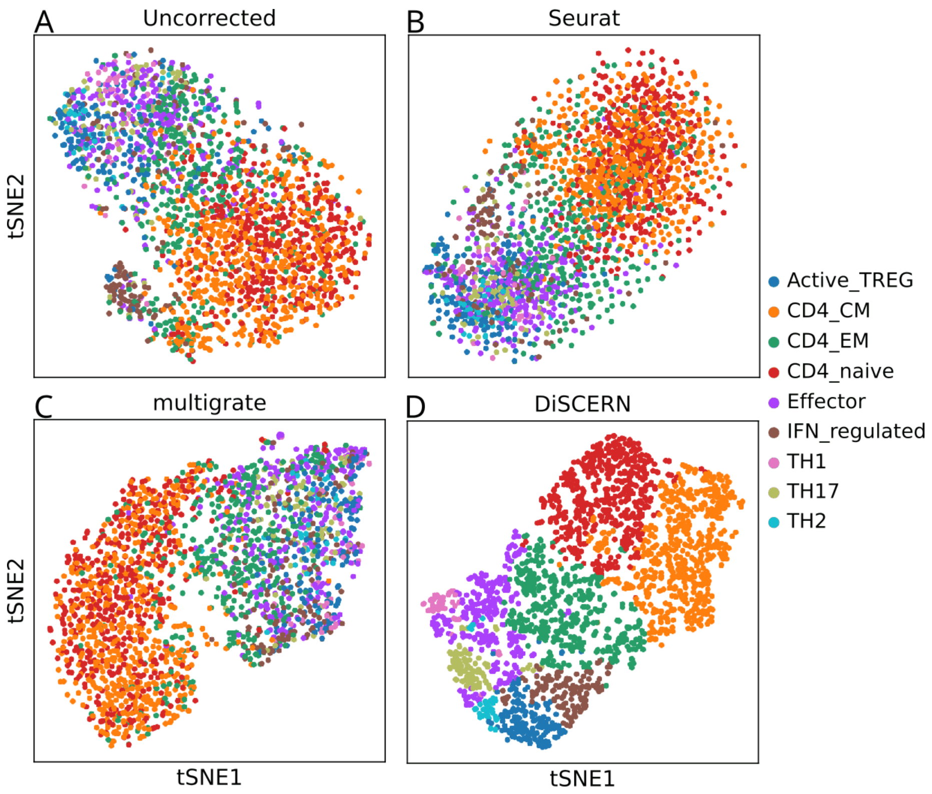


Figure S39: *tSNE representation of $CD4^+$ T cells in the citeseq dataset after annotation of the cell types found after expression reconstruction with DISCERN.* **A**: Uncorrected citeseq-lq $CD4^+$ T cells show some clustering of IFN_regulated, Active_TREG, CD4_CM and CD4_naive cells. **B**: Seurat reconstruction results in strong cluster and cell type mixing, a potential sign of overintegration. **C**: Multigrate imputed data shows strong mixing and splitting of clusters and cell types, for instance $CD4^+$ T cells are split into two clusters. **D**: DISCERN reconstructed cite-hq data provides a clear separation of functionally distinct cell types.

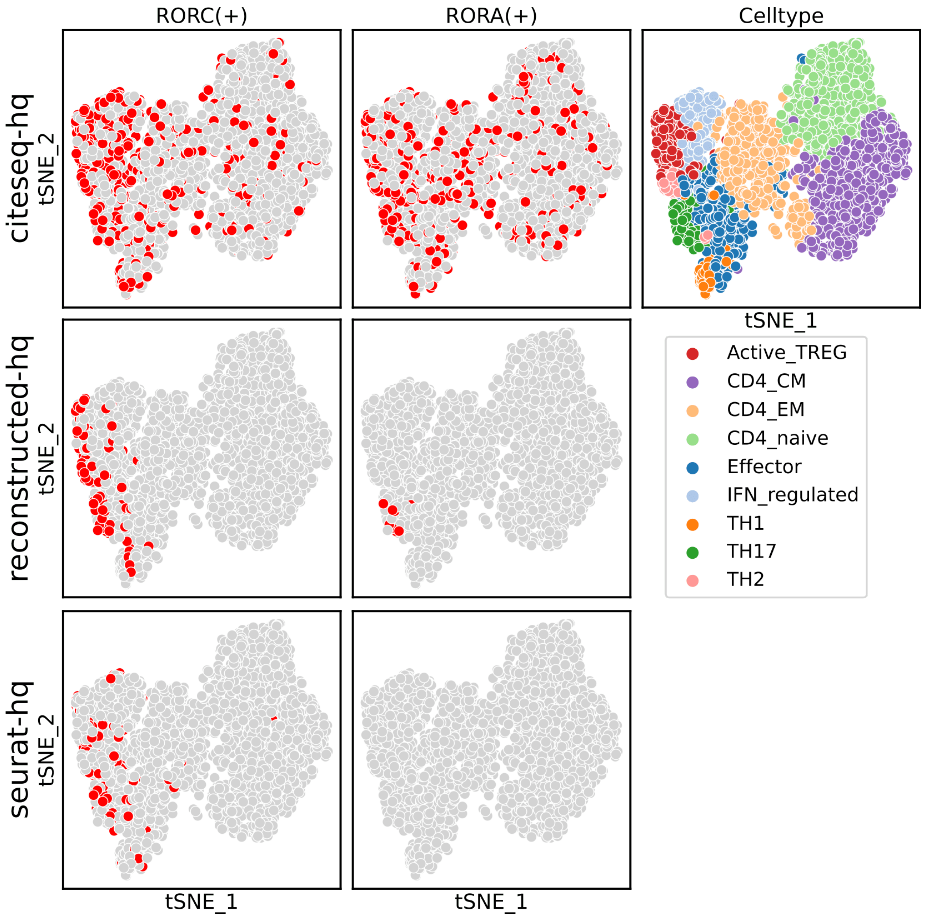


Figure S40: *t-SNE representation of a SCENIC transcriptional regulation analysis of citeseq T helper cells before and after correction with DISCERN and Seurat.* Gene expression levels before reconstruction of the cite-lq data (first row) and after reconstruction to reconstructed-hq data using a bulk-hq reference with DISCERN (second row) and Seurat (third row). The first column displays cells that express the RORC(+) regulon. The second column displays cells that express the RORA(+) regulon. Red color in the first two columns depends on the binarized AUCell score of the SCENIC discovered regulons. The third column displays the detected cell types in reconstructed-hq data. RORA(+) and RORC(+) regulons are expected to be specific for TH17 cells. The tSNE representation is calculated on DISCERN reconstructed data.

A

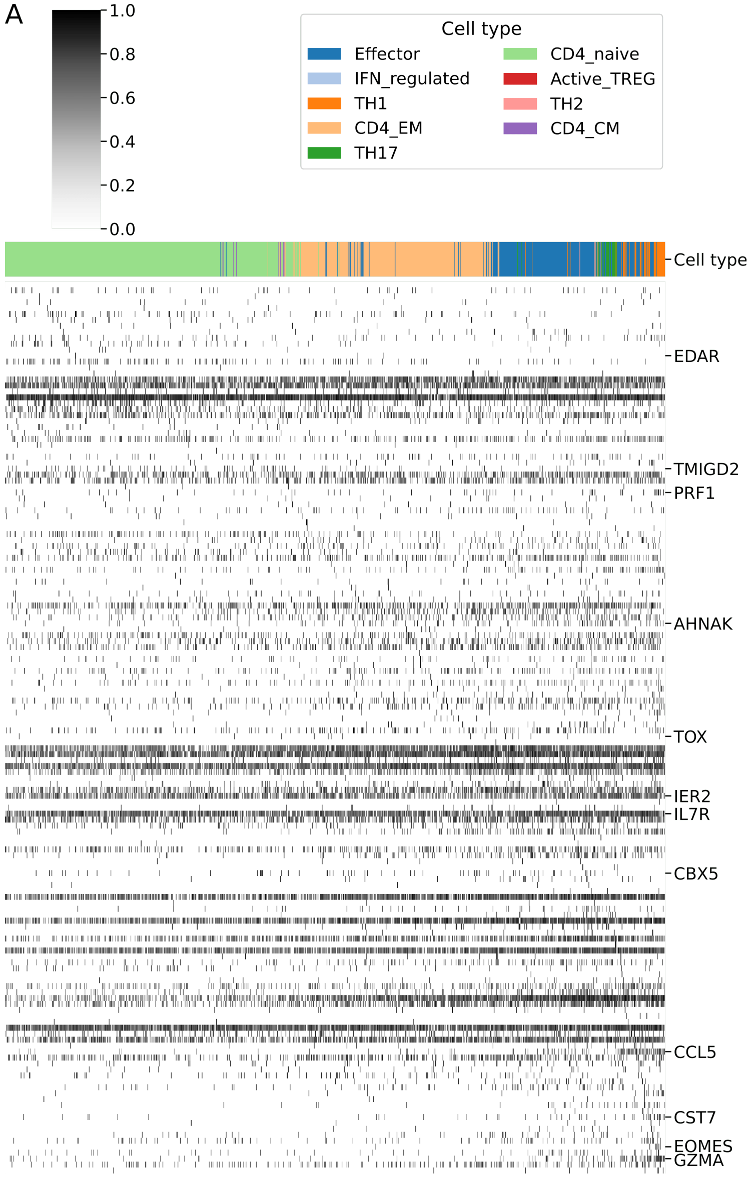

1.0
0.8
0.6
0.4
0.2
0.0
Cell type
Effector
CD4_naive
IFN_regulated
Active_TREG
TH1
TH2
CD4_EM
CD4_CM
TH17
Cell type
EDAR
TMIGD2
PRF1
AHNAK
TOX
IER2
IL7R
CBX5
CCL5
CST7
EOMES
GZMA

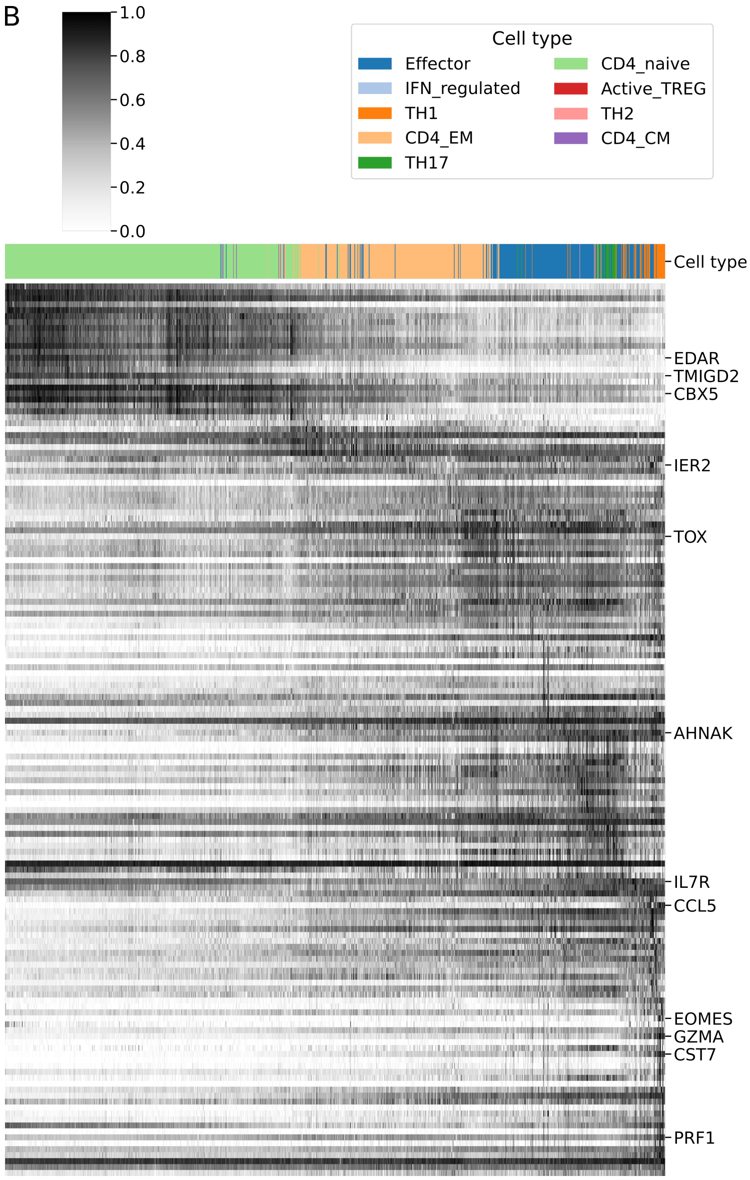
B
1.0
0.8
0.6
0.4
0.2
0.0
Cell type
Effector
IFN_regulated
TH1
CD4_EM
TH17
CD4_naive
Active_TREG
TH2
CD4_CM
Cell type
EDAR
TMIGD2
CBX5
IER2
TOX
AHNAK
IL7R
CCL5
EOMES
GZMA
CST7
PRF1

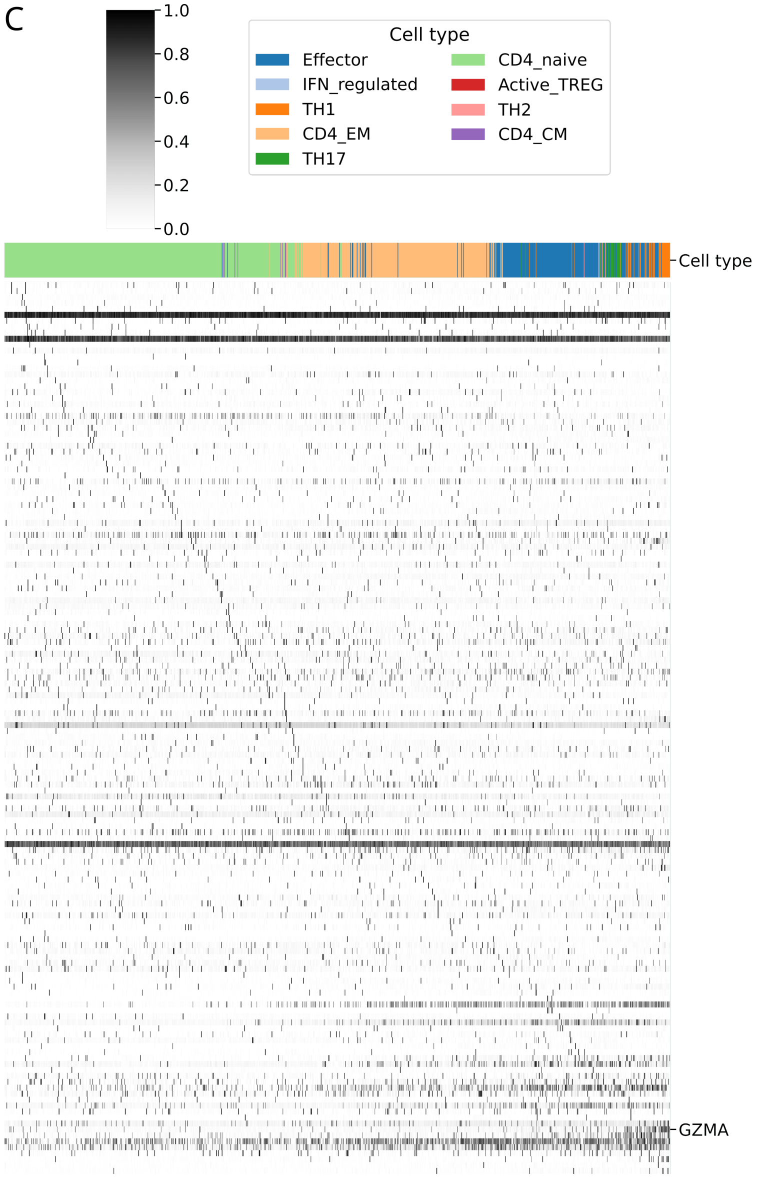


Figure S41: *Expression of differentially regulated genes of the CD4_naive to TH1 lineage (Lineage1) defined by a Slingshot trajectory analysis.* The top 150 genes by p-value are shown. Only a selection of T cell marker genes is shown by name. The cell types are color-coded and cells are sorted by pseudotime. **A**: Expression using the cite-lq data before reconstruction. **B**: Expression using the reconstructed-hq data that was reconstructed with DISCERN using a bulk-hq reference. In the reconstructed-hq data, Lineage1 shows a trajectory from *TMIGD2*,

*EDAR* and *CBX5* expressing CD4_naive cells [11, 12] to TH1 cells expressing cytotoxicity-related genes like *EOMES*, *CST7*, *GZMA*, *IL7R*, *CCL5* and *PRF1* [13, 14, 15, 16, 17, 18, 19]. The cells develop through CD4_EM cells to a (pre-) effector state (Effector cells) to the final TH1 subtype (fig. S42B). Effector and CD4_EM cells show higher expression of *IER2* [20], *AHNAK* [21] and *TOX* [22], as reported in the literature. In general, reconstructed data show cell trajectories that are biologically reasonable, while uncorrected data shows little structure. **C**: Expression using the Seurat to reconstruct the data (seurat-hq). While the heatmap shows a small trajectory line, most of the marker genes are not found along the trajectory.

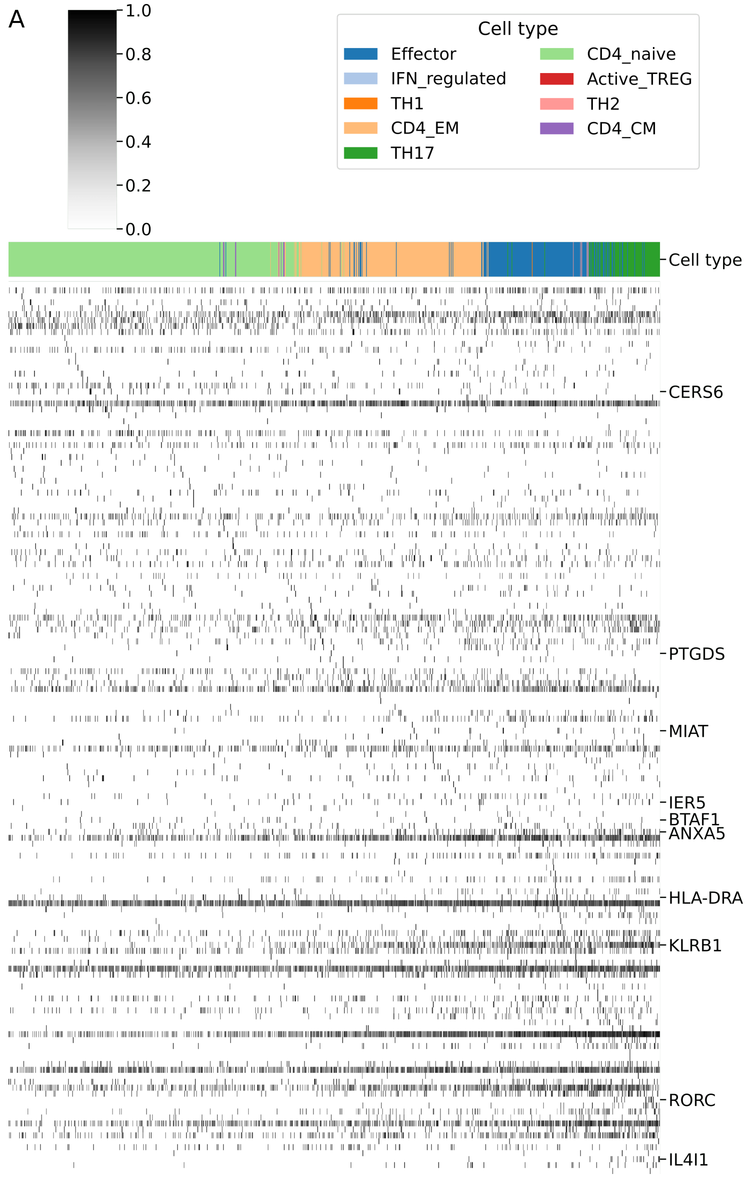

A
1.0
0.8
0.6
0.4
0.2
0.0
Cell type
Effector
IFN_regulated
TH1
CD4_EM
TH17
CD4_naive
Active_TREG
TH2
CD4_CM
Cell type
CERS6
PTGDS
MIAT
IER5
BTAF1
ANXA5
HLA-DRA
KLRB1
RORC
IL4I1

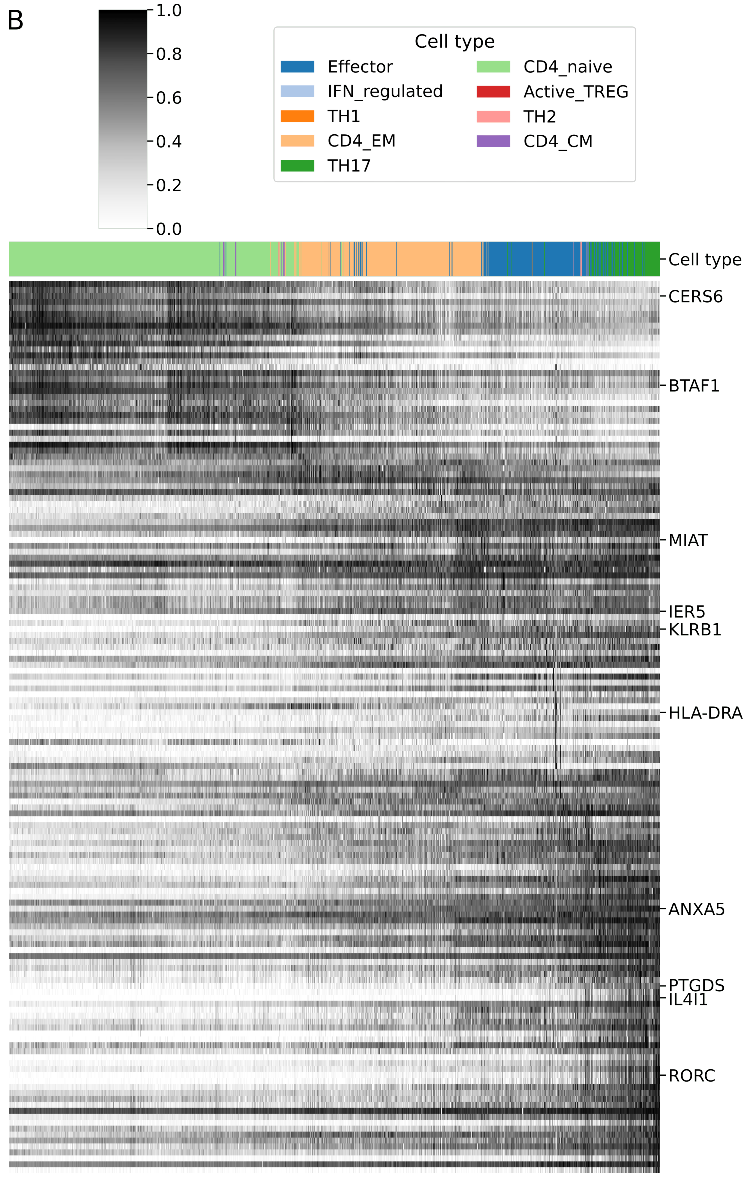
B
1.0
0.8
0.6
0.4
0.2
0.0
Cell type
Effector
IFN_regulated
TH1
CD4_EM
TH17
CD4_naive
Active_TREG
TH2
CD4_CM
Cell type
CERS6
BTAF1
MIAT
IER5
KLRB1
HLA-DRA
ANXA5
PTGDS
IL4I1
RORC

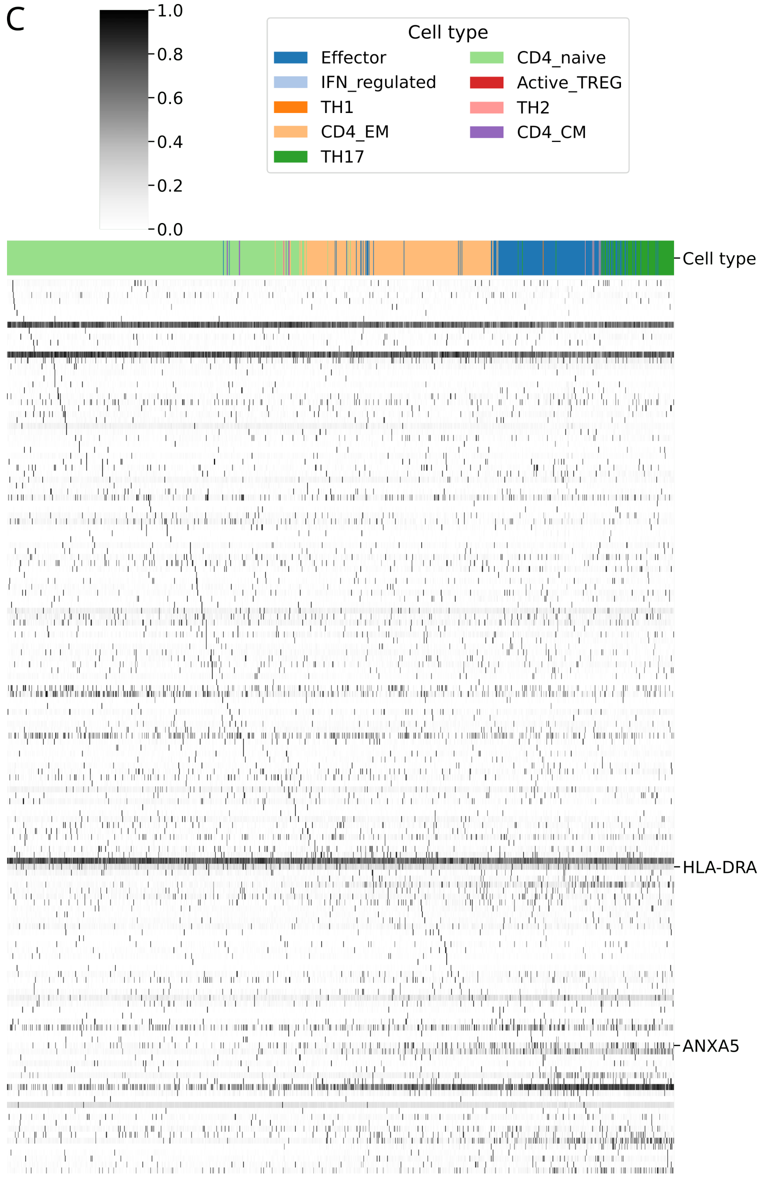


Figure S42: *Expression of differentially regulated genes of the CD4_naive to TH17 lineage (Lineage2) defined by a Slingshot trajectory analysis.* The top 150 genes by p-value are shown. Only a selection of T cell marker genes is shown by name. The cell types are color-coded and cells are sorted by pseudotime. **A:** Expression using the cite-lq data before reconstruction. **B:** Expression using the reconstructed-hq data that was reconstructed with DISCERN using a bulk-hq reference. In the reconstructed-hq data, the CD4_naive cells show the expected

*BTAF1* and *CERS6* expression [12, 23], whereas effector cells express activation markers as MIAT [24], *HLA-DRA* [25], *IER5* [26] and *KLRB1* [27]. Finally the trajectory terminates in TH17 cells, expressing *ANXA5* [12], *RORC* [5], *IL4I1* [28] and *PTGDS* [29], showing that the found trajectory (Fig. 3C) is in line with known expression patterns. **C**: Expression using the Seurat to reconstruct the data (seurat-hq). While the heatmap shows a small trajectory line, most of the marker genes are not found along the trajectory.

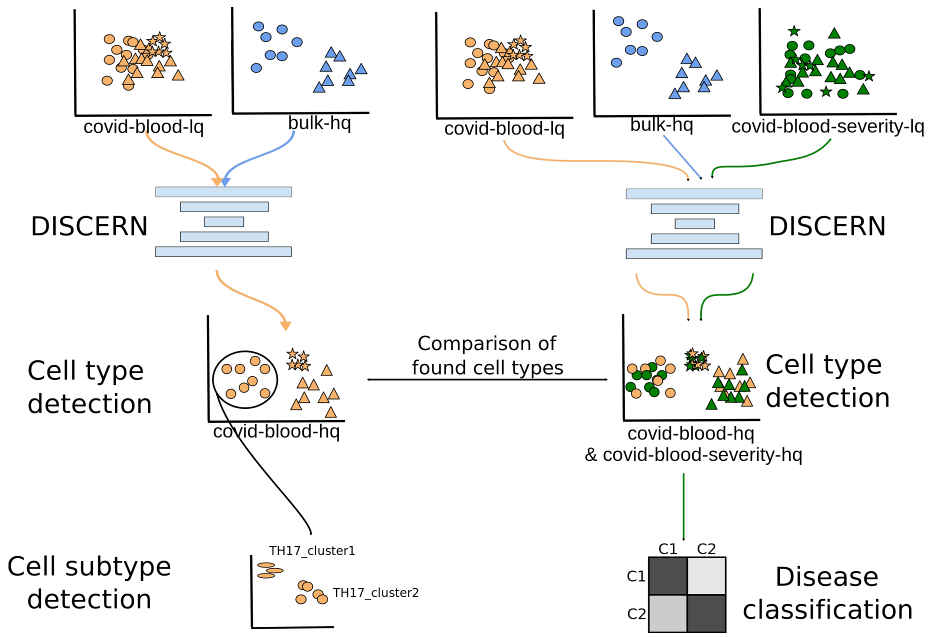


Figure S43: *Schematic representation of the experiments conducted with the covid-blood and covid-blood-severity datasets.* To improve covid-blood-lq and covid-blood-severity-lq data we reconstructed it using bulk-hq data to obtain covid-blood-hq and covid-blood-severity-hq data, respectively. We then performed cell type detection using the reconstructed data. We investigated T helper cell subtypes in great detail in the covid-blood-hq data and compared them to the ones found in the covid-blood-severity-hq data. Finally, we used the covid-blood-severity dataset and its disease severity information for COVID-19 patients to classify mild and severe cases using a GBM. TH17 cell subtypes could be detected in the covid-blood-hq data and linked to cells found in the covid–lung dataset using T cell receptor clonal information.

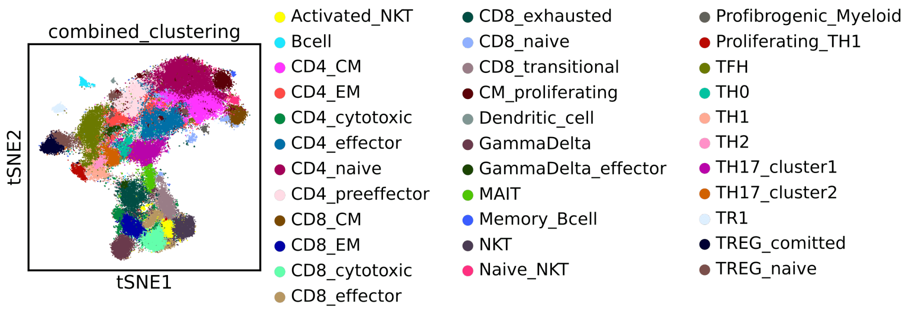


Figure S44: *t-SNE representation of cell types found in the covid-blood-hq data.* The covid-blood-lq data was reconstructed using bulk-hq data to obtain covid-blood-hq data. Colors indicate the annotated cell types. Especially T cell subtypes could not be annotated before reconstruction with DISCERN. It is especially interesting that TH17 subtypes can be detected, which are usually observed in FACS PBMC data only after stimulation.

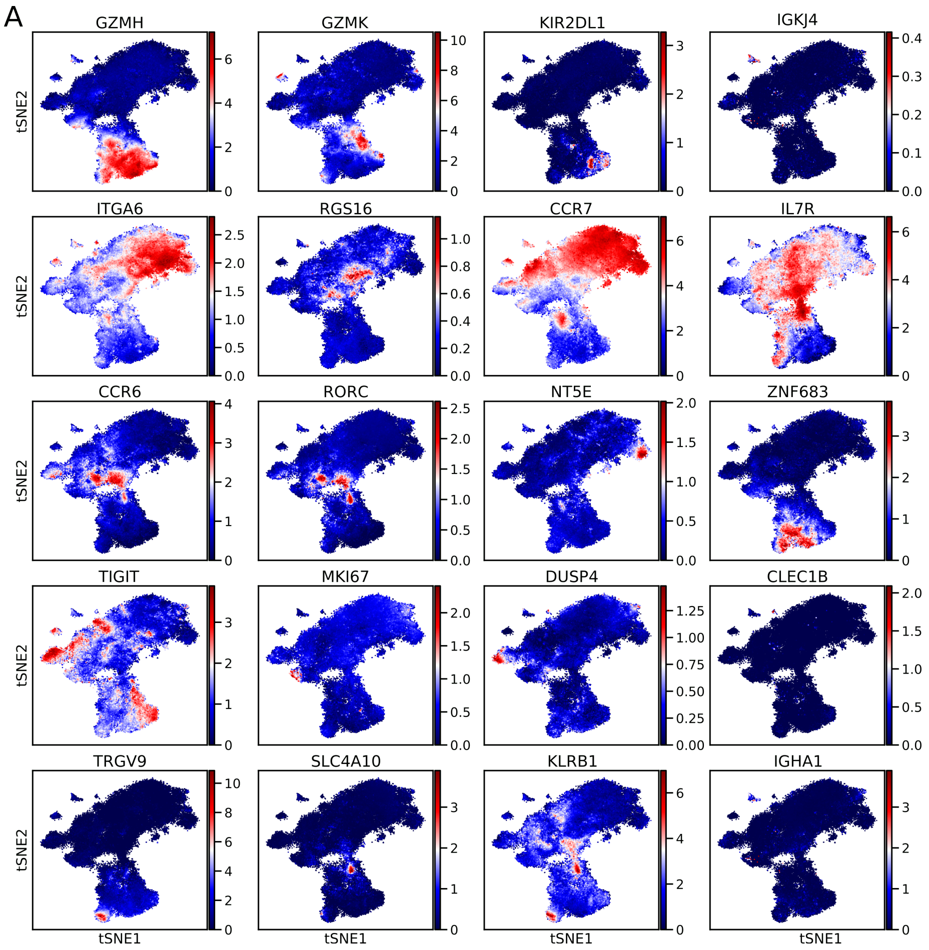
A
GZMH
GZMK
KIR2DL1
IGKJ4
ITGA6
RGS16
CCR7
IL7R
CCR6
RORC
NT5E
ZNF683
TIGIT
MKI67
DUSP4
CLEC1B
TRGV9
SLC4A10
KLRB1
IGHA1
tSNE2
tSNE1

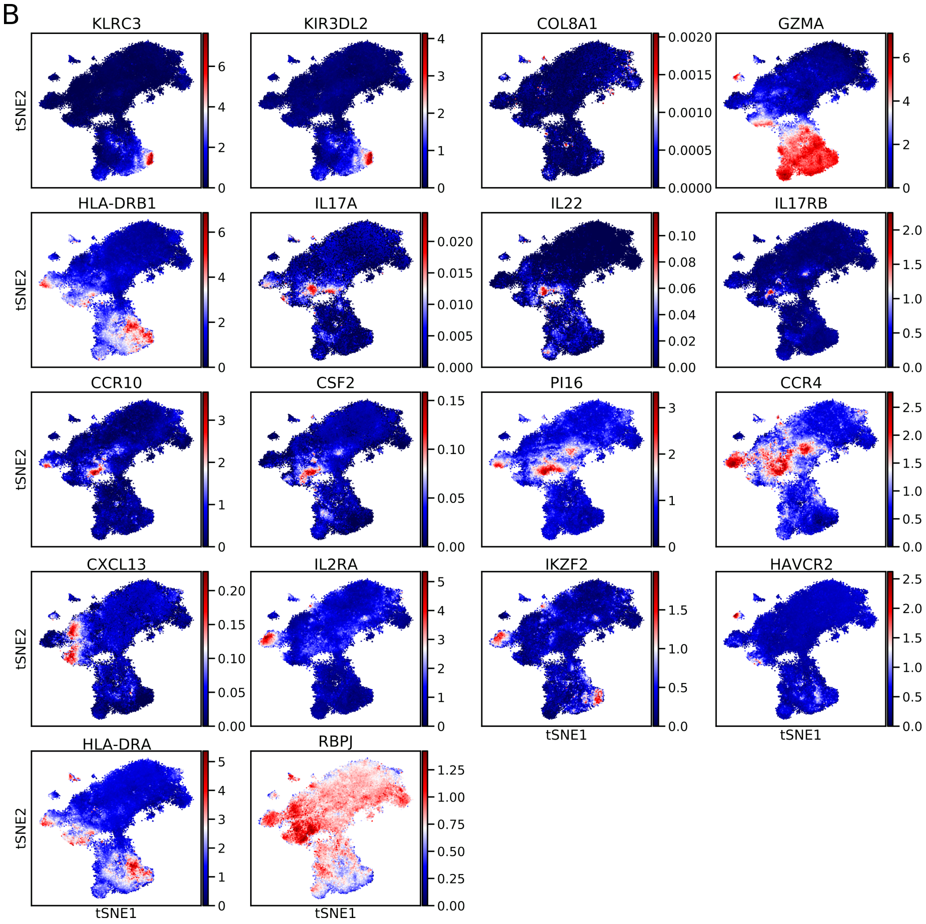


Figure S45: *t-SNE representation of the gene expression of several established marker genes in covid-blood-hq data.* **A** & **B**: The covid-blood-lq dataset after reconstruction with DISCERN using bulk-hq data to obtain covid-blood-hq data. The t-SNE representation was computed on the covid-blood-hq data. Gene expression levels are displayed in blue for low to red for high expression. In general, the cell type-specific expression of published marker genes in the covid-blood-hq data show good correspondence with the identified cell types.

A

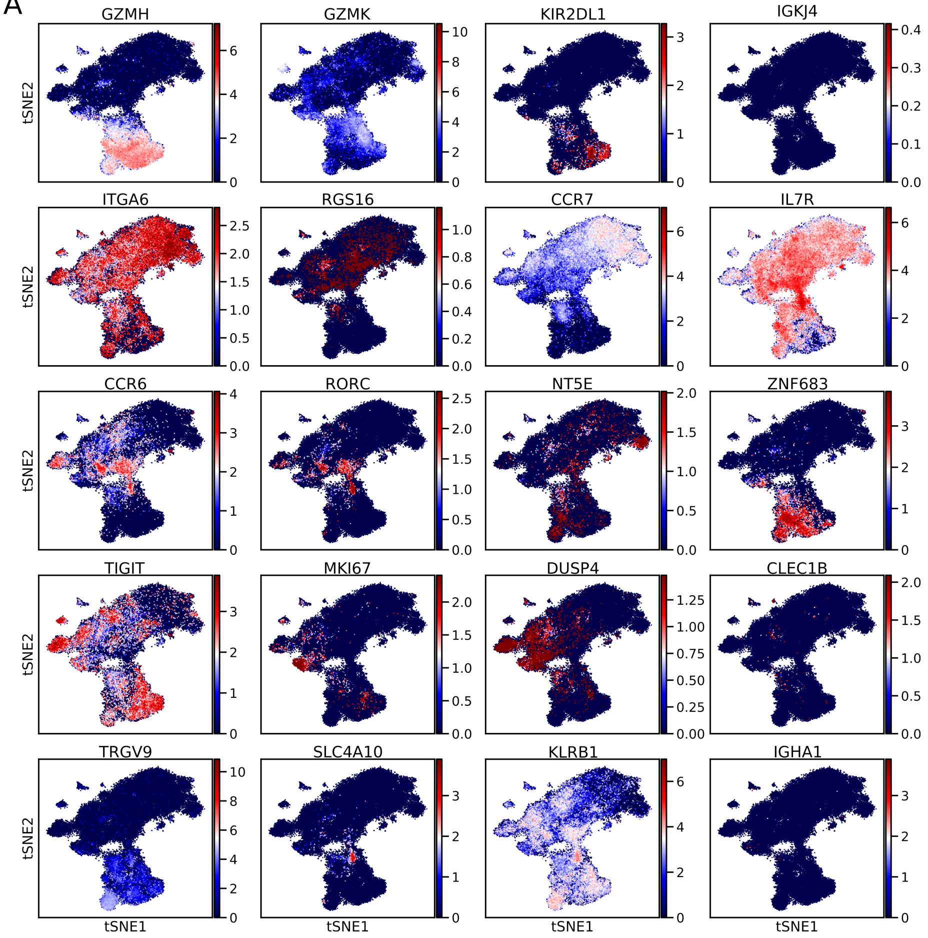
GZMH
GZMK
KIR2DL1
IGKJ4
ITGA6
RGS16
CCR7
IL7R
CCR6
RORC
NT5E
ZNF683
TIGIT
MKI67
DUSP4
CLEC1B
TRGV9
SLC4A10
KLRB1
IGHA1
tSNE2
tSNE1

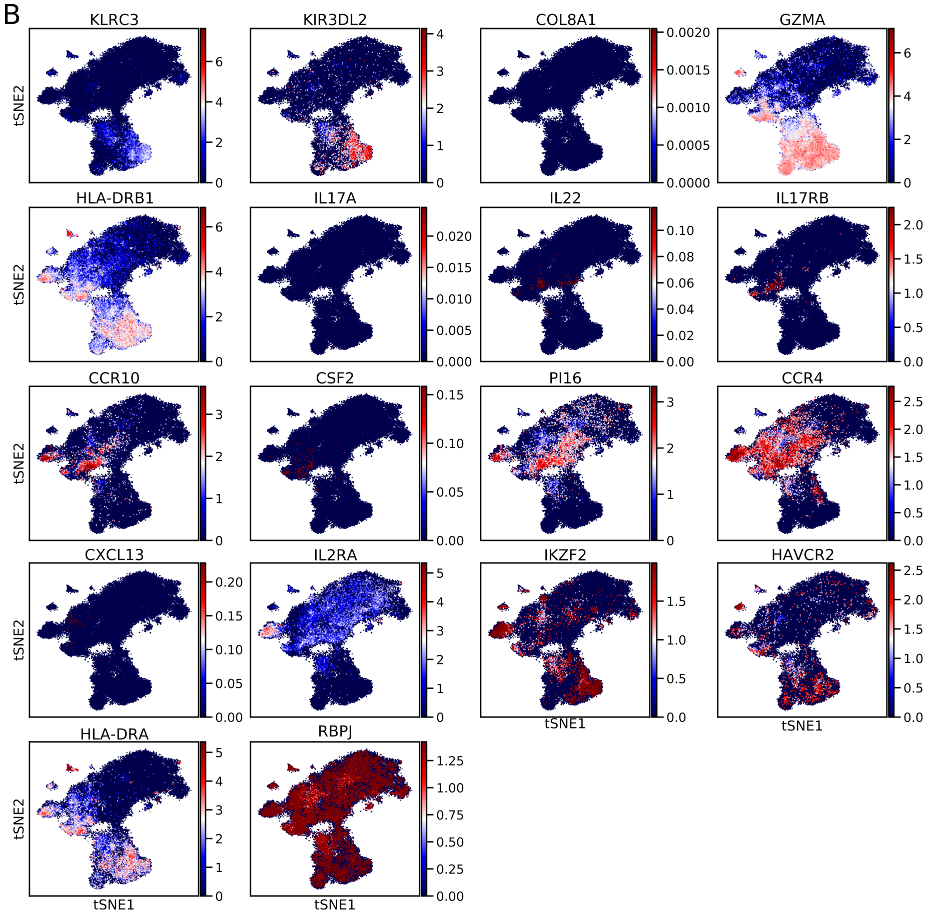


Figure S46: t-SNE representation of the gene expression of several established marker genes in covid-blood-lq data. **A** & **B**: The t-SNE representation was computed on the covid-blood-lq data without reconstruction. Gene expression levels are displayed in blue for low to red for high expression. In general, the cell type-specific expression of published marker genes in the covid-blood-lq data shows worse correspondence with the identified cell types as compared to the covid-blood-hq data in fig. S45.

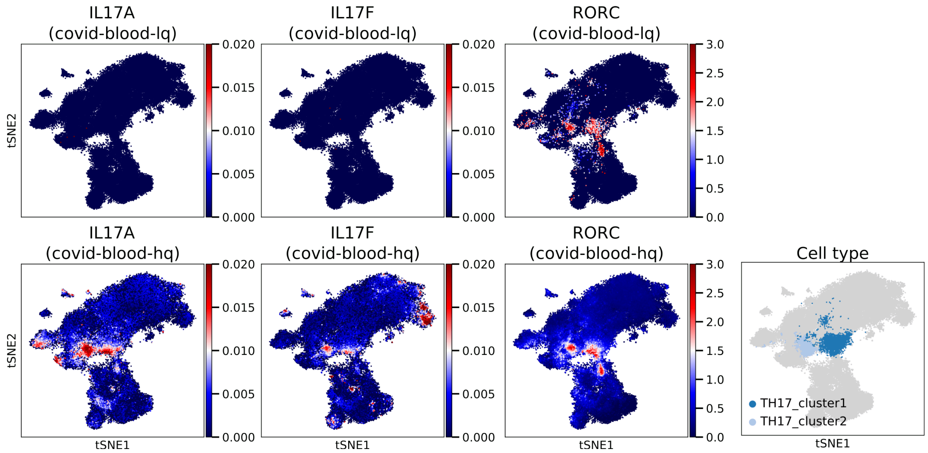


Figure S47: *t-SNE representation of TH17 marker genes in two TH17 subtypes detected in COVID-19 patient blood.* t-SNEs were calculated for $CD4^+$ T cells on covid-blood-hq data. The first row shows the expression of marker genes for uncorrected covid-blood-lq data. The second row displays the expression of the same marker genes for reconstructed covid-blood-hq data. The covid-blood-lq data was reconstructed using the bulk-hq reference to obtain covid-blood-hq data. The TH17 cell subclusters were found by louvain clustering after reconstruction. Colors represent the expression levels of genes as mentioned in the plot titles (*IL17A*, *IL17F*, *RORC*; from left to right). Expression levels of TH17 marker gene expression is barely visible for *IL17A/F* before reconstruction but can be detected after reconstruction with DISCERN. *RORC*, as transcription factor for TH17 cells, confirms the correct annotation of TH17 cells.

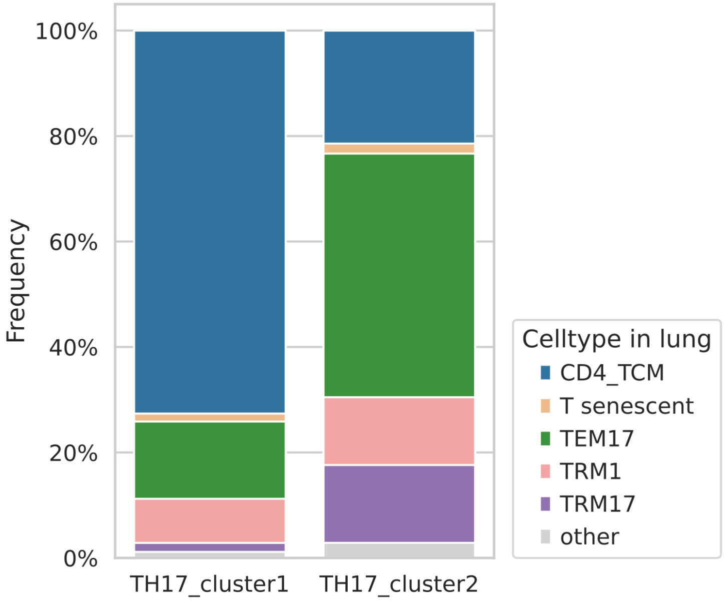


Figure S48: *Fraction of TH17 cells sharing the T cell receptor clonotype in covid-blood-hq and covid-lung data.* Cell type annotations of lung data were used as provided in the original publication. Cell types with an overlap $< 1$ % in both TH17 clusters were labeled as other. TH17_cluster1, detected in covid-blood-hq data, shares T cell receptor clones with CD4_TCM cells in the covid-lung data. TH17_cluster2, detected in covid-blood-hq data, shares most T cell receptor clones with TEM17 cells in covid-lung data. This corroborates the definition of the two TH17 subtypes detected in covid-blood-hq data and raises the question if these cells stay peripheral or re-enter tissues to promote inflammation.

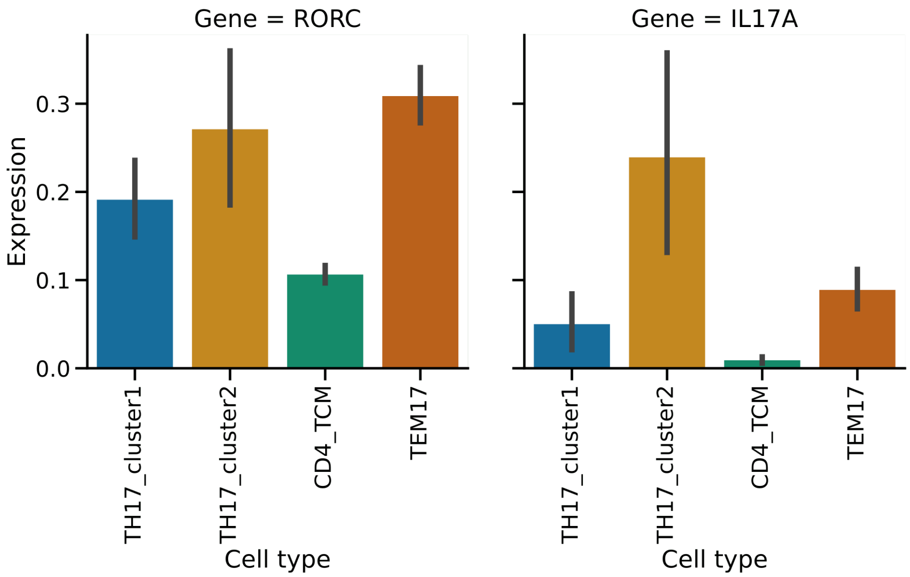


Figure S49: *Mean expression of RORC and IL17-A of covid-lung cells sharing a clonotype with TH17 cells of the covid-blood-hq data.* TH17_cluster1 and TH17_cluster2 are determined using the TCR clonotype information of reconstructed covid-blood-hq data and CD4_TCM or TEM17 covid-lung cells were annotated as in the original publication (see also fig. S48). A single cell can contribute to more than one bar, e.g. by being annotated as TEM and having a shared clonotype with TH17_cluster2 cell in covid-blood. Cell types sharing a clonotype with TH17_cluster1 and TH17_cluster2 cells from covid-blood have on average a higher or similar expression of the TH17 marker genes (*RORC* and *IL17A*) than cells in CD4_TCM or TEM17 cells in lung. This shows that CD4_TCM and TEM17 can most likely be further subdivided into clusters matching TH17_cluster1 and TH17_cluster2 in covid-blood and thus giving more evidence that these cell subtypes have a biological role in blood and lung.

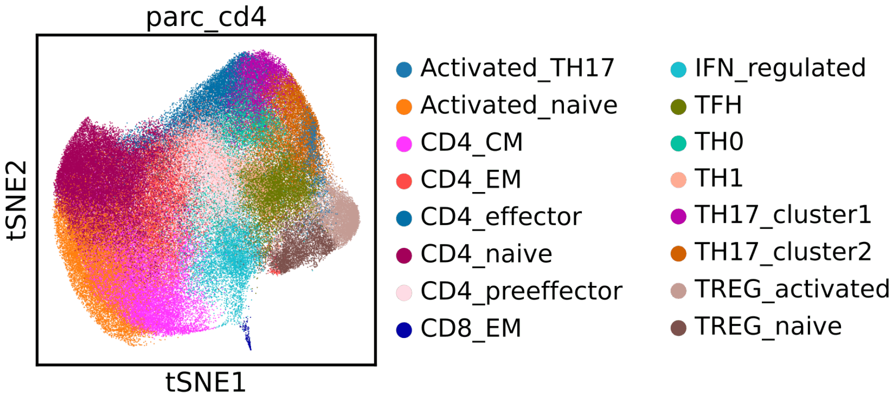


Figure S50: *t-SNE representation of $CD4^+$ T cells found in the covid-blood-severity-hq data.* The t-SNE representation and clustering was computed on DISCERN reconstructed expression, using covid-blood-severity-lq and covid-blood-lq data as input and bulk-hq as reference. Cell types are labeled by color. It is interesting to observe that the detected cell types largely overlap for the two studies.

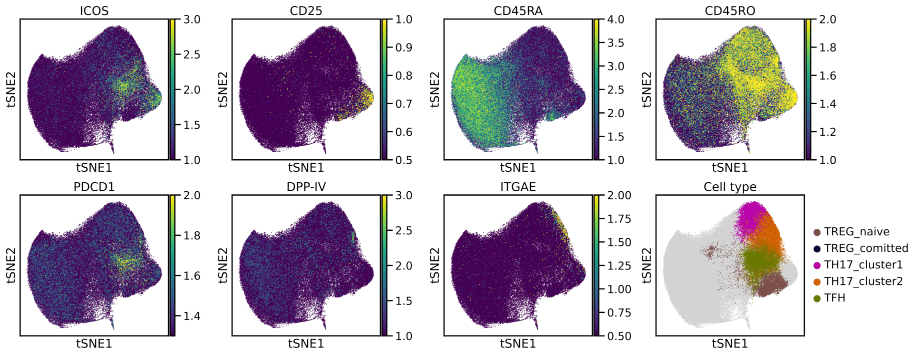


Figure S51: *t-SNE representation of the marker protein abundance in the covid-blood-severity dataset for TREG, TH17 and TFH cells provided by CITE-seq information.* The t-SNE representation and clustering was computed on DISCERN reconstructed expression, using covid-blood-severity-lq and covid-blood-lq data as input and bulk-hq as reference. The CITE-seq protein abundance of the covid-blood-severity data for seven marker proteins is displayed in color (blue - low to yellow - high abundance). The region we identified as regulatory T cells is positive for CD25 and $CD45RO^{+}$ and activated TREGs are high in ICOS as described for highly suppressive TREG [30]. TFH cells are PDCD1 and ICOS surface protein positive cells [31] and DPP-IV is markedly increased in activated TH17 cells expressing IL17A, a TH17-specific signal as previously described [32]. ITGAE abundance was described for resident T Helper cells in the skin reentering circulation [33]. In general, the CITE-seq information confirms the cell type identification of DISCERN reconstructed covid-blood-severity-hq data.

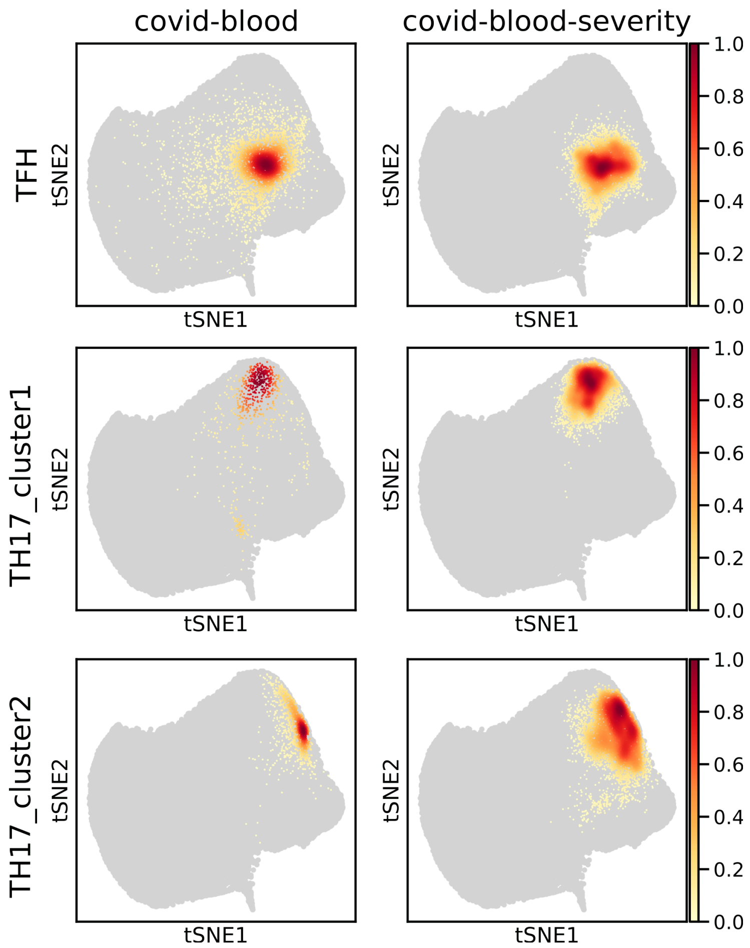


Figure S52: *t-SNE representation of three T helper cell clusters found in reconstructed covid-blood-hq and covid-blood-severity-hq data.* The t-SNE representation and clustering was computed on DISCERN reconstructed expression, using covid-blood-severity-lq and covid-blood-lq data as input and bulk-hq as reference. Cell type annotations for covid-blood-hq data are shown in the first column and covid-blood-severity-hq data in the second column. Cell densities are represented using white - low to red - high cell type density.

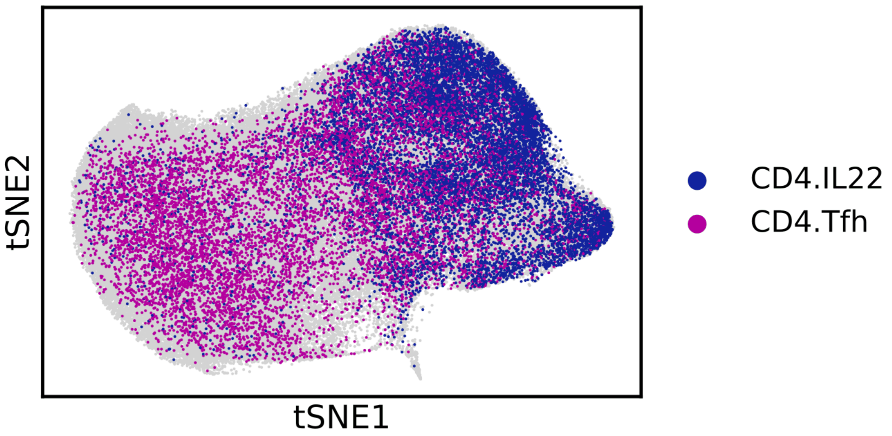


Figure S53: *tSNE representation of three T helper cell clusters of the reconstructed covid-blood and covid-blood-severity datasets.* The t-SNE representation and clustering was computed on DISCERN reconstructed expression, using covid-blood-severity-lq and covid-blood-lq data as input and bulk-hq as reference. Cell types are color-coded according to the covid-blood-severity-hq dataset. TFH cells from the original publication (CD4.Tfh) show significant overlap with naive $CD4^+$ T cells and $CD4^+$ $IL22^+$ cells (CD4.IL22) show marked overlap with TREG cells (compared with fig. S50).

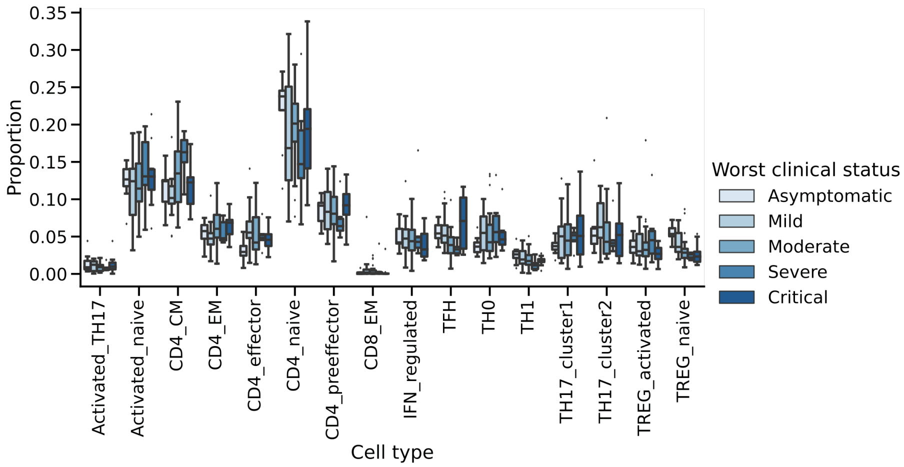


Figure S54: *Proportion of T cell subtypes in the covid-blood-severity-hq data grouped by disease severity.* The disease severity per patient is determined as the worst clinical status during hospitalization. Colors indicate disease severity, from light blue - asymptomatic to dark blue - critical. Boxplots represent median, quantiles, minimum, maximum, and potential outliers.

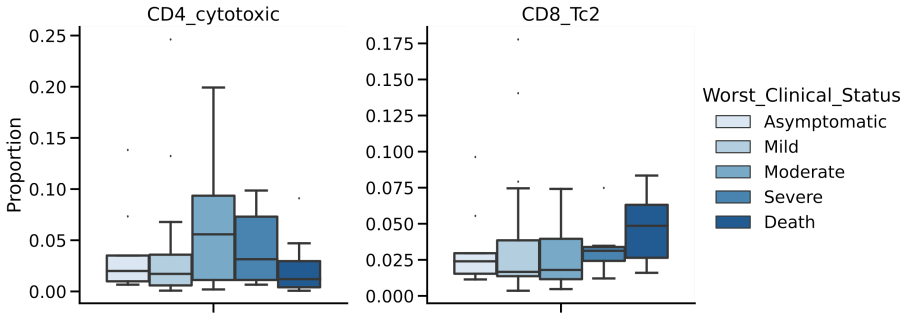


Figure S55: Proportion of 'unexpected' CD4_cytotoxic and CD8_Tc2 cell subtypes in the covid-blood-severity-hq data grouped by disease severity. The disease severity per patient is determined as the worst clinical status during hospitalization. Colors indicate disease severity, from light blue - asymptomatic to dark blue - critical. Boxplots represent median, quantiles, minimum, maximum, and potential outliers.

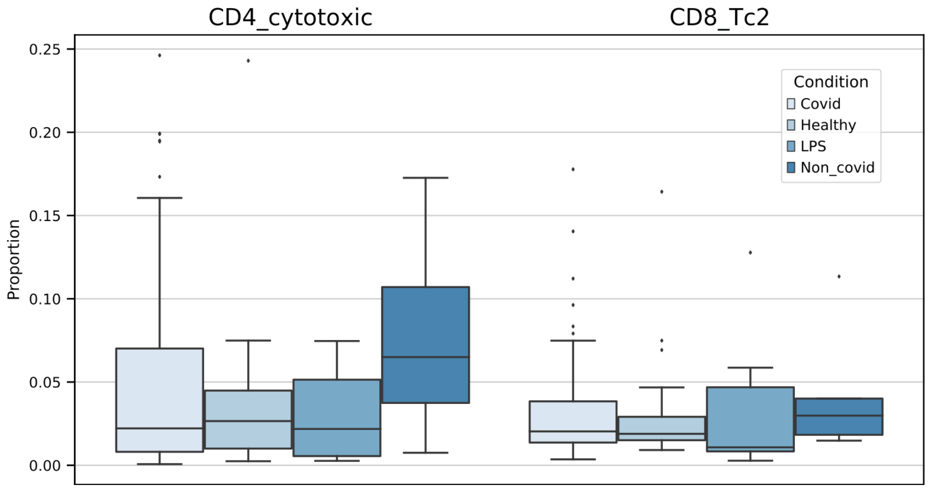


Figure S56: Proportion of 'unexpected' CD4_cytotoxic and CD8_Tc2 cell subtypes in the covid-blood-severity-hq data grouped by disease etiology. Colors indicate disease etiology, from light blue - COVID-19 to dark blue - non COVID-19. Boxplots represent median, quantiles, minimum, maximum, and potential outliers.

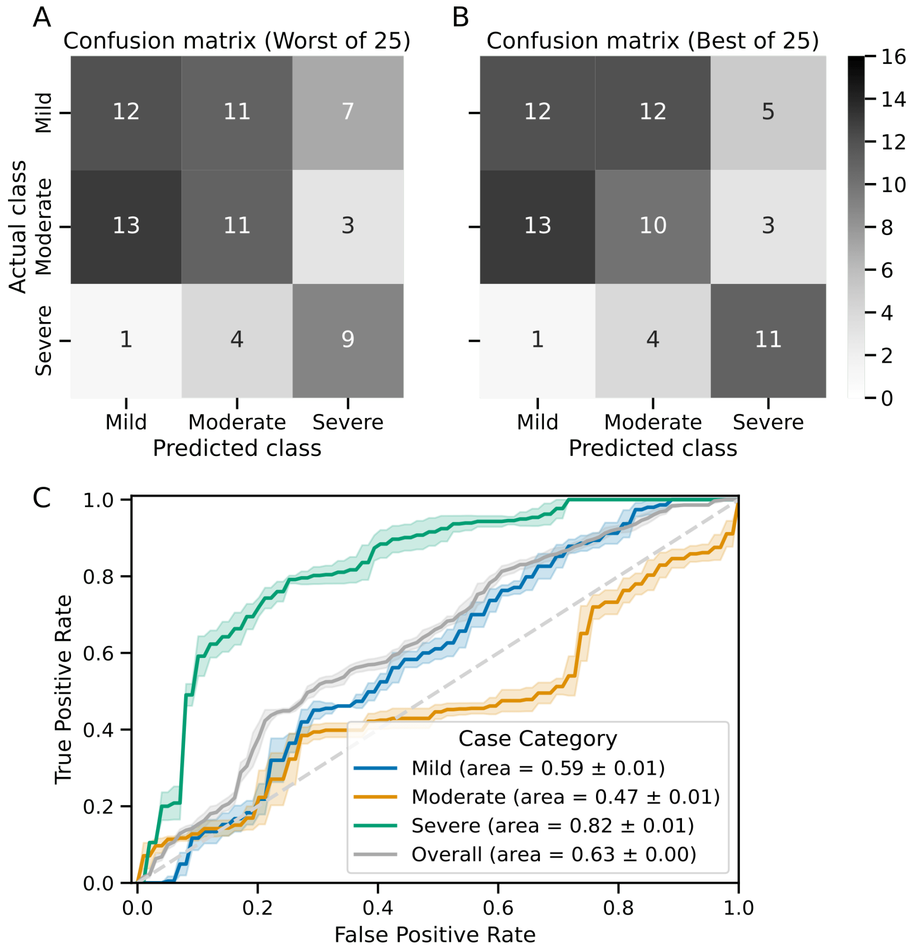


Figure S57: Disease-severity prediction using GBM classifiers trained on fractions of five T cell types of the covid-blood-severity-hq data. The five T cell types (CD8_EM, CD8_Tc2, TFH, TH17_cluster1, Treg_active) were selected using forward feature selection of the reconstructed covid-blood-severity-hq data. Confidence intervals were calculated using 25 runs of LOOCV. The disease category "critical" was combined with "severe" and "asymptomatic" with "mild". **A:** Confusion matrix for the worst run of LOOCV. **B:** Confusion matrix for the best run of LOOCV. **C:** ROC curve for the prediction of mild - blue, moderate - yellow, severe - green, and all categories (overall) - gray. Confidence intervals indicate one standard deviation.

Table S1: *Overview of all single cell and bulk sequencing datasets used in this study.* The table shows the dataset name, size of the dataset, the sequencing technology, cell types as annotated in the original study and a hyperlink to the publication.

| ***Dataset*** | ***Method*** | ***Cell Types*** | ***Publication or Download link*** |
|---|---|---|---|
| **pancreas** (8569 cells) | SMARTSeq2, Fluidigm C1, CelSeq, CelSeq2, inDrops | alpha, beta, ductal, acinar, delta, gamma, activated_stellate, endothelial, quiescent_stellate, macrophage, mast, epsilon, schwann | [34] |
| **difftec** (31 021 cells) | 10x Chromium v2, 10x Chromium v3, SMARTSeq2, Seq-Well, inDrops, Drop-seq, CelSeq2 | Cytotoxic T cell, $CD4^+$ T cell, $CD14^+$ monocyte, B cell, Natural killer cell, Megakaryocyte, $CD16^+$ monocyte, Dendritic cell, Plasmacytoid dendritic cell, Unassigned | [35] |
| **snRNA-seq & scRNA-seq** (12 423 cells) | snRNA-seq and scRNA-seq using Chromium single-cell 3' v3 | Epithelial cells, Macrophages, Hepatocytes, T cells, Endothelial cell, Fibroblasts, B cells, NK cells | `https://www.ncbi.nlm.nih.gov/geo/query/acc.cgi?acc=GSM4186980` `https://www.ncbi.nlm.nih.gov/geo/query/acc.cgi?acc=GSM4186974` |

Table S1: *Overview of all single cell and bulk sequencing datasets used in this study continued.*

| ***Dataset*** | ***Method*** | ***Cell Types*** | ***Publication or Download link*** |
|---|---|---|---|
| **covid-lung** (56 645 cells) | 10X Genomics Chromium Single Cell 5'v1.1 | CD8 T, TREG, CD4_CD8 proliferating, B cell, CD4_TCM, TRM1, TR1, CD8_TCM, T senescent, CD8_TEM, TEM17, T antiviral, alveolar MΦ, TRM17, M1, CD4_CD8 stressed TCM, CD4_TSCM, MAIT, Innate like, Neutrophils, doublets, CD4_CD8 lnc rich, aged Neutrophils, M1 $HSP^{+}$, Mast, DC, M1 Mono-derived, M2 profibrotic, Epithelial, Neutrophil, Macrophage | [36] |
| **covid-blood** (83 709 cells) | 10X Genomics Chromium Single Cell 5'v1.1 | $CD3^{+}$ cells | [36] |

Table S1: *Overview of all single cell and bulk sequencing datasets used in this study continued.*

| ***Dataset*** | ***Method*** | ***Cell Types*** | ***Publication or Download link*** |
|---|---|---|---|
| **citeseq** (6592 cells) | 10x Genomics Single Cell and CITE-seq | B cells, CD4 T cells, NK cells, $CD14^+$ Monocytes, $FCGR3A^+$ Monocytes, CD8 T cells | `https://www.ncbi.nlm.nih.gov/geo/query/acc.cgi?acc=GSE100866` `https://github.com/YosefLab/scVI-data/raw/master/pbmc_metadata.pickle` |
| **bulk** (9852 cells) | SMART-seq v4 | Naive CD4, Memory CD4, TH1, TH2, TH17, Tfh, Fr. I nTreg, Fr. II eTreg, Fr. III T, Naive CD8, Memory CD8, CM CD8, EM CD8, TEMRA CD8, NK, Naive B, USM B, SM B, Plasmablast, DN B, CL Monocytes, Int Monocytes, NC Monocytes, mDC, pDC, Neutrophils, LDG | [37] |

Table S1: *Overview of all single cell and bulk sequencing datasets used in this study continued.*

| ***Dataset*** | ***Method*** | ***Cell Types*** | ***Publication or Download link*** |
|---|---|---|---|
| **covid-blood-severity** (636 836 cells) | 10X Genomics Chromium Single Cell 5'v1.1 | ASDC, B_exhausted, B_immature, B_malignant, B_naive, B_non-switched_memory, B_switched_memory, C1_CD16_mono, CD4.CM, CD4.EM, CD4.IL22, CD4.Naive, CD4.Prolif, CD4.Tfh, CD4.Th1, CD4.Th2', CD4.Th17, CD8.EM, CD8.Naive, CD8.Prolif, CD8.TE, CD14_mono, CD16_mono, CD83_CD14_mono, DC1, DC2, DC3, DC_prolif, HSC_CD38neg, HSC_CD38pos, HSC_MK, HSC_erythroid, HSC_myeloid, HSC_prolif, ILC1_3, ILC2, MAIT, Mono_prolif, NKT, NK_16hi, NK_56hi, NK_prolif, Plasma_cell_IgA, Plasma_cell_IgG, Plasma_cell_IgM, Plasmablast, Platelets, RBC, Treg, gdT, pDC | [38] |

Table S1: *Overview of all single cell and bulk sequencing datasets used in this study continued.*

| ***Dataset*** | ***Method*** | ***Cell Types*** | ***Publication or Download link*** |
|---|---|---|---|
| **Kidney snRNA-seq & scRNA-seq** (82 701 cells) | 10x Genomics Chromium | None (not annotated) | `https://atlas.kpmp.org/repository/?facetTab=participants` Patients: 3010018, 3010034, 3210003, 3210034, 3310005, 3310006, 3410050, 3410184, 3410187 |

Table S2: *Detailed quality and batch information for all single cell and bulk sequencing datasets used in this study.* For each batch, the number of cells, the mean number of counts per cell, and the mean number of expressed genes per cell are listed. For the difftec dataset, the batch names were slightly adjusted. Their published batch names are written in brackets.

| **Dataset** | **Batch** | **Number of cells** | **Mean number of counts per cell** | **Mean number of genes** |
|---|---|---|---|---|
| | smartseq2 | 2394 | 451021.4 | 6214.0 |
| | fluidigmc1 | 638 | 1580155.4 | 8127.4 |
| **pancreas** | celseq | 2285 | 11161.1 | 3466.8 |

Table S2: *Detailed quality and batch information for all single cell and bulk sequencing datasets used in this study continued.*

| **Dataset** | **Batch** | **Number of cells** | **Mean number of counts per cell** | **Mean number of genes** |
|---|---|---|---|---|
| **difftec** | celseq2 | 1004 | 23394.2 | 5274.9 |
| | indrop | 8569 | 5828.2 | 1887.2 |
| | dropseq (pbmc1_Drop-seq) | 3222 | 1282.0 | 676.0 |
| | indrops (pbmc1_inDrops) | 3222 | 566.3 | 362.4 |
| | seqwell (pmbc1_Seq-Well) | 3222 | 1035.3 | 567.2 |
| | chromium-v3 (pbmc1_10x Chromium (v3)) | 3222 | 4891.3 | 1514.1 |
| | chromium-v2 (pbmc1_10x Chromium (v2) A) | 3222 | 2120.0 | 795.4 |
| | chromium-v2B (pmbc1_10x Chromium (v2) B) | 3222 | 2512.4 | 870.8 |
| | smartseq2 (pbmc1_Smart-seq2) | 253 | 385914.3 | 2434.6 |
| | celseq2 (pbmc1_CEL-Seq2) | 253 | 6057.3 | 2585.4 |
| | dropseq-2 (pbmc2_Drop-seq) | 3362 | 2141.0 | 977.7 |
| | seqwell-2 (pbmc2_Seq-Well) | 551 | 692.6 | 421.8 |

Table S2: *Detailed quality and batch information for all single cell and bulk sequencing datasets used in this study continued.*

| **Dataset** | **Batch** | **Number of cells** | **Mean number of counts per cell** | **Mean number of genes** |
|---|---|---|---|---|
| | smartseq2-2 (pbmc2_Smart-seq2) | 273 | 292924.3 | 2795.4 |
| | celseq2-2 (pbmc2_CEL-Seq2) | 273 | 5949.3 | 2556.6 |
| | chromium-v2-2 (pbmc2_10x Chromium (v2)) | 3362 | 2860.7 | 1131.4 |
| | indrops-2 (pbmc2_inDrops) | 3362 | 1249.5 | 619.5 |
| **snRNA-seq & scRNA-seq** | sn-lq | 7260 | 2206.6 | 1308.7 |
| | sc-hq | 5163 | 4634.5 | 1214.6 |
| **covid-lung** | Bacterial | 14591 | 9627.2 | 1617.4 |
| | SARS-CoV-2 | 42054 | 10284.4 | 1719.5 |
| **covid-blood** | Bacterial | 22199 | 5861.6 | 1703.0 |
| | SARS-CoV-2 | 61510 | 5388.6 | 1700.7 |
| **citeseq** | citeseq | 6592 | 1391.8 | 797.8 |
| **bulk** | bulk | 9852 | 881440.6 | 13103.8 |
| **covid-blood-severity** | cambridge | 130637 | 4798.9 | 1485.9 |
| | ncl | 431733 | 3520.0 | 1276.1 |
| | sanger | 74466 | 3640.2 | 1445.1 |
| **Kidney snRNA-seq & scRNA-seq** | kidney-lq (snRNA-seq) | 52934 | 6532.8 | 2462.7 |
| | kidney-hq (snRNA-seq) | 29767 | 4449.6 | 1546.0 |

Table S3: Antibodies used in the CITE-seq experiments of the citeseq dataset (see table S1 & table S2).

| Antibody | Clone | Supplier | Target Protein | Target Gene |
|---|---|---|---|---|
| CD3e | UCHT1 | BioLegend, USA | CD3 | *CD3E, CD3D* |
| CD19 | HIB19 | BioLegend, USA | CD19 | *CD19* |
| CD4 | RPA-T4 | BioLegend, USA | CD4 | *CD4* |
| CD8a | RPA-T8 | BioLegend, USA | CD8 | *CD8A* |
| CD56 | MEM-188 | BioLegend, USA | NCAM1 | *NCAM1* |
| CD16 | B73.1 | BioLegend, USA | FCG3A | *FCGR3A* |
| CD11c | B-ly6 | BD Pharmingen, USA | CD11c | *ITGAX* |
| CCR7 | 150603 | RD Systems, USA | CCR7 | *CCR7* |
| CCR5 | J418F1 | BioLegend, USA | CCR5 | *CCR5* |
| CD34 | 581 | BioLegend, USA | CD34 | *CD34* |
| CD14 | M5E2 | BioLegend, USA | CD14 | *CD14* |
| CD10 | HI10a | BioLegend, USA | NEP | *MME, CD10* |
| CD45RA | HI100 | BioLegend, USA | PTPRC, CD45RA | *PTPRC* |
| CD2 | RPA-2.10 | BioLegend, USA | CD2 | *CD2* |
| CD57 | H-NK1 | BioLegend, USA | B3GA1, CD57 | *B3GAT1* |

Table S4: *Disease-severity prediction performance using GBM classifiers trained on T cell fractions.* Column one (3 - classes) and column two (2 - classes) displays the classification performance using cell type fractions obtained with reconstructed covid-blood-severity-hq data for three classes (mild, moderate, and severe) and two classes (mild and severe), respectively. The third column (2 - classes published) shows the classification performance for the fractions based on the originally published T cell annotations. All classifications were conducted with a GBM using 25 runs of LOOCV (confidence intervals) and forward feature selection. The T cell subtypes used by the GBM for column one are CD8_EM, CD8_Tc2, TFH, TH17_cluster1, Treg_active. For column two the features are CD4_CM, CD4_cytotoxic, CD4_naive, CD8_EM, CD8_effector. For column three CD4.CM, CD4.Tfh, CD8.EM, NKT, and Treg cells were used. It is striking to observe the strong increase in performance in the 2 class case between reconstructed cell type (column 2) and originally published (column 3) cell type information.

| | **3 - classes** (mild, moderate, severe) | **2 - classes** (mild, severe) | **2 - classes published** (mild, severe) |
|---|---|---|---|
| **F1-Score (Micro)** | $0.46 \pm 0.01$ | $\mathbf{0.82 \pm 0.01}$ | $0.61 \pm 0.01$ |

| | | | |
|---|---|---|---|
| **F1-Score (Macro)** | $0.47 \pm 0.01$ | $\mathbf{0.82 \pm 0.01}$ | $0.58 \pm 0.01$ |
| **AUROC** | $0.63 \pm 0.00$ | $\mathbf{0.81 \pm 0.00}$ | $0.55 \pm 0.01$ |
| **Accuracy** | $0.46 \pm 0.01$ | $\mathbf{0.82 \pm 0.01}$ | $0.61 \pm 0.01$ |
